# Age-associated increased stiffness of the ovarian microenvironment impairs follicle development and oocyte quality and rapidly alters follicle gene expression

**DOI:** 10.1101/2024.06.09.598134

**Authors:** Sara Pietroforte, Makenzie Plough, Farners Amargant

## Abstract

In humans, aging triggers cellular and tissue deterioration, and the female reproductive system is the first to show signs of decline. Reproductive aging is associated with decreased ovarian reserve, decreased quality of the remaining oocytes, and decreased production of the ovarian hormones estrogen and progesterone. With aging, both mouse and human ovaries become pro-fibrotic and stiff. However, whether stiffness directly impairs ovarian function, folliculogenesis, and oocyte quality is unknown. To answer this question, we cultured mouse follicles in alginate gels that mimicked the stiffness of reproductively young and old ovaries. Follicles cultured in stiff hydrogels exhibited decreased survival and growth, decreased granulosa cell viability and estradiol synthesis, and decreased oocyte quality. We also observed a reduction in the number of granulosa cell-oocyte transzonal projections. RNA sequencing revealed early changes in the follicle transcriptome in response to stiffness. Follicles cultured in a stiff environment had lower expression of genes related to follicle development and greater expression of genes related to inflammation and extracellular matrix remodeling than follicles cultured in a soft environment. Altogether, our findings suggest that ovarian stiffness directly modulates folliculogenesis and contributes to the progressive decline in oocyte quantity and quality observed in women of advanced maternal age.

## Introduction

The female reproductive system shows signs of age-associated cellular and tissue deterioration before any other organ in the human body [1, 2]. Reproductive aging is characterized by a decline in oocyte quantity and quality, as well as a significant reduction in the ovarian-produced hormones estrogen and progesterone [1]. Female reproductive aging both causes infertility and affects women’s health, as estrogen and progesterone regulate many tissues such as bones and the liver [3–5]. We and others reported that mouse and human ovaries become stiffer with advanced maternal age and that this increased stiffness is triggered by increased collagen deposition (fibrosis) and decreased abundance of the extracellular matrix (ECM) glycosaminoglycan hyaluronan [6, 7].

The physical properties of the tissue have emerged as an important regulator of cell and tissue function because cells detect mechanical stiffness by a variety of receptors, including integrins, and trigger complex signaling pathways that result in cell proliferation, cell death, ECM production, or inflammation, key processes in development and tissue homeostasis [8, 9]. The ovarian follicle, which comprise an oocyte surrounded by supporting granulosa cells, is a unique system to study biomechanics and mechanobiology [10]. Follicles are exposed to the surrounding ovarian microenvironment for decades. This is because follicles form before birth and can remain in a quiescent state for decades before they become activated to grow and ovulate. In fact, we know that mechanical stress regulates primordial follicle dormancy in mice [11].

However, the age-associated increase in ovarian stiffness is also likely to contribute to the decrease in follicle number and oocyte quality observed with advanced maternal age for several reasons. First, multiple studies have reported a correlation between increased tissue stiffness and pathologies such as cardiovascular disease and diabetes [12, 13]. Second, mouse follicles grew more rapidly when cultured in a low percentage 3D alginate hydrogel than when cultured in a high percentage alginate hydrogel [14–17]. Finally, treatment of reproductively old mice with anti-fibrotic drugs to reduce fibrosis, and therefore stiffness, restored ovulation and increased follicle development [18] *(Manuscript in preparation)*. However, how mechanical stiffness regulates folliculogenesis and oocyte quality, and how these signals are integrated by the follicle, is largely unknown.

Here, we tested the hypothesis that the age-associated increase in ovarian stiffness triggers the decline in follicle growth and oocyte quality observed with advanced maternal age. To do so, we used a 3D *in vitro* follicle growth system in which mouse secondary follicles are isolated from the ovary and cultured in alginate hydrogels [19, 20]. We generated alginate hydrogels that recapitulate the stiffness of reproductively old mouse ovaries. Follicles cultured in stiff hydrogels had slower growth, less survival, less estradiol production, more granulosa cell death, and lower oocyte quality than those cultured in non-stiff hydrogels. RNA sequencing (RNA-seq) demonstrated that follicles responded to biomechanical cues within three hours. Follicles cultured in stiff environment showed reduced expression of genes related to follicle development and increased expression of genes related to inflammation and ECM remodeling. Overall, we have developed a physiologically relevant system to study the impact of ovarian stiffness on folliculogenesis. Importantly, all the changes we report are observed in ovaries from reproductively old mice and women [1], suggesting that stiffness contribute to the decline in oocyte quantity and quality observed in women of advanced reproductive age.

## Materials and Methods

### Animals

Pre-pubertal (12-13 days) CB6F1 and CD1 mice were used to maximize the yield of secondary follicles. CB6F1 hybrids were generated by housing two Balb/c females with one C57BL/6 male per cage. CD-1 pre-pubertal mice were obtained together with a dam. All mice were purchased from Envigo (Indianapolis, IN, USA). Ovaries from 4-month-old CB6F1 female mice were used for the nanoindentation control. Mice were housed at Northwestern University’s Center for Comparative Medicine or at Washington University’s Division of Comparative Medicine under constant temperature, humidity, and light (12 hours light/12 hours dark) with free access to food and water. All mouse procedures were reviewed and approved by the Institutional Animal Care and Use Committee of Northwestern University (Chicago, IL, US) or Washington University in St. Louis (St. Louis, MO, US) and were performed in accordance with National Institutes of Health guidelines.

### Alginate gel preparation

To prepare alginate solutions, distilled phosphate buffered saline (DPBS) without calcium and magnesium (14190250, ThermoFisher Scientific, Waltham, MA, US) was filtered (0.2 μm) and mixed with Alginic acid solution salt from brown algae (71238, Sigma-Aldrich, St. Louis, MO, US) to create final alginate solutions of 0.5%, 2%, and 4% weight/volume (w/v). The alginate solutions were placed on a rocker at 4 °C for 24 hours, then stored at 4 °C and used within one month. To prepare alginate gels, a 7 μl drop of alginate solution was placed in a calcium solution (50 mM CaCl_2_ and 140 mM NaCl_2_) to crosslink the gel for 3 minutes, then washed at least three times with DPBS [20]. To create flat alginate gels for nanoindentation, 1 ml of alginate solution was placed in a 24 mm round dish, calcium solution was added for 3 minutes, then gels were washed at least three times with DPBS.

### Nanoindentation

CB6F1 mouse ovaries were removed from the bursa and placed in L15 medium supplemented with 3 mg/ml polyvinylpyrrolidone (PVP, P0930, Sigma-Aldrich) and 0.5% Pen-Strep. Each ovary was then attached to a plastic petri dish with tissue glue (PELCO® Pro CA44 Tissue Adhesive, Fresno, CA, US) and covered with warm 1x DPBS with calcium and magnesium (14040182, ThermoFisher Scientific). Flat 0.5% and 2% alginate gels were covered with warm 1x DPBS. Ovaries and gels were indented with a Piuma nanoindenter (Optics 11). The probe had a glass spherical tip of 100 μm in diameter in a cantilever with a spring constant of 0.5 N m-1. For all measurements, the indentation depth was set at a maximum of 10% of the probe diameter. A minimum of 10 indentations were performed in each ovary or gel. To obtain the elastic moduli (E), the data were analyzed by using the Hertzian contact model in the Piuma software.

### Diffusion test

Drops of alginate were crosslinked with calcium solution for 3 minutes as described above to produce beads, which were washed three times in PBS and incubated with a solution of Alcian Blue for 3 and 5 minutes. Then, beads were rinsed in PBS three times, and the blue staining was visually evaluated. Beads were also bisected to determine whether the Alcian blue dye reached the center of the bead.

### Follicle isolation, encapsulation, and growth

Follicles were cultured and analyzed as previously described [20]. Briefly, ovaries from day 12 or 13 prepubertal CB6F1 and CD1 mice were placed in dissection media containing L15, 1% fetal bovine serum (FBS), and 0.5% penicillin-streptomycin (Pen-Strep) (Gibco, ThermoFisher). Early secondary follicles (120-130 μm) were mechanically isolated with insulin syringes. Follicles were incubated for 1 hour at 37 °C in a-MEM-Glutamax containing 1% FBS and 0.5% Pen-Strep in a humidified atmosphere of 5% CO_2_. Only follicles with an intact basement membrane and visible and healthy oocytes surrounded by two layers of granulosa cells were selected for culture. To encapsulate the follicles, the parafilm technique was performed [20]. Groups of 10 follicles were mixed with 7 μl of 0.5%, 2%, or 4% w/v alginate solution. The alginate drops were crosslinked as described above. The resulting alginate beads were then incubated in α-MEM-Glutamax containing 1% FBS and 0.5% Pen-Strep for 1 hour. Finally, alginate beads were transferred to a 96-well ultra-low attachment plate (Corning, Corning, NY, US) with one bead per well. Each well contained 100 μl of growth media (α-MEM-Glutamax containing 3 mg/ml bovine serum albumin (MP Biomedicals), 1 mg/ml fetuin (Sigma-Aldrich), 0.1% insulin-transferrin-selenium (ThermoFisher Scientific), and 10 mIU/mL follicle stimulating hormone (Gonal-F, EMD Serono, Inc. Rockland, MA, US)). Follicles were cultured for up to 12 days at 37 °C, 5% CO_2_. Every other day, half of the media was removed, frozen for downstream analysis, and replaced with fresh media.

### Follicle survival, growth, and granular pattern

Follicles were imaged every other day by using 4x, 10x, and 20x objectives on an Evos FL Auto imaging system (ThermoFisher Scientific). Individual follicles were tracked by analyzing their relative position inside the bead in the 4x objective images. Follicle size and survival were measured in Fiji (ImageJ, National Institute of Health, Bethesda, MD, US). Follicle size measurements were only performed until day 10 because follicles close to one another tended to fuse by day 12 of culture. Follicle growth was analyzed by calculating the average size of two perpendicular diameter measurements for each follicle. Follicles that did not grow for 2 to 4 consecutive days were considered dead. Follicle survival and growth values were plotted in Prism (GraphPad). The presence or absence of a granular appearance of the follicles was assessed in 20x objective images.

### Live/dead assay

Follicles cultured in 0.5% and 2% alginate beads were isolated from alginate gels as described above at 2, 4, 6, 8, and 10 days. They were then rinsed three times in dissection media, incubated for 30 minutes in a live/dead Viability/Cytotoxicity solution containing 4 mM Calcein and 8 mM Ethidium homodimer-1 (L3224, ThermoFisher Scientific) at 37 °C, 5% CO_2_, washed three times in DPBS, and imaged at 4x, 10x, and 20x magnification with an EVOS system (Calcein 494/517nm, Ethidium homodimer-1 528/617nm).

### Estradiol measurement

The concentration of 17β-estradiol in the frozen conditioned media was measured by using ELISA kits (Cayman Chemicals, Ann Arbor, MI, US) according to the manufacturer’s instructions. Media samples collected from wells without follicles were used as negative controls. Media was diluted 1:10 in assay buffer to ensure the readings were within the detection limits. Concentrations are reported in ng/ml.

### Staining trans-zonal projections (TZPs)

Follicles were fixed with 3.8% paraformaldehyde in PBS containing 1% Triton for 20 min at 37 °C, washed three times in PBS-PVP, and stored at 4 °C until processed for immunofluorescence. Follicles were permeabilized in PBS containing 1% BSA and 0.1% Triton X-100 for 15 min at RT, washed three times 5 min in blocking buffer (PBS with 1% BSA and 0.1% Tween-20), then incubated with Phalloidin 633 (A22284, ThermoFisher Scientific) for 1 h at RT with gentle rocking. Then, follicles were washed three times 20 min with blocking buffer, then mounted onto microscope slides with Vectashield Plus Antifade Mounting Medium containing 4’,6’-diamidino-2-phenylindole (DAPI, Vector Laboratories). Slides were imaged with a Leica TCS Sp5 confocal system with 40x or 63x objectives (Leica Microsystems, Wetzlar, Germany). In Fiji software, a background threshold was determined in each follicle by measuring the intensity in a region without actin staining. Then, a 10 μm line was drawn between the granulosa cells and the oocyte interface (Zona Pellucida), and a phalloidin intensity profile was generated. The number of TZPs was determined by counting the number of peaks above the background threshold.

### Oocyte analysis

Follicles were removed from the alginate on day 12 by incubating the beads in L15 media containing 1 mg/ml of alginate lyase for 20 min at RT. The follicles were then washed three times in L15 media. Oocytes were mechanically isolated from the follicle and imaged with the EVOS system. Oocytes with a visible germinal vesicle (GV) and without fragmentation were considered to be good quality.

### RNA sequencing and data analysis

Follicles were isolated from pre-pubertal (12 – 13 days) CB6F1 mice and cultured in 0.5% or 2% alginate beads as described above but in groups of 5 follicles per bead. Follicles were isolated at 3h, 6h, 12h, and 24h and incubated for 5 min in 4 mg/ml alginate lyase. Isolated follicles were snap-frozen in a minimum volume of PBS and submitted to the Northwestern University NUSeq Core for processing and RNA sequencing in parallel. The SMART-seq mRNA kit (Takara, San Jose, California, USA) was used to construct cDNA, and libraries were sequenced as 50 bp single-end reads on a Hiseq 4000 sequencer (Illumina, San Diego, California, USA), with about 90 million reads in total, to produce the raw *fastq* files. Raw data were obtained for 160 follicles divided in two groups (0.5% and 2%) at four time points (3h, 6h, 12h, and 24h) with four replicates of five follicles per condition (Supplementary Table S1). FastQC (Babraham Bioinformatics, Cambridge, UK) was used to assess quality of the sequencing data.

Reads were aligned to the UCSC mm10 genome (http://genome.ucsc.edu) by using STAR 2.7.0a with gene models from Ensembl (http://www.ensembl.org) and counted by using the function SummarizeOverlap of the Bioconductor/R package Genomic Features [21]. DESeq2 [22] was then used to detect differentially expressed genes (DEGs) between experimental groups, and genes with adjusted p-value <0.05 were considered statistically significant. The threshold of log2 Fold Change (log2FC) = |2| was considered to calculate up- and down-regulated genes. Volcano plots of the statistically significant genes for each comparison were created by using the ‘plot’ function in R (log2FC and -log10(padj)).

To analyze genes with stable expression across conditions and replicates, DESeq2 was used to select genes with low log2FC and high base mean values (log2FC = 0.5 and base mean = 100). ShilyGO 0.80 [23] was used for pathway analysis of these stably expressed genes.

Gene set enrichment analysis (GSEA) was performed by WebGestalt [24], using the gene ontology (GO) database for biological function and selecting the standard parameters for analysis (Organism: *mus musculus*; Enrichment Categories: *geneontology Biological Process noRedundant*; ID type: *ensemble gene ID*; Minimum number of ID in the category: *3*; Significance level: *top 20*). Up- and down-regulated genes were specifically analyzed individually in WebGestalt by using Over-Representation Analysis and the GO database for biological function, selecting the same above-mentioned standard parameters. Additional GSEA of pathways of interest was performed with GSEA Software 4.3.3 [25, 26] with *Mouse MSigDB v2023.2.Mm* (October 2023) for GO biological process and/or Reactome pathways provided by the software.

### Statistics

The normality of the data was analyzed by using the Shapiro-Wilk test, and normally distributed data were analyzed by using Student t-test for comparisons between two groups or one-way ANOVA for more than two groups. The statistical package in Prism 9.4 (GraphPad Software) was used for all analyses.

## Results

### A stiff microenvironment mimicking that of reproductively old ovaries impairs follicle survival and growth

Mouse ovaries become stiffer with advanced reproductive age (6–12 weeks: 1.98±0.42 kPa, 14– 17 months: 4.36± 1.24 kPa) [6]. To examine the effect of the age-associated increased ovarian stiffness on follicle development, we used a system in which follicles are cultured *in vitro* in alginate hydrogel beads. We generated 0.5% alginate, which is commonly used to support growth of mouse follicles to the antral stage [14, 15, 19, 20], and 2% alginate, which is stiffer. We first used instrumental indentation to measure the stiffness of 0.5% and 2% alginate beads. In control experiments, ovaries from 4-month-old mice had a stiffness of 2.00±0.17 kPa, confirming that the indentation procedure yielded values in the expected range. The 0.5% alginate had a stiffness of 0.84±0.16 kPa, whereas the 2% alginate was 7-fold stiffer (5.85±0.28 kPa, p < 0.05), similar to the stiffness in a reproductively old mouse ovary (Figure 1A). These stiffness values are in line with reported rheometry data [15] and indicate that the 0.5% alginate mimics the extremely soft and permissive environment of a reproductively young ovary, and the 2% alginate mimics the stiff environment of an old mouse ovary.

**Figure 1:**
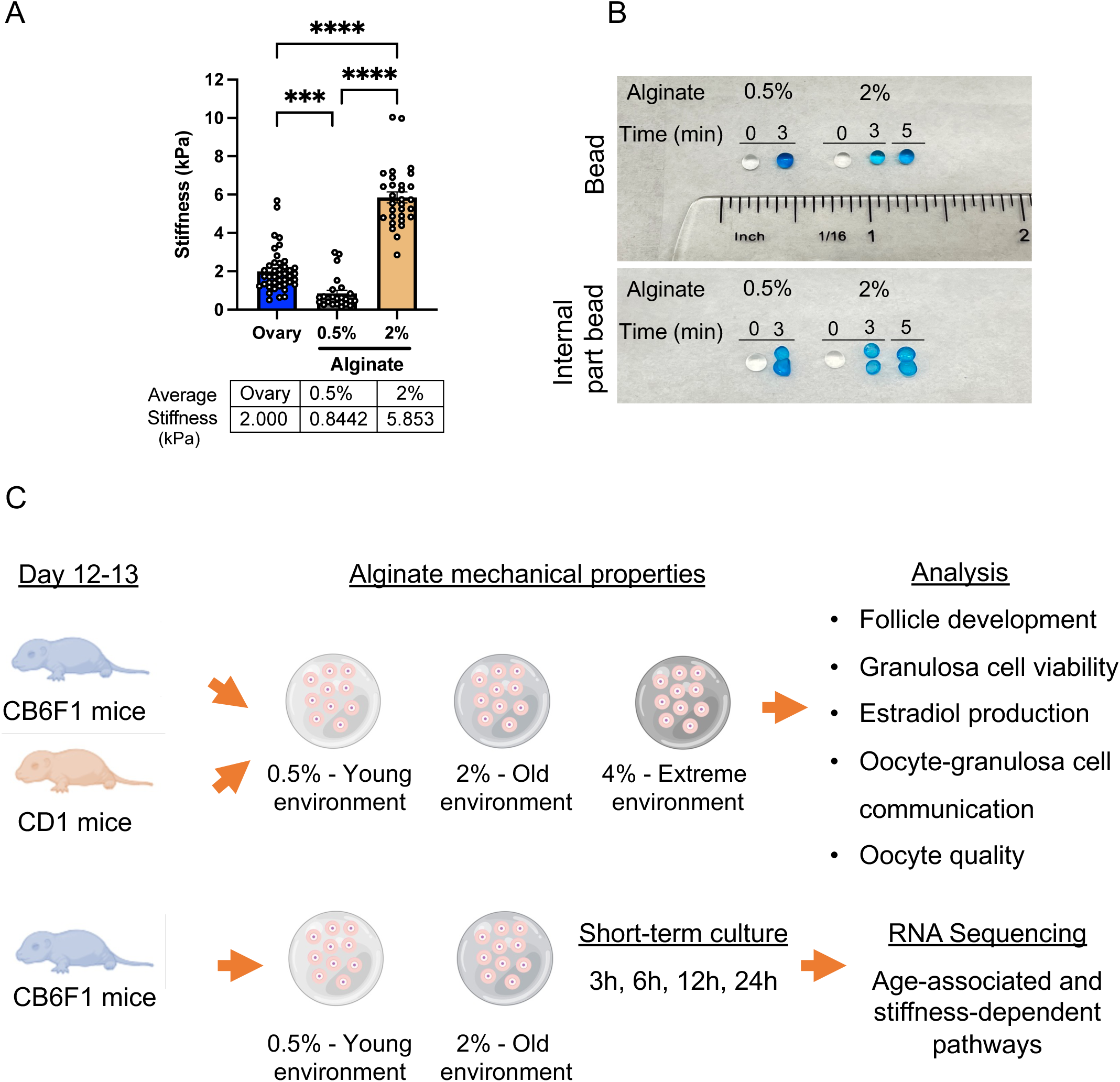
Alginate hydrogels recapitulate the stiffness of reproductively young and old mouse ovaries. **(A)** Quantification of the stiffness of whole ovaries from reproductively young CB6F1 mice and 0.5% and 2% w/v alginate gels. **(B)** Representative images of 0.5% and 2% w/v alginate gels before and after staining with Alcian blue for the indicated times. Top image, whole beads; bottom image, bisected beads. *** p<0.001, ****p<0.0001. **(C)** Schematic representation of the *ex vivo in vitro* follicle growth system used to examine the impact of the aging microenvironment stiffnesses on follicle development and oocyte quality.

To assess whether 2% alginate would prevent nutrients in the media from reaching the follicles, we generated 0.5% and 2% alginate beads and incubated them in the dye Alcian blue, which has a molecular weight of 1298.86 g/mol. After 3 minutes, both 0.5% and 2% alginate beads were equally blue, and the blue intensity did not increase between 3 and 5 minutes of incubation in the dye. We then bisected the beads and confirmed that the dye diffused across the entire bead (Figure 1B). Overall, these results show that this system allows us to examine the impact of ovarian stiffness on follicle development.

To explore whether increased stiffness of the microenvironment affects follicle survival and growth, we cultured early secondary follicles in 0.5% and 2% alginate for 12 days. We also included follicles cultured in 4% alginate as an extremely stiff environment condition (Figure 1C). Given that mouse strain-dependent differences in early follicle growth patterns were reported [27], we performed these experiments with follicles from both the hybrid CB6F1 mouse strain and the outbred CD1 strain (Figure 1C). On day 12 of culture, 100% of CB6F1 follicles and over 80% of CD1 follicles cultured in 0.5% alginate were viable (Figure 2 A-D). In contrast, only 6.66% of CB6F1 and 30% of CD1 follicles cultured in 2% alginate were viable, and none of the follicles cultured in 4% alginate were viable (Figure 2 A-D). These results indicate that follicles are highly sensitive to changes in the mechanical properties of the microenvironment.

**Figure 2:**
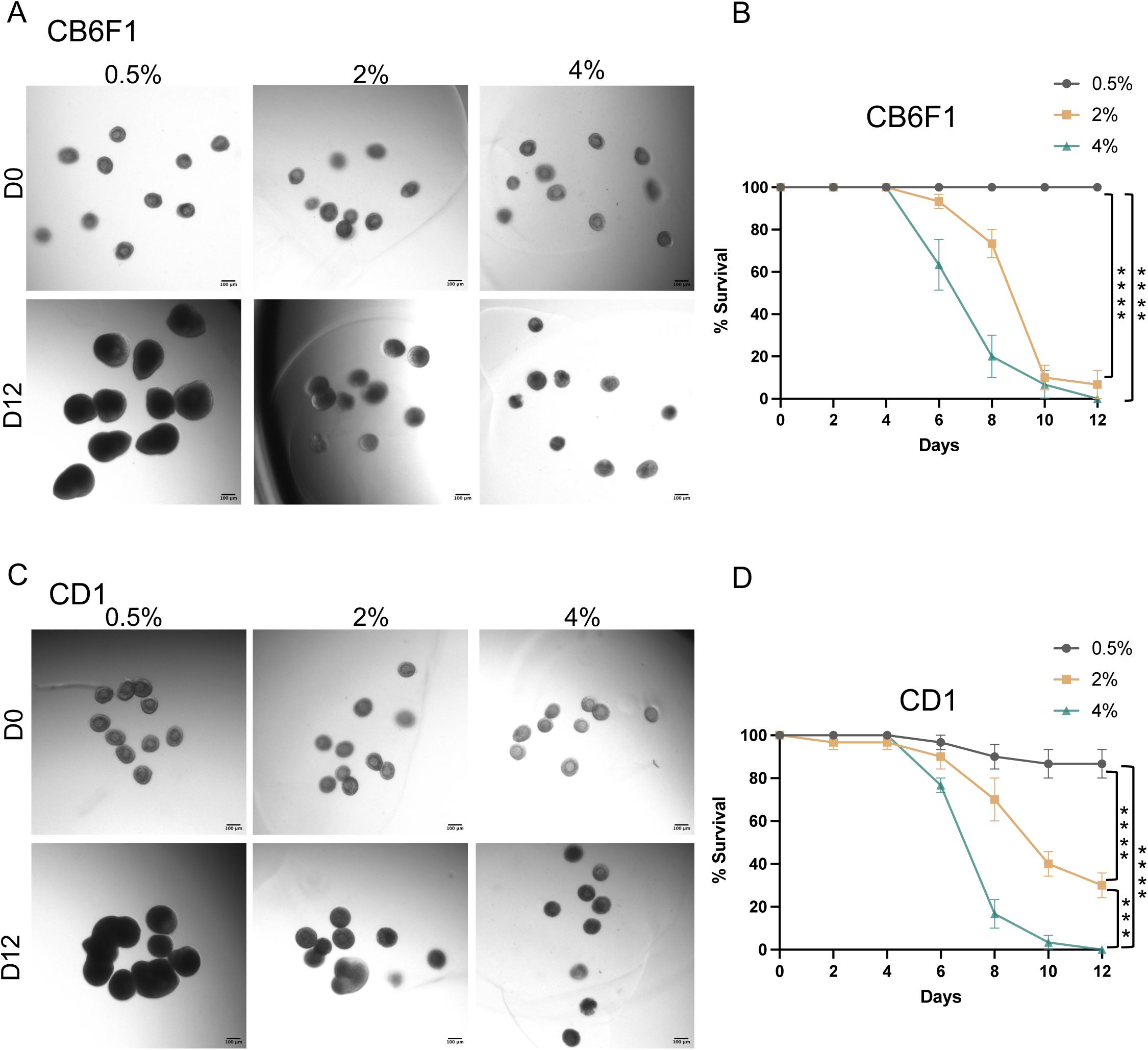
Age-associated increased ovarian stiffness impairs follicle viability *in vitro*. **(A)** Representative transmitted light images of CB6F1follicles cultured in 0.5%, 2%, and 4% alginate beads at day 0 (D0) and day 12 (D12). **(B)** Graphs showing the CB6F1 follicle survival percentage along the culture and comparing D0 and D12. **(C)** Representative transmitted light images are shown of CD1 follicles that were either cultured in 0.5%, 2% or 4% alginate gels. **(D)** Quantification of CD1 follicle survival during the 12 days of culture. N=3 independent experiments (N=30 follicles per condition). *** p<0.001, ****p<0.0001.

We next analyzed the growth rates of the follicles that survived during the 12-day culture period. However, we only analyzed follicle growth until day 10 because follicles tended to fuse by day 12. On day 0, follicles from CB6F1 mice were 122.0 ± 2.48 µm in diameter and those from CD1 mice were 125.3 ± 1.88 µm. Follicles cultured in 0.5% alginate showed significant growth by day 6 (CB6F1, 171.6±13.58 μm; CD1 200.4±23.01 μm; Figure 3 A, D, E, H) and reached 247.6±13.76 μm (CB6F1) and 226.9±8.69 μm (CD1) by day 10 of culture. In 2% and 4% alginate, follicles stopped growing by day 6. Follicles from CB6F1 mice reached a final size of 157.5±10.33 μm in 2% alginate and 136.1±2.77 μm in 4% alginate. Follicles from CD1 reached a final size of 160.8±4.03 μm in 2% alginate and 129.4±2.57μm in 4% alginate (Figure 3 A, D, E, H). These results indicate that stiff microenvironments slow follicle growth during the active expansion phase of folliculogenesis.

**Figure 3:**
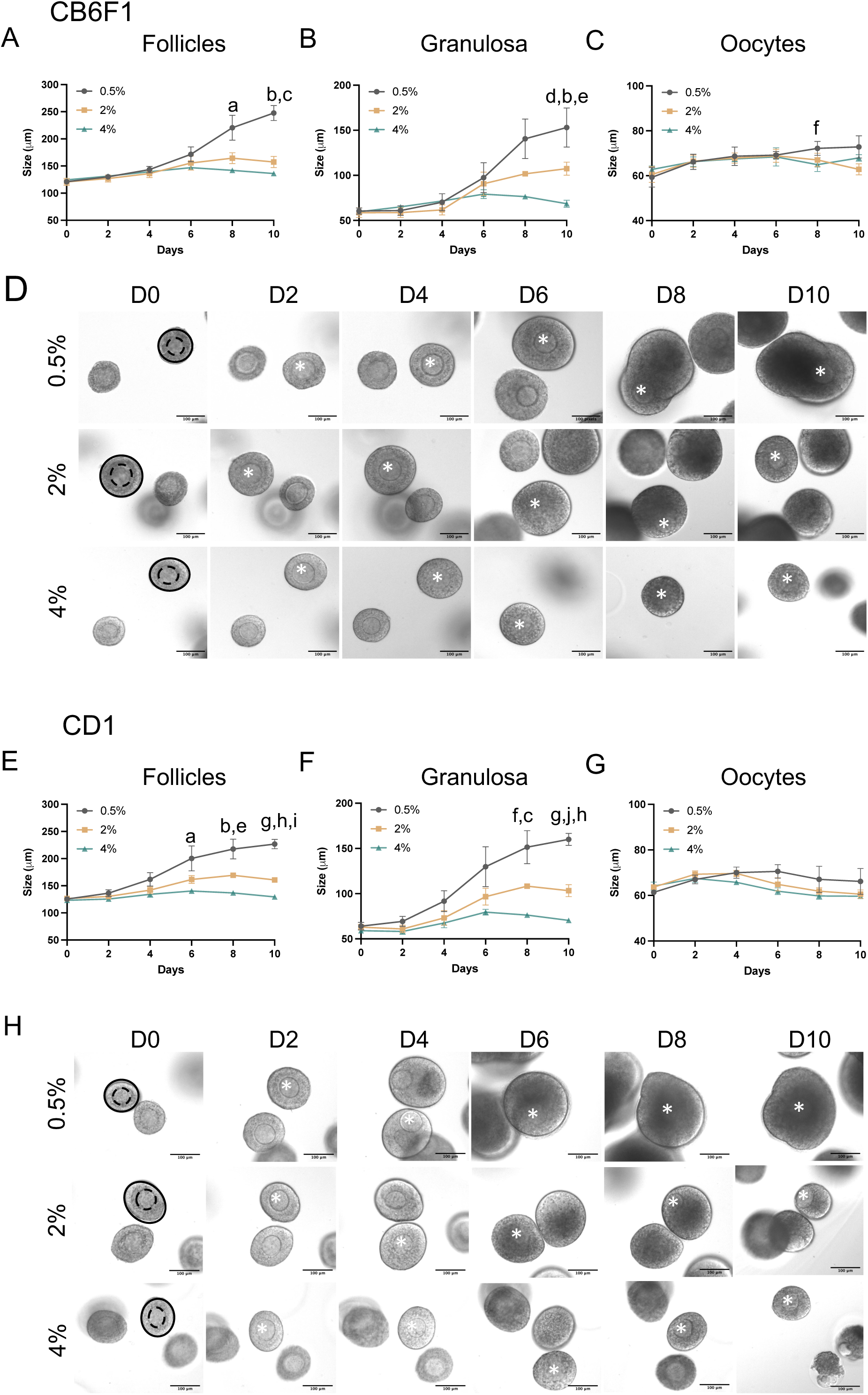
Follicle growth is severely impaired when follicles are cultured in stiff hydrogels. **(A-C)** Graph showing the growing dynamics of CB6F1 follicles (A), granulosa cells (B), and oocytes (C) during 10 days of culture. **(D)** Representative images of CB6F1 follicles grown with various alginate concentrations over a 10-days culture. **(E-G)** Quantifications of the CD1 follicle growth dynamics (E), granulosa cells (F), and oocytes (G) during the 10 days of culture. **(H)** Representative brightfield images of individual CD1 follicles during the 10 days of culture. Scale bar 100μm. At least N=3 independent experiments, with 10 follicles each, were performed. ^a^p<0.05 0.5% vs 4%. ^b^p<0.01 0.5% vs 2%. ^c^p<0.01 0.5% vs 4%. ^d^p<0.05 2% vs 4%. ^e^p<0.001 0.5% vs 4%. ^f^p<0.05 0.5% vs 2%. ^g^p<0.01 2% vs 4%. ^h^p<0.0001 0.5% vs 2%.^i^p<0.0001 0.5% vs 4%. ^j^p<0.001 0.5% vs 2%

We wondered whether the lack of follicle growth in stiff hydrogels reflected poor granulosa cell growth, poor oocyte growth, or both and thus analyzed the size of these two cell types separately. On day 0, granulosa cells from CB6F1 mice were 64.0 ± 4.16 µm in diameter and those from CD1 mice were 65.2 ± 2.98 µm. Granulosa cells from CB6F1 mice in 0.5% alginate reached a maximum size of 133.0±21.66 μm, and those from CD1 mice reached a maximum size of 160.0±6.71 μm. In 2% and 4% alginate, growth of granulosa cells from both strains was significantly impaired. On day 10, granulosa cells from CB6F1 had reached a final size of 107.4±7.26 μm in 2% alginate and 68.5±3.95 μm in 4% alginate. Granulosa cells from CD1 mice reached a final size of 103.4±6.74 μm in 2% alginate and 70.5±2.40 μm in 4% alginate (Figure 3 B and F). Additionally, beginning on day 4 of culture, some granulosa cells from both strains began to appear granular, a marker of cellular stress, and the percentage of granulated granulosa cells increased with higher alginate concentrations (Supplementary Figure 2). However, no major differences were observed in oocyte size, except for those from CB6F1 mice at day 10 (0.5% 72.8±2.49 μm, 2% 62.8±2.44 μm; 4% 67.9±1.47 μm; Figure 3 C, G). Altogether, these data show that the age-associated increased microenvironment stiffness significantly impairs growth and survival of follicles. Because the stiffness of 2% alginate matches that of age-associated ovarian stiffness, we focused our next experiments on comparing the effects of 0.5% and 2% alginate.

### Age-associated ovarian stiffness impairs granulosa cell viability, estradiol production and granulosa cell-oocyte communication

To determine whether granulosa cells cultured in the stiffer environment stopped growing or died in culture, we performed a live/dead assay. As expected, most of the follicles cultured in 0.5% alginate were alive on day 10 as indicated by the green fluorescent signal (Figure 4 A and B). In contrast, a red fluorescent signal (cell death marker) was present on day 6 in follicles cultured in 2% alginate, and the proportion of dead cells increased substantially during the culture period (Figure 4 A and B). The first cells to die were those in the external part of the follicle, and by the end of the culture period in 2% alginate, most granulosa cells and oocytes were dead. Consistent with the death noted in the granulosa cells, follicles cultured in 2% alginate produced less estradiol than follicles cultured in 0.5% alginate. By day 10 of culture, estradiol concentrations were 533-fold lower in CB6F1 follicles and 35.1-fold lower in CD1 follicles cultured in 2% alginate than in follicles cultured in 0.5% alginate (Figure 4 C and D).

**Figure 4:**
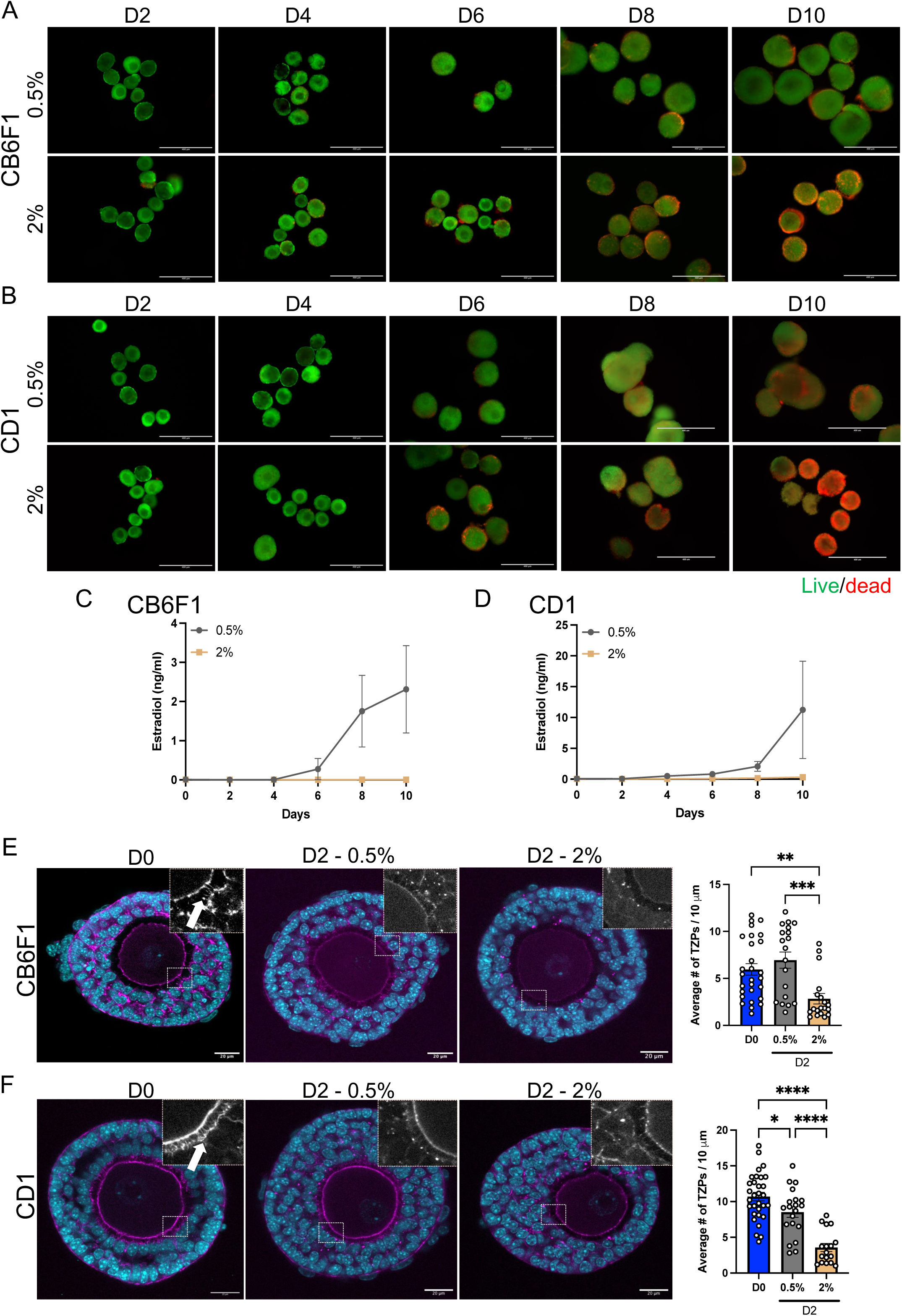
Granulosa cell survival and function are impaired by culture in stiff environments. (**A, B**) Representative images of CB6F1 (A) and CD1(B) mouse follicles cultured in 0.5% and 2% hydrogels analyzed with a live (green)/dead (red) assay. N≥10 follicles per condition. Scale bar, 400 μm. **(C, D)** Quantification of estradiol concentration in media from CB6F1(C) and CD1 (D) follicles. N≥3 independent experiments, with 10 follicles each. **(E, F)** Representative images of CB6F1 (E) and CD1 (F) follicles at day 0 (D0) and day 2 (D2) of culture stained with DNA dye (cyan) and Phalloidin (magenta). Insets show a magnification in greyscale. White arrows indicate TZPS. Graphs on the right show the number of TZPs per 10 μm. N≥3 independent experiments with 13 to 17 follicles per condition. *p<0.05, **p<0.01, ***p<0.001, ****p<0.0001.

Granulosa cells communicate with and provide nutrients to the oocyte that it cannot produce on its own [28]. Thus, we next investigated whether stiffness affected oocyte-granulosa transzonal projections (TZPs). These cytoplasmic projections emanate from granulosa cells and transport nutrients and other molecules to the oocyte via gap junctions [28]. The number of TZPs declines with age [29], so we hypothesized that stiffness could mediate this age-associated decline in TZP number. To test this idea, we cultured follicles in 0.5% and 2% alginate gels for 2 days. Freshly isolated follicles served as controls. We analyzed TZPs after two days of culture because follicle expansion hinders visualization of TZPs by confocal microscopy. Follicles from both mouse strains in 2% alginate had significantly fewer TZPs than follicles in 0.5% alginate (Figure 4 E-F). Together, these results suggest that a stiff microenvironment impairs granulosa function, viability, and ability to communicate with the oocyte via TZPs. The stiff environment first affects those cells in closest contact with the environment, and the negative impact is then transmitted to the internal layer of granulosa cells and the oocyte.

### Microenvironment stiffness impairs oocyte quality

To determine whether stiffness impairs oocyte viability, we removed follicles from the alginate on day 12 and mechanically isolated the oocytes. Because follicles in 2% alginate gels did not reach 180 μm in diameter, we did not induce *ex vivo* oocyte maturation and instead only examined whether oocytes reached the germinal vesicle (GV) stage. In line with previous observations, 73.5±4.91% and 76.4±4.61% of the isolated CB6F1 and CD1 oocytes, respectively, were classified as good-quality GV oocytes (GVs) [15] (Figure 5). However, in 2% alginate, only 7.9±3.96% and 31.1±6.87% of CB6F1 and CD1 oocytes, respectively, were good quality (Figure 5). These results suggest that follicles respond to changes in the biomechanical properties of the microenvironment and impair oocyte quality.

**Figure 5:**
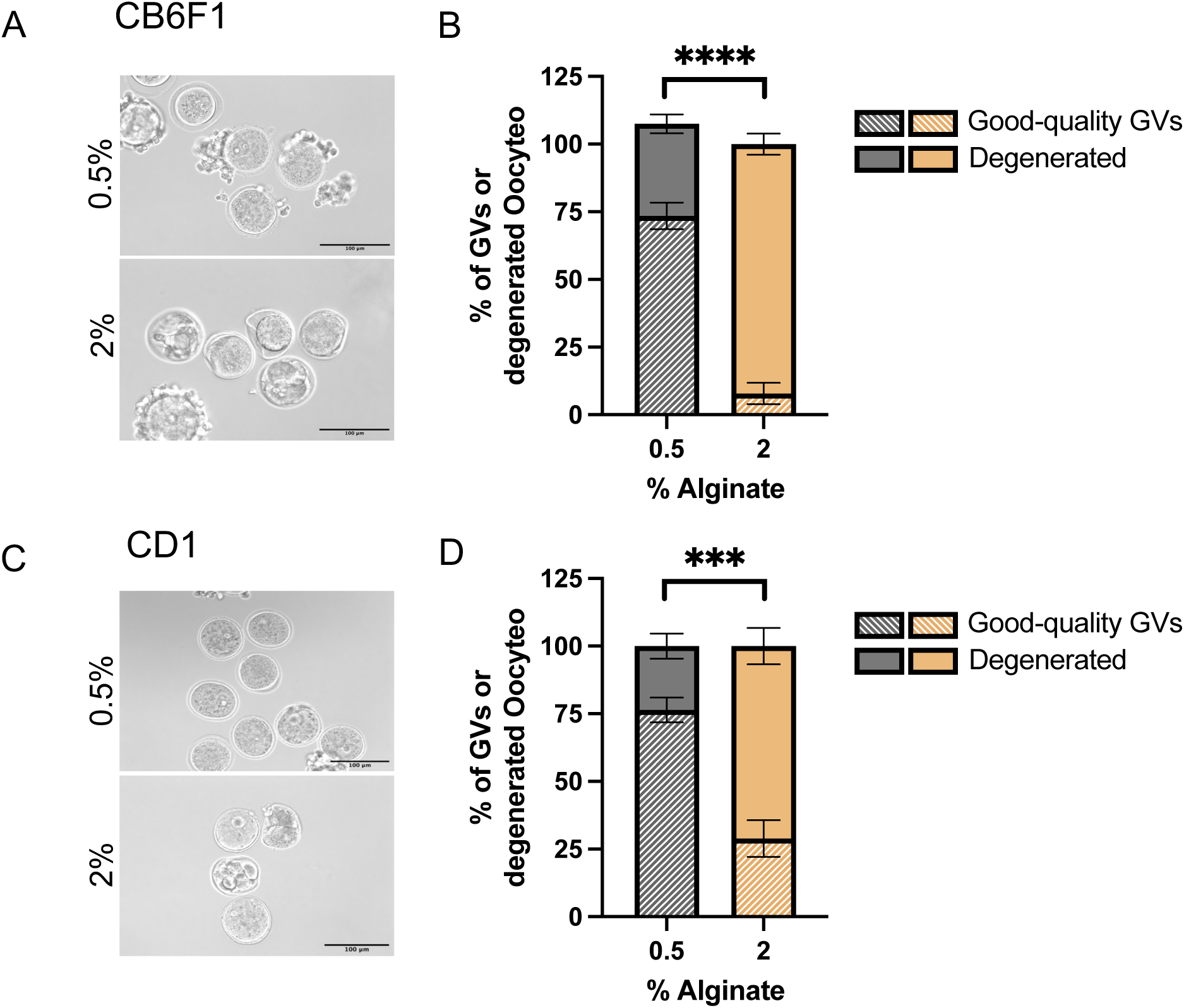
Morphological examination reveals that stiffness impairs oocyte quality. **(A, C)** Representative images of CB6F1 (A) and CD1 (C) oocytes isolated from follicles cultured in 0.5% and 2% alginate gels for 12 days. Scale bar, 100 μm. **(B, D)** Quantification of the percentage of CB6F1 (B) and CD1 (D) GVs that were good-quality or degenerated. N≥3 independent experiments, with 10 follicles each. ***p<0.001, ****p<0.0001.

### The follicle transcriptome quickly responds to a stiff microenvironment

To gain insights into early potential mechanisms that can explain the long-term effects of stiffness on granulosa cell and oocyte viability, we cultured secondary follicles in 0.5% and 2% alginate and processed them for RNA sequencing (RNA-seq) at 3, 6, 12, and 24 hours. At the 3 hour timepoint, we identified 51 differentially expressed genes (DEGs) between follicles in 0.5% and 2% alginate. The number of DEGs peaked at 6 hours (135 DEGS) and declined to reach a plateau by 12 hours (Figure 6A). Volcano plots revealed that at 3 hours, 40% DEGs were downregulated and 60% DEGs were upregulated in follicles cultured at 2% alginate relative to those in 0.5% alginate (Figure 6B). At 6 hours, 70% DEGs were downregulated and 30% were upregulated in follicles cultured in 2% alginate relative to those in 0.5% alginate (Figure 6E). We then performed gene ontology (GO) analysis for the down- and up-regulated genes to identify pathways affected by stiffness. At 3 hours, genes down-regulated in follicles cultured in 2% alginate were associated with phagosome maturation, morphogenesis, and development, including *Nog* (Noggin), which is an antagonist to bone morphogenetic proteins [30, 31] (Figure 6C and Supplementary Table S2). At 3 hours, up-regulated genes in 2% alginate were associated with the immune response mediated by chemokines and interleukin-1, as well as negative regulation of response to external stimuli, such as *Foxf1* (Forkhead Box F1), a transcription factor associated with matrix remodeling and cell junction organization [32–34] (Figure 6D and Supplementary Table S2).

**Figure 6:**
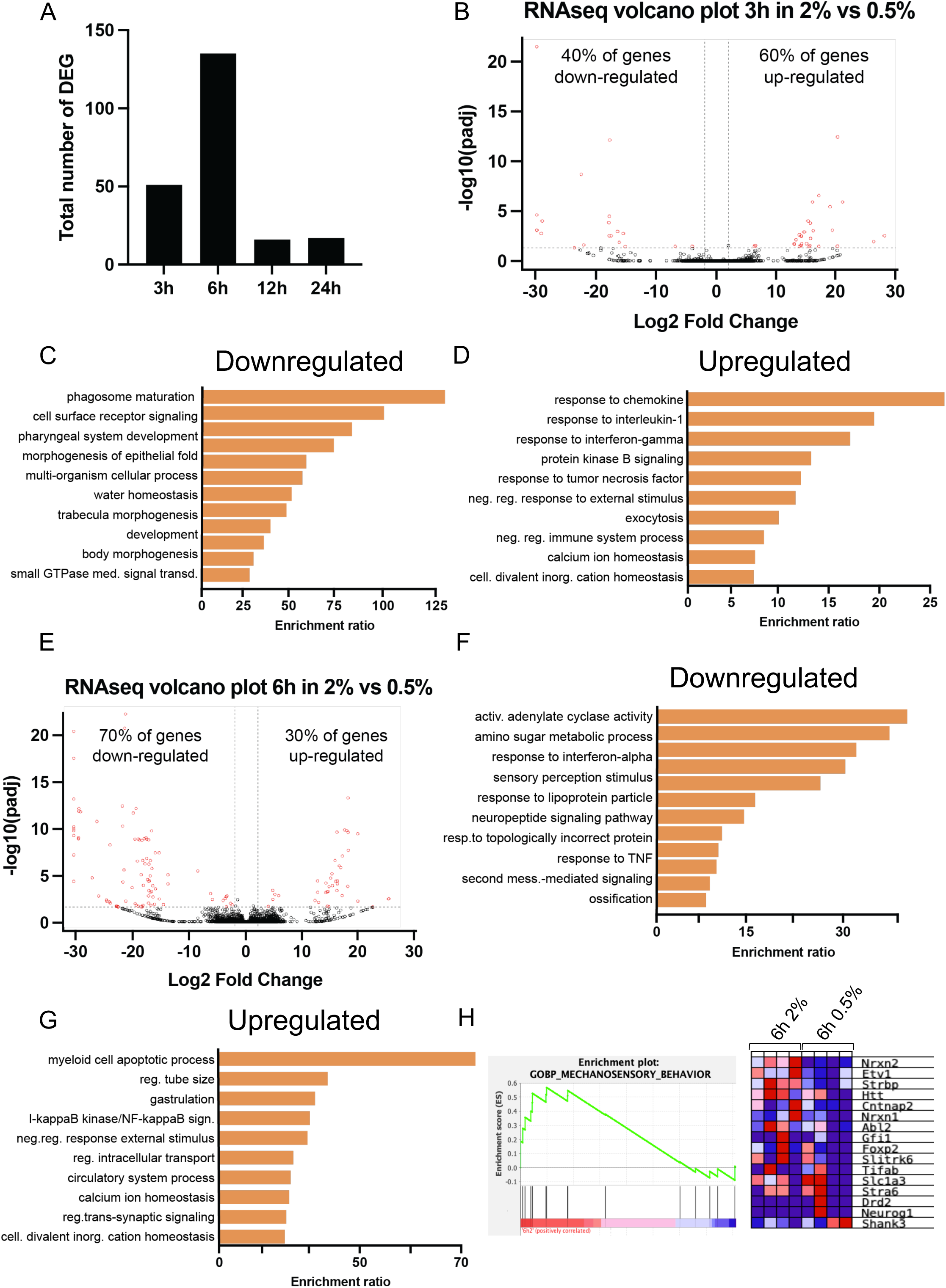
Genes involved in metabolism and inflammation are differentially expressed in follicles cultured in stiff environments. **(A)** Graph showing the total number of differentially expressed genes (DEG) between the 0.5% and 2% alginate encapsulated follicles at the indicated time points. **(B)** Volcano plot showing DEGs in follicles cultured in 2% alginate compared to those cultured in 0.5% at 3h. **(C, D)** Graphs showing GO analysis of downregulated (C) and upregulated (D) DEGs in 2% alginate gels compared to 0.5% at 3h. **(E)** Volcano plot showing DEGs in follicles cultured in 2% alginate compared to those cultured in 0.5% at 6h. **(F,G)** Graphs showing GO analysis of downregulated (F) and upregulated (G) DEGs in 2% alginate gels compared to 0.5% at 6h. **(H)** Graph showing the GSEA analysis of mechanosensory behavior pathway and its associated genes. Red indicates over-represented genes and blue indicates under-represented genes.

At 6 hours, metabolic-associated processes were the most downregulated pathways in follicles cultured in 2% alginate (Figure 6F), while up-regulated genes were associated with biological processes such as apoptosis, inflammation, and cellular adaptation to the environment (Figure 6G and Supplementary Table S3). Consistent with cellular adaptation to the environment, Gene Set Enrichment Analysis (GSEA) identified mechanosensory behavior as one of the most enriched pathways in the 2% condition compared to 0.5% at 6 hours. For example, *Nrxn2,* which is involved in cell adhesion, was upregulated (Figure 6I) [35]. No differences were noted in gene expression between follicles in 0.5% alginate and 2% alginate at 12 and 24 hours of culture, suggesting that follicles reach an equilibrium with the stiff microenvironment (Supplementary Figure S2A-D).

We also analyzed the genes that did not differ in expression across all comparisons and conditions and identified 10,798 such genes. In GO analysis, these genes were associated with biological processes important for oocyte maturation, cell division, metabolic processes, and cell survival (Supplementary Figure S4E). This finding was consistent with the fact that the stiff microenvironment did not impair follicle viability during the first 24 hours of culture (Figure 4).

### Pathways associated with inflammation and mechanosensation are upregulated in a stiff microenvironment

GO analysis at 3 hours revealed an increased inflammatory profile in the follicles cultured in 2% alginate (Figure 6D). To gain more insights into these inflammatory pathways, we performed GSEA analysis. Multiple inflammatory-associated pathways were enriched in follicles cultured in 2% alginate, including positive regulation of T helper cells, humoral immune response mediated by circulating immunoglobulins, eosinophil chemotaxis, and neutrophil migration (Supplementary Figure S3). Some of the most upregulated genes were *Card9* (Caspase recruitment domain family member 9), which is involved in NF-kappa B activation and cell apoptosis [36]; *Cfi* (complement factor I), involved in regulation of the complement cascade [37]; *Ccl3* (C-C motif chemokine ligand 3) involved in monocyte/macrophage chemotaxis [38]; and *Nckap1l* (NCK associated protein 1 like), involved in neutrophil migration [39].

At 6 hours, genes related to mechanosensation were elevated in follicles in 2% alginate. Thus, we analyzed the Hippo pathway, which is one of the main mechanotransduction pathways in cells [8]. We found no differences in expression of the Hippo-associated genes *Yap1, Taz,* or *Lats1/2* between follicles in 0.5% and 2% alginate at 3, 6, 12, or 24 hours (Supplementary Figure S4). However, the Yap1 downstream genes *Ctgf, Ankrd1,* and *Cyr61* were significantly downregulated in the 2% condition (Supplementary Figure S4), suggesting that stiffness impairs the Hippo signaling pathway. Therefore, from these observations, we conclude that follicles are highly mechanosensitive and that a stiff microenvironment impairs pathways that are essential for follicle development and growth such as metabolism and inflammation.

### A stiff microenvironment increases expression of genes associated with ECM remodeling

We next analyzed gene expression changes in a time-dependent manner by comparing DEGs at 3 and 24 hours in the 0.5% and 2% conditions. We first removed the 51 DEGs between the 0.5% and 2% conditions to obtain the most homogeneous population at 3 hours. We identified 1282 DEGs between follicles cultured in 0.5% alginate at 24 hours compared to 3 hours, with molecular pathways associated with protein and ion binding, and biological pathways associated with metabolic processes and response to stimuli (Figure 7A and B). Among the 39% of DEGs that were down-regulated at 24 hours relative to 3 hours (Figure 7C and Supplementary Table S4), GO analysis revealed genes involved in metabolic processes such as amino acid synthesis or hormone-related processes. A subset of the up-regulated genes classified in GO were known as “benzene and ether metabolism, peptidyl-arginine modification, and detoxification”. Because these GO pathways and did not match the molecular and biological processes results, we investigated the genes in each GO term. The benzene and ether metabolism term included genes such as acetyl-Coenzyme A acyltransferase 1B, glutathione S-transferase, hydroxyacid oxidase 2, and phospholipase A2, which are involved in fatty acid metabolism or acetyl CoA-transferase activity (Figure 7E). All of these genes are related to metabolic processes and energy production, which are essential for follicle growth.

**Figure 7:**
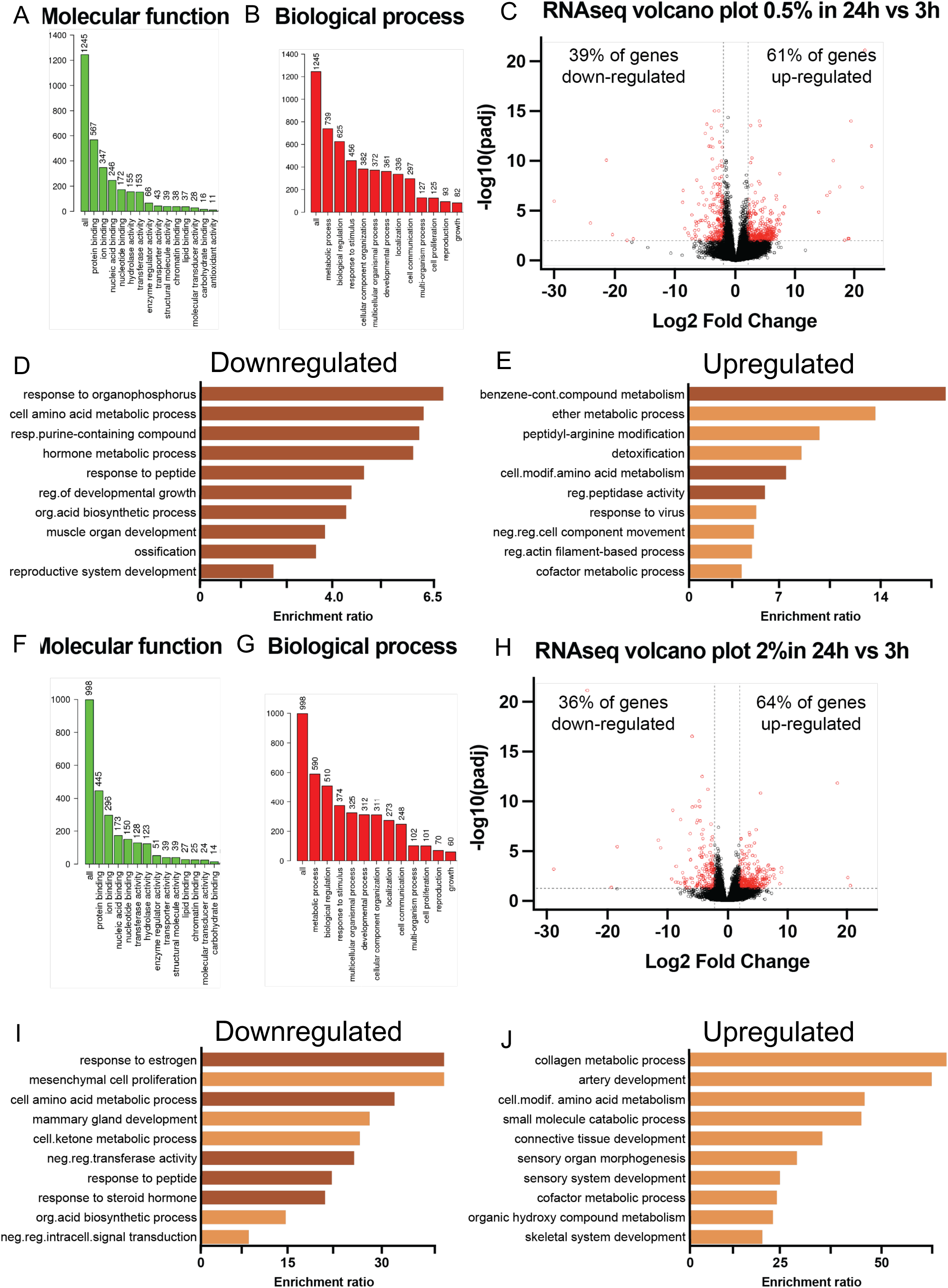
Stiffness changes the follicle transcriptome in a time-dependent manner. **(A,B)** Graphs showing the Gene Ontology enrichment of molecular functions (A) and biological processes (B) of DEGs obtained from follicles cultured in 0.5% alginate gels at 24h versus 3h. **(C)** Volcano plot showing DEGs in follicles cultured in 0.5% alginate gels at 24h versus 3h. **(D,E)** Graphs showing GO analysis of downregulated (D) and upregulated (E) DEGs at 24h compared to 3h in 0.5% alginate. **(F, G)** Graphs showing the Gene Ontology enrichment of molecular functions (F) and biological processes (G) of DEGs obtained from follicles cultured in 2% alginate gels at 24h versus 3h. **(H)** Volcano plot showing DEGs in follicles cultured in 2% alginate gels at 24h versus 3h. **(I, J)** Graphs showing GO analysis of downregulated (I) and upregulated (J) DEGs at 24h compared to 3h in 2% alginate.

We next analyzed time-dependent gene expression changes in 2% alginate and identified 1029 DEGs in follicles between 3 and 24 hours. These genes were associated with protein and ion biding, and metabolic and response to stimulus pathways as in the 0.5% condition (Figure 7F and G). Of the 36% of DEGs that were down-regulated at 24h (Figure 7H and Supplementary Table S5), GO analysis revealed genes that were associated with response to hormones, estrogens, and development (Figure 7I). These included *Eif4bp1* (eukaryotic translation initiation factor 4E binding protein 1), *Klf5* (Kruppel-like factor 5), *Ptgs2* (prostaglandin-endoperoxide synthease 2), and *Egr3* (early growth response 3). The up-regulated genes were associated with matrix remodeling, including collagen metabolic processes, and connective tissue development (Figure 7J). Analysis of the ECM-associated genes by GSEA revealed that pathways related to ECM matrix organization were enriched in 2% alginate. This included 91 significant genes such as *Adam8* (Desintegrin and Metalloproteinase Domain 8), *Itga2b* (Integrin Subunit Alpha 2b), *Col11a1* (Collagen Type XI Alpha 1 Chain), *Fbn2* (Fibrillin 2), and *Emilin* 3 (elastin microfibril interfacer 3) (Supplementary Figure S5 A). The most abundant ECM gene families included 34 genes related to collagen formation (Supplementary Figure S5 B and C). Together, these results suggest that a stiff microenvironment that mimics the one in the reproductively old ovary triggers the follicles to express genes that could remodel the ECM.

## Discussion

Together, the data presented here support the hypothesis that age-associated increased ovarian stiffness directly impairs follicle development and oocyte quality. *In vitro* culture of mouse follicles in alginate hydrogels that mimic the stiff environment of reproductively old ovaries impaired follicle survival and growth, granulosa cell proliferation, estradiol synthesis, and oocyte quality. RNA-seq analysis revealed that follicles are highly mechanosensitive, as we detected numerous DEGs as early as three hours of culture in stiff hydrogels. Stiffness induced expression of inflammatory and ECM-remodeling genes, suggesting bi-directional communication between the extra-follicular environment and the follicle. Because we isolated follicles from the ovary, the effects we observed were independent of the ECM composition and architecture. Moreover, all the phenotypes we observed mirror well-known markers of human ovarian aging such as increased follicle death, reduced oocyte quality, and increased inflammation [1, 40–43], suggesting that tissue stiffness can directly trigger age-associated ovarian dysfunction.

Increased ovarian stiffness has been reported in multiple reproductive-associated pathologies such as polycystic ovarian syndrome (PCOS) [44], primary ovarian insufficiency (POI) [45], and aging [6, 7], but how normal and pathologic stiffness modulate ovarian function remains poorly understood. Using the well-stablished *in vitro* follicle culture system [20], we synthesized alginate hydrogels that precisely mimic the stiffness values of ovaries from reproductively old mice [6], allowing us to determine the effect of physiologically relevant ovarian stiffness parameters on folliculogenesis. We found that microenvironment stiffness especially impaired follicle growth during the active expansion phase and led to decreased oocyte quality. These observations are in line with previous reports in which different concentrations of alginate and crosslinking conditions were tested [15]. An advantage of our work is that we tested the effect of stiffness in two different mouse strains, the hybrid CB6F1 mouse strain, and the outbred CD1 strain. This approach allowed us to discover that the impact of stiffness on folliculogenesis and oocyte quality is a robust and conserved mechanism since follicles form both strains were severely impacted by the age-associated increase in ovarian stiffness. The current study has greatly broadened our understanding of the mechanical cues in regulating folliculogenesis. We know that mechanical stress can impair follicle activation [11] and that ovarian drilling to relieve stress can improve follicle development in PCOS patients [46, 47]. Therefore, our data support the idea that strategies to modulate stiffness might improve ovarian function in women of advanced maternal age and patients with PCOS or POI.

Follicles are composed of granulosa cells and an oocyte. We found that granulosa cells were the first to be affected by the stiff microenvironment, as evidenced by reduced proliferation and increased cell death. This finding resembles the recent finding that theca cells, which surround the follicles, can exert a compressive stress on the follicles to regulate granulosa cell proliferation and growth [48]. The fact that granulosa cells are the first follicular cells to be affected by stiffness is not surprising as they are in contact with the basement membrane and extra-follicular environment. In addition, previous studies have revealed effects of aging on granulosa cells. For example, ovaries from both reproductively old mice [49] and ovaries from women of advanced maternal age [50] contain follicles with a smaller granulosa cell layer. In fact, transplanting an oocyte from a reproductively old mouse to granulosa cells from a reproductively young mouse can rejuvenate oocyte quality [51]. These results suggest that aging impairs oocyte quality by reducing granulosa cell quantity and quality. Our findings that increased ovarian stiffness impaired granulosa cell viability and follicle size supports the idea that a stiff microenvironment mediates the age-associated decline in granulosa cell viability, and therefore, oocyte quality, in ovaries from reproductively old mice.

To begin to examine the mechanism by which the effects of tissue stiffness are transmitted from granulosa cells to the oocyte, we examined transzonal projections (TZPs). These actin-based filaments mediate oocyte-granulosa communication and participate in delivering essential nutrients, such as pyruvate [28], from the granulosa cells to the oocyte. Additionally, the oocyte uses TZPs to signal to granulosa cells and regulate the rate of follicle development [52]. We found that follicles cultured in stiff hydrogel had significantly fewer TZPs than follicles cultured in soft hydrogel. Thus, TZPs, or the loss of TZPs, may transmit the stiffness signals from granulosa cells to the oocyte.

To further identify mechanisms by which ovarian stiffness impairs follicle development, we performed RNASeq analysis. By day 4 of culture, granulosa cells from follicles cultured in stiff hydrogels become granulated, a sign of cellular stress [53]. Therefore, we performed transcriptomic analysis within the first 24 hours of culture to identify early pathways that might trigger the observed phenotypes. We found that follicles respond to mechanical cues by altering gene expression within three hours, with DEGs between the 2% and 0.5% alginate conditions peaking at 6 hours. In GO pathway analysis, genes associated with response to the environment were differentially expressed in the two conditions. The Hippo pathway is one of the main mechanotransduction pathways of the cell, and it regulates essential biological processes such as cell proliferation, migration and differentiation [8, 54]. In follicles, the Hippo pathway regulates follicle activation, granulosa cell proliferation, and oocyte cytoplasmic maturation [47, 54–57], and single nucleotide polymorphisms in the Hippo pathway transcription factor *Yap1* have been identified in PCOS patients [58, 59]. Although we did not observe differences in expression of the core components of the Hippo signaling pathway (*Last1/2, Yap1, and Taz*) between 0.5% and 2% alginate, we did detect stiffness-associated decreased follicle expression of some of the Yap1 downstream pathways (*Ctgf*, *Ankrd1*, and *Cyr61*). This finding contrasts with observations in cell culture that a stiff environment induces nuclear transport and activation of Yap1. Future effort will be directed at exploring the interplay between age-associated ovarian stiffness and Hippo pathway activation or downregulation.

The transcriptomic analysis also revealed that multiple inflammatory pathways were upregulated in follicles cultured in 2% alginate. This finding is consistent with RNA-seq data revealing that inflammation genes were more highly expressed in follicles isolated from reproductively old mice than in those from young mice [49]. In addition, follicular fluid obtained from women of advanced maternal age is enriched in pro-inflammatory cytokines [42]. However, we do not know how this inflammatory environment is generated in the follicle. When cultured in stiff conditions, several cell types, including vascular smooth muscle, develop a pro-inflammatory environment [60]. Therefore, the stiff environment in the ovary might mediate the inflammatory profile observed in follicles from reproductively old mice and women of advanced maternal age. Our transcriptomic data also revealed that genes related to ECM remodeling were higher in follicles cultured in 2% alginate than in those cultured in 0.5% alginate. With advanced maternal age, the ovary becomes fibrotic and undergoes a complex ECM remodeling process [6, 18, 61–64]. Moreover, follicles can degrade and remodel the surrounding ECM to accommodate volumetric expansion [65–67]. Thus, further studies to investigate how the age-associated increase in stiffness affects follicular ECM remodeling and expansion are warranted.

Together, the work presented here demonstrates that the age-associated stiff microenvironment of the ovary directly impairs follicle survival, granulosa cell expansion, estradiol synthesis, and oocyte quality. Additionally, we provide initial evidence that these phenotypes might be triggered by inflammation and ECM remodeling. Importantly, all of the phenotypes we observed in this culture system have been observed in ovaries from women of advanced maternal age, suggesting that ovarian stiffness mediates the age-associated progressive decline in oocyte quantity and quality.

## Supplementary Figures

**Supplementary Figure 1: (A-C)** Representative images of the granular morphology in follicles cultured in 0.5% (A), 2% (B), and 4% (C) alginate at day 4. Scale bar, 50 μm. **(D-E)** Graphs showing the percentage of CB6F1 (D) and CD1 (E) follicles with a granulated morphology when cultured at 0.5%, 2% and 4% between day 4 and day 10 of culture.

**Supplementary Figure 2: (A, B)** Volcano plots showing DEGs in follicles cultured in 2% alginate compared to those cultured in 0.5% at 12h (A) and 24h (B). **(C, D)** Table showing representative DEGs at 12h (C) and 24h (D). **(E)** Graph showing the GO analysis of stably expressed genes across all comparisons.

**Supplementary Figure 3:** On the left in each panel, GSEA analysis of genes enriched in (**A**) T-helper, (**B**) humoral immune response, (**C**) eosinophil chemotaxis, and (**D**) neutrophil migration pathways. On the right in each panel, the cluster graphs only show genes that are significantly (p<0.05) enriched in 2% alginate compared to 0.5% alginate for each pathway.

**Supplementary Figure 4:** Normalized expression of genes associated with the Hippo signaling pathway in the indicated conditions and time points.

**Supplementary Figure 5: (A)** GSEA plot demonstrating up-regulated expression of the extracellular matrix organization pathway in 2% alginate at 24 h relative to 3 h. **(B, C)** GSEA plot demonstrating up-regulated expression of the collagen formation pathway (B) and its associated significantly enriched genes (C). Only genes that are significantly different are shown in the cluster plots (p<0.05).

**Supplementary Table 1: List of samples analyzed by RNA-seq.**

**Supplementary Table 2: List of DEGs between 0.5% and 2% at 3h.**

**Supplementary Table 3: List of DEGs between 0.5% and 2% at 6h.**

**Supplementary Table 4: List of DEGs between 3h and 24h in 0.5% alginate.**

**Supplementary Table 5: List of DEGs between 3h and 24h in 2% alginate.**

## Author contribution

S.P. and F.A. designed the methodology, performed experiments, data analysis and visualization, and wrote the manuscript. M.P. Performed TZPs imaging and quantification. F.A. conceived the study, supervised the study, and provided the resources and funding.

## Data availability statement

All data generated in this study are available upon request from the corresponding author. RNA-seq data will be uploaded to Gene Expression Omnibus repository when the manuscript is accepted for publication.

## Acknowledgements

We would like to acknowledge all the Amargant i Riera laboratory members as well as the members of the Center for Reproductive Health Sciences at Washington University in St. Louis, School of Medicine for their support and insightful discussion regarding this work. The authors also thank Dr. Deborah J. Frank for editing the manuscript. We thank Dr. Francesca E. Duncan for her mentoring as well as all the Duncan lab members for their support and feedback on this work. We would like to thank Dr. Serdar Bulun, Dr. Jerry Strauss, Dr. Qing Tu, Dr. Ariella Shikanov, Dr John Davis, and Dr. Luisa Iruela-Arispe for their insightful comments on this project and their support during the grant funding acquisition. Finally, we acknowledge that Figure 1C was created with BioRender.

