## Supplemental Figures and Tables for "Age-associated increased stiffness of the ovarian microenvironment impairs follicle development and oocyte quality and rapidly alters follicle gene expression"

### Supplementary Figure S1

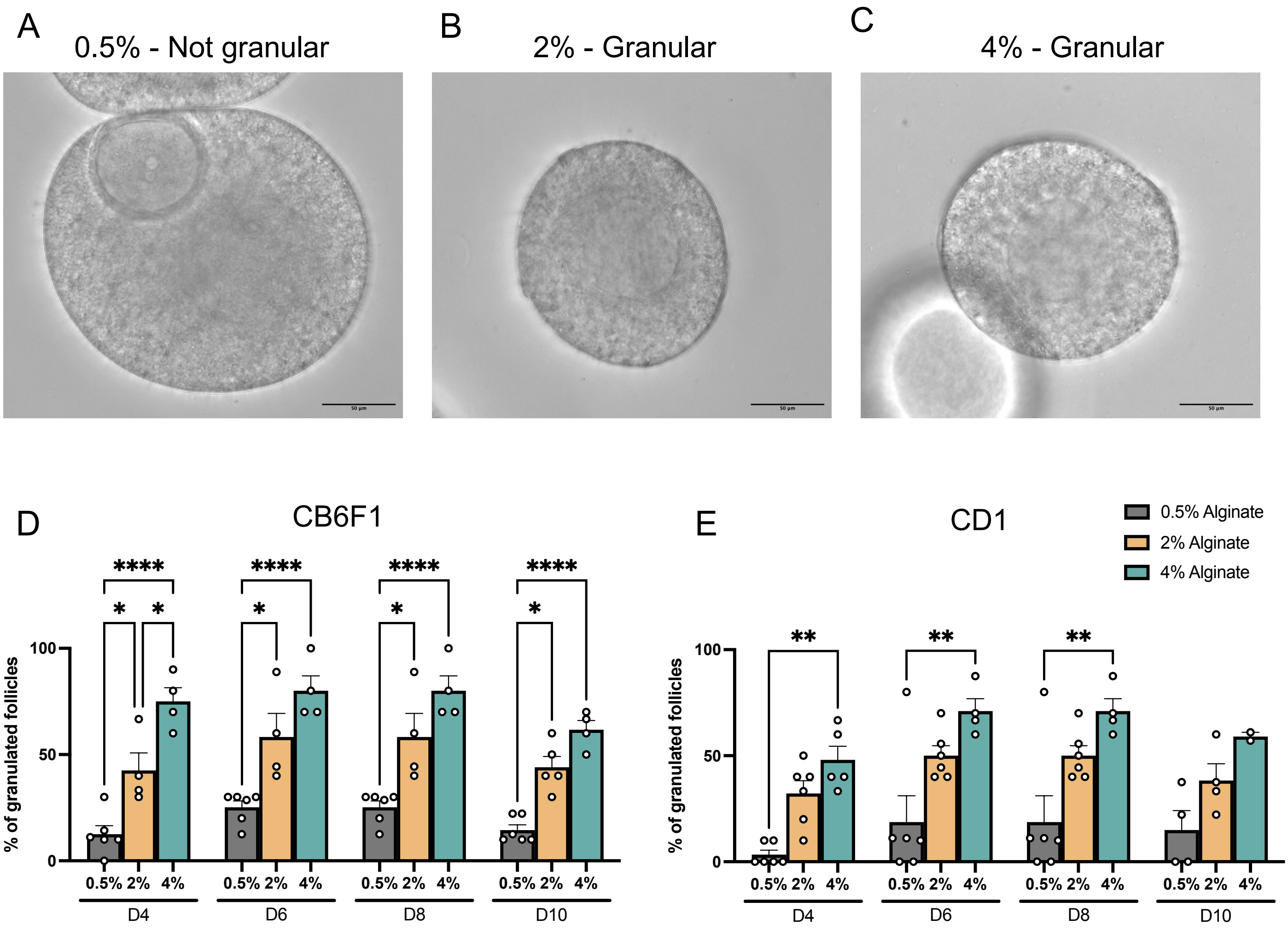

Supplementary Figure S2

A

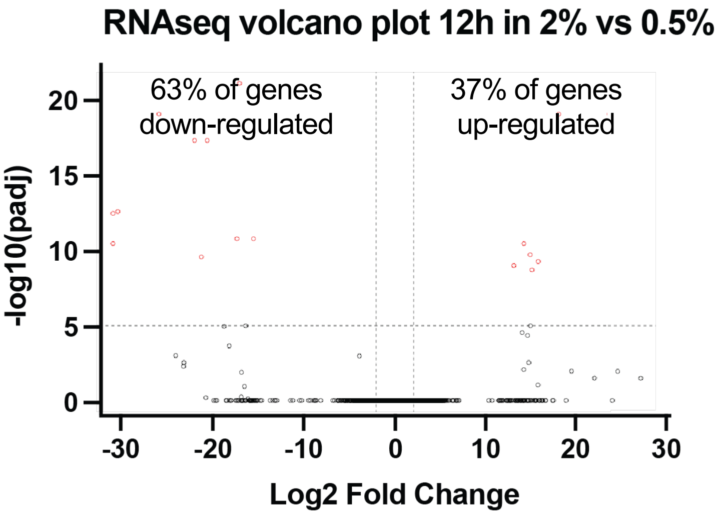


B

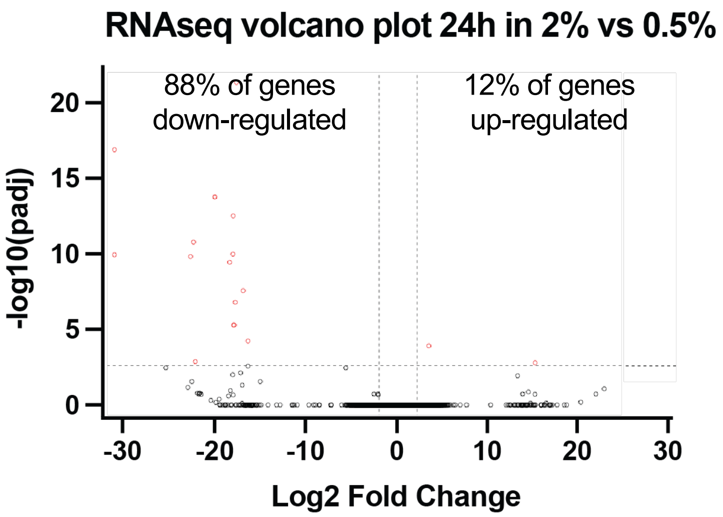


C

| Ensembl_ID | Symbol | Gene name | log2FoldChange | padj |
|---|---|---|---|---|
| ENSMUSG00000030235 | *Slco1c1* | Solute Carrier Organic Anion Transporter Family Member 1C1 | -25.102 | 1.048E-05 |
| ENSMUSG00000042812 | *Foxf1* | Forkhead Box F1 | 17.451 | 1.048E-05 |
| ENSMUSG00000024397 | *Aif1* | Allograft Inflammatory Factor 1 | -19.956 | 3.012E-05 |
| ENSMUSG00000043385 | *Olfr267* | Olfactory Receptor Family 2 Subfamily K Member 2 | -21.308 | 3.012E-05 |
| ENSMUSG00000096351 | *Samd11* | Sterile Alpha Motif Domain Containing 11 | -29.470 | 5.212E-04 |
| ENSMUSG00000036816 | *Atoh7* | Atonal BHLH Transcription Factor 7 | -29.986 | 5.589E-04 |
| ENSMUSG00000050134 | *Olfr430* | Olfactory Receptor Family 6 Subfamily N Member 2 | -16.798 | 1.541E-03 |
| ENSMUSG00000075104 | *Olfr1219* | Olfactory Receptor Family 4 Subfamily C Member 15 | -15.044 | 1.541E-03 |
| ENSMUSG00000038530 | *Rgs4* | Regulator Of G Protein Signaling 4 | 13.731 | 1.875E-03 |
| ENSMUSG00000024910 | *Ctsw* | Cathepsin W | 14.403 | 2.937E-03 |
| ENSMUSG00000021620 | *Acot12* | Acyl-CoA Thioesterase 12 | 12.677 | 4.513E-03 |

D

| Ensembl_ID | Symbol | Gene name | log2FoldChange | padj |
|---|---|---|---|---|
| ENSMUSG00000024027 | Glp1r | Glucagon Like Peptide 1 Receptor | -21.634 | 3.928E-06 |
| ENSMUSG00000002341 | Ncan | Neurocan | -16.371 | 1.609E-04 |
| ENSMUSG00000035283 | Adrb1 | Adrenoceptor Beta 1 | -17.317 | 2.211E-03 |
| ENSMUSG00000043385 | Olfr267 | Olfactory Receptor Family 2 Subfamily K Member 2 | -17.409 | 2.211E-03 |
| ENSMUSG00000045731 | Pnoc | Prepronociceptin | 3.257 | 1.086E-02 |
| ENSMUSG00000044055 | Otos | Otospiralin | 14.492 | 3.958E-02 |

E

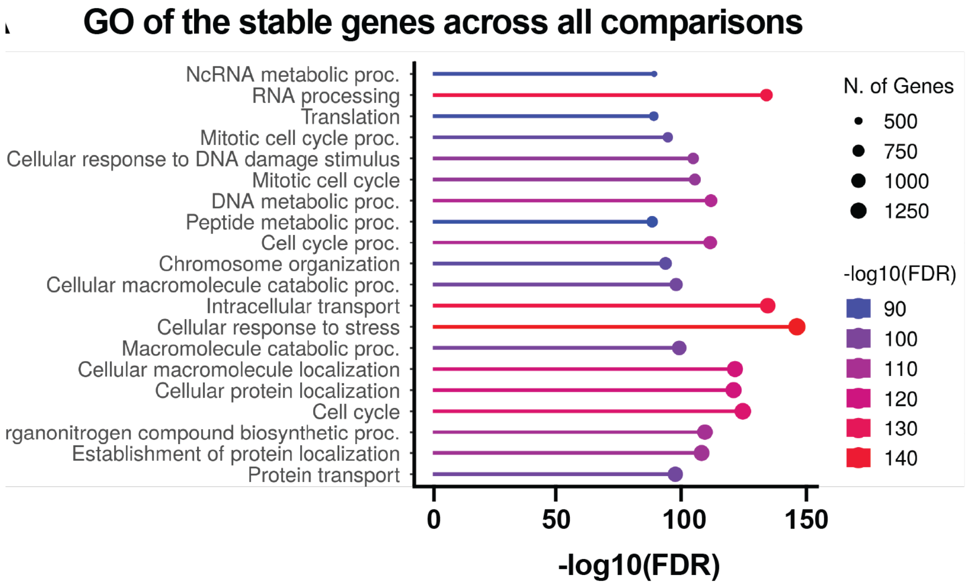

Supplementary Figure S3

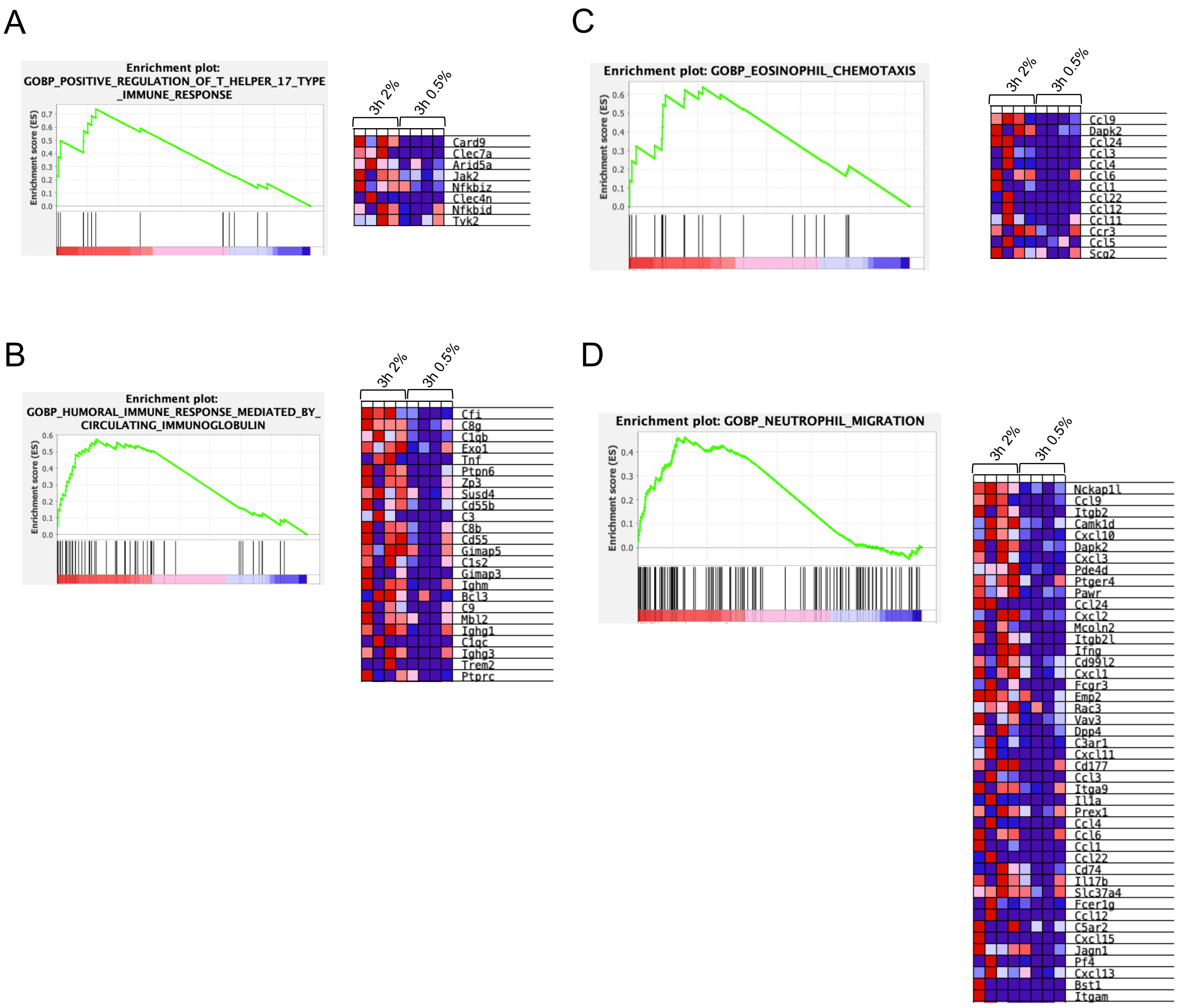

### Supplementary Figure S4

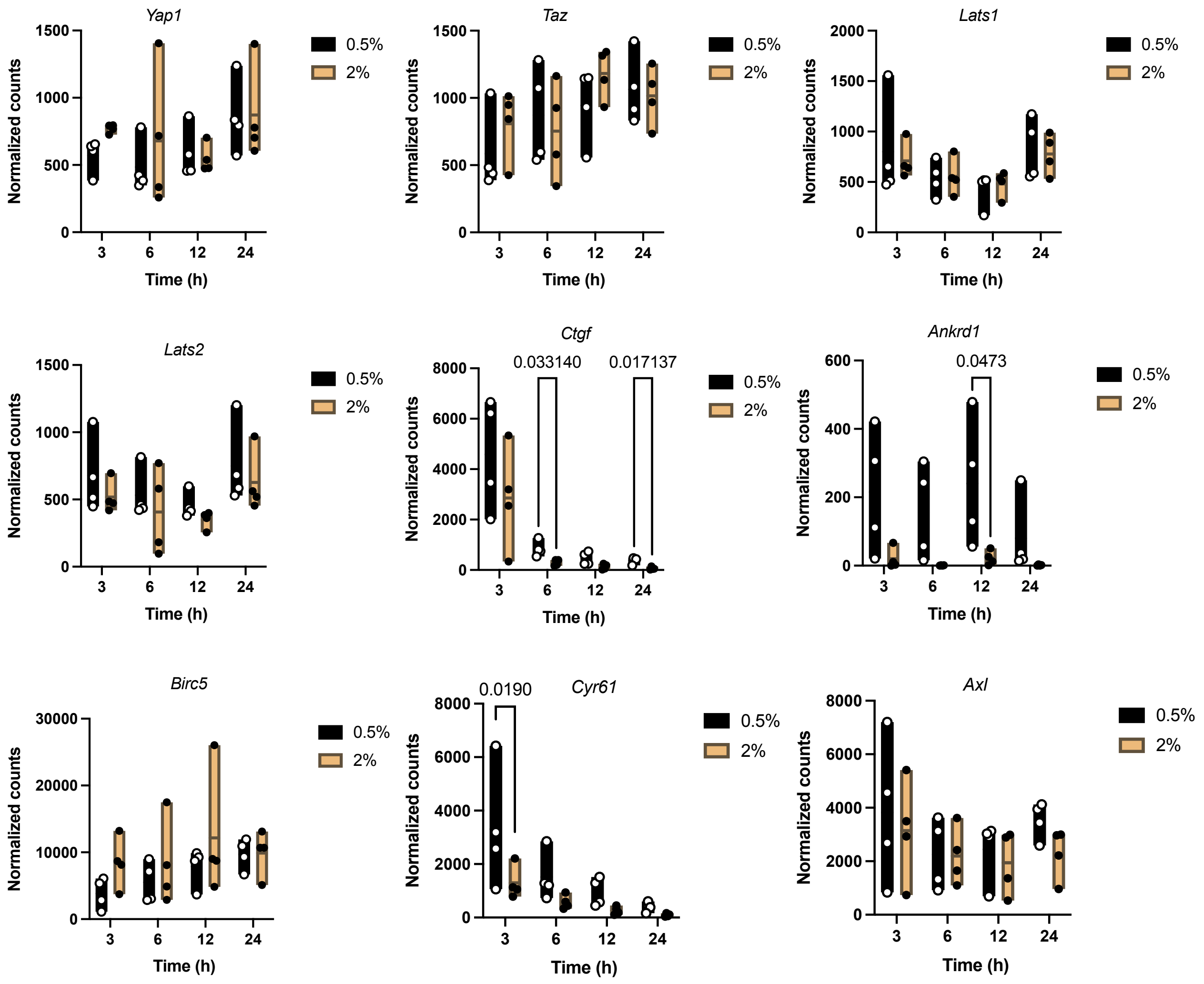

Supplementary Figure S5

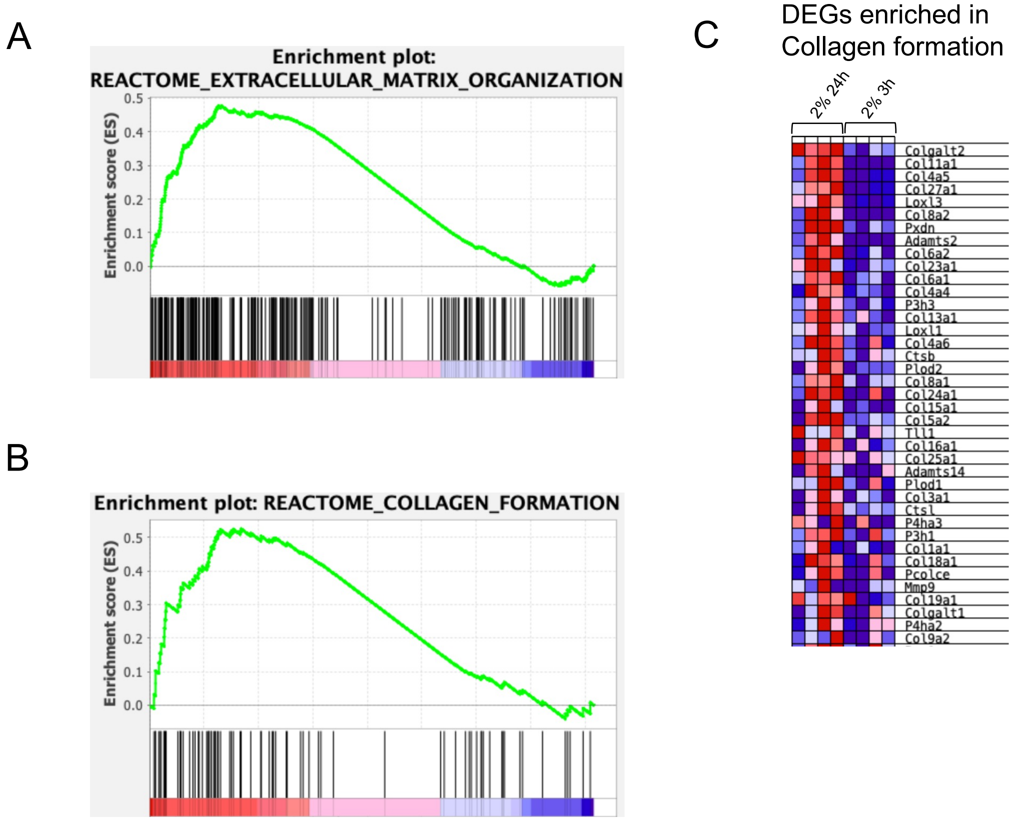

**Supplementary Table S1.**

**List of analyzed samples**

| Sample ID | Condition | Replicate |
|---|---|---|
| S0_5pct_12h_Replica_1_S5 | 0_5_12h | 1 |
| S0_5pct_12h_Replica_2_S13 | 0_5_12h | 2 |
| S0_5pct_12h_Replica_3_S21 | 0_5_12h | 3 |
| S0_5pct_12h_Replica_4_S29 | 0_5_12h | 4 |
| S0_5pct_24hh_Replica_1_S7 | 0_5_24h | 1 |
| S0_5pct_24hh_Replica_2_S15 | 0_5_24h | 2 |
| S0_5pct_24h_Replica_3_S23 | 0_5_24h | 3 |
| S0_5pct_24h_Replica_4_S31 | 0_5_24h | 4 |
| S0_5pct_3h_Replica_1_S1 | 0_5_3h | 1 |
| S0_5pct_3h_Replica_2_S9 | 0_5_3h | 2 |
| S0_5pct_3h_Replica_3_S17 | 0_5_3h | 3 |
| S0_5pct_3h_Replica_4_S25 | 0_5_3h | 4 |
| S0_5pct_6h_Replica_1_S3 | 0_5_6h | 1 |
| S0_5pct_6h_Replica_2_S11 | 0_5_6h | 2 |
| S0_5pct_6h_Replica_3_S19 | 0_5_6h | 3 |
| S0_5pct_6h_Replica_4_S27 | 0_5_6h | 4 |
| S2pct_12h_Replica_1_S6 | 2_12h | 1 |
| S2pct_12h_Replica_2_S14 | 2_12h | 2 |
| S2pct_12h_Replica_3_S22 | 2_12h | 3 |
| S2pct_12h_Replica_4_S30 | 2_12h | 4 |
| S2pct_24h_Replica_1_S8 | 2_24h | 1 |
| S2pct_24h_Replica_2_S16 | 2_24h | 2 |
| S2pct_24h_Replica_3_S24 | 2_24h | 3 |
| S2pct_24h_Replica_4_S32 | 2_24h | 4 |
| S2pct_3h_Replica_1_S2 | 2_3h | 1 |
| S2pct_3h_Replica_2_S10 | 2_3h | 2 |
| S2pct_3h_Replica_3_S18 | 2_3h | 3 |
| S2pct_3h_Replica_4_S26 | 2_3h | 4 |
| S2pct_6h_Replica_1_S4 | 2_6h | 1 |
| S2pct_6h_Replica_2_S12 | 2_6h | 2 |
| S2pct_6h_Replica_3_S20 | 2_6h | 3 |
| S2pct_6h_Replica_4_S28 | 2_6h | 4 |

**Supplementary Table S2.**

**List of DEGs  between 0.5% and 2% at 3h**

| Ensembl_ID | SYMBOL | og2FoldChang | pvalue | padj |
|---|---|---|---|---|
| ENSMUSG00000055069 | Rab39 | -30.000 | 5.66E-27 | 2.46E-22 |
| ENSMUSG00000028037 | Ifi44 | 20.185 | 1.45E-17 | 3.15E-13 |
| ENSMUSG00000028841 | Cnksr1 | -17.824 | 4.50E-17 | 6.52E-13 |
| ENSMUSG00000116220 | AC147041.1 | -22.603 | 1.73E-13 | 1.87E-09 |
| ENSMUSG00000074311 | Vmn1r139 | 17.062 | 2.98E-11 | 2.59E-07 |
| ENSMUSG00000058976 | Usp17lc | 16.021 | 1.89E-10 | 1.17E-06 |
| ENSMUSG00000069516 | Lyz2 | 21.039 | 1.81E-10 | 1.17E-06 |
| ENSMUSG00000040747 | Cd53 | 18.919 | 6.42E-10 | 3.48E-06 |
| ENSMUSG00000115509 | Gm49012 | -29.997 | 4.90E-09 | 2.36E-05 |
| ENSMUSG00000115108 | Gm49228 | -17.883 | 7.41E-09 | 3.22E-05 |
| ENSMUSG00000022875 | Kng1 | 15.263 | 2.64E-08 | 9.56E-05 |
| ENSMUSG00000091930 | Vmn2r52 | -29.081 | 2.62E-08 | 9.56E-05 |
| ENSMUSG00000053977 | Cd8a | -17.971 | 4.22E-08 | 1.41E-04 |
| ENSMUSG00000042812 | Foxf1 | 15.652 | 5.10E-08 | 1.58E-04 |
| ENSMUSG00000004730 | Adgre1 | 19.307 | 3.15E-07 | 8.16E-04 |
| ENSMUSG00000102701 | 9430083B18 | -29.991 | 3.19E-07 | 8.16E-04 |
| ENSMUSG00000102912 | Gm20731 | -29.990 | 3.18E-07 | 8.16E-04 |
| ENSMUSG00000087374 | Gm15457 | 16.137 | 3.80E-07 | 9.17E-04 |
| ENSMUSG00000085791 | Rpl30-ps9 | -16.508 | 4.78E-07 | 1.09E-03 |
| ENSMUSG00000002341 | Ncan | 14.540 | 6.02E-07 | 1.24E-03 |
| ENSMUSG00000098439 | Hm629797 | 14.691 | 5.79E-07 | 1.24E-03 |
| ENSMUSG00000032257 | Ankk1 | -29.233 | 8.88E-07 | 1.73E-03 |
| ENSMUSG00000116804 | AC115797.3 | -15.562 | 9.16E-07 | 1.73E-03 |
| ENSMUSG00000085479 | 9430073C21 | 13.811 | 1.33E-06 | 2.41E-03 |
| ENSMUSG00000059279 | Olfr224 | 28.011 | 1.83E-06 | 2.95E-03 |
| ENSMUSG00000113746 | Gm48242 | -17.602 | 1.80E-06 | 2.95E-03 |
| ENSMUSG00000114649 | 3110006O06 | -17.890 | 1.70E-06 | 2.95E-03 |
| ENSMUSG00000068048 | Rhox9 | 14.092 | 2.04E-06 | 3.16E-03 |
| ENSMUSG00000043329 | Gm8849 | 14.114 | 2.16E-06 | 3.23E-03 |
| ENSMUSG00000073929 | Trim30e-ps1 | 15.671 | 3.69E-06 | 5.33E-03 |
| ENSMUSG00000114157 | Gm47870 | 13.255 | 5.33E-06 | 7.46E-03 |
| ENSMUSG00000035352 | Ccl12 | 26.223 | 8.47E-06 | 1.15E-02 |
| ENSMUSG00000087681 | Gm13565 | -16.237 | 1.04E-05 | 1.36E-02 |
| ENSMUSG00000066688 | Gm13083 | 15.014 | 1.49E-05 | 1.90E-02 |
| ENSMUSG00000108891 | Olfr1291-ps1 | 14.162 | 1.55E-05 | 1.92E-02 |
| ENSMUSG00000028750 | Pla2g2c | 12.921 | 1.65E-05 | 1.99E-02 |
| ENSMUSG00000103877 | Gm21358 | 12.964 | 1.78E-05 | 2.09E-02 |
| ENSMUSG00000110162 | Gm45356 | -22.147 | 2.21E-05 | 2.53E-02 |
| ENSMUSG00000000982 | Ccl3 | 6.486 | 2.64E-05 | 2.86E-02 |
| ENSMUSG00000027880 | Slc25a54 | 14.065 | 2.63E-05 | 2.86E-02 |
| ENSMUSG00000002781 | Tmem143 | 1.987 | 2.84E-05 | 2.93E-02 |

| | | | | |
|---|---|---|---|---|
| ENSMUSG00000057903 | Olfr739 | 15.568 | 2.81E-05 | 2.93E-02 |
| ENSMUSG00000021768 | Dusp13 | 6.350 | 3.32E-05 | 3.35E-02 |
| ENSMUSG00000063779 | Chil4 | 20.156 | 3.39E-05 | 3.35E-02 |
| ENSMUSG00000048616 | Nog | -6.883 | 3.67E-05 | 3.48E-02 |
| ENSMUSG00000057766 | Ankrd29 | -4.069 | 3.84E-05 | 3.48E-02 |
| ENSMUSG00000076939 | Iglv3 | 13.885 | 3.85E-05 | 3.48E-02 |
| ENSMUSG00000097903 | Gm26679 | 17.051 | 3.80E-05 | 3.48E-02 |
| ENSMUSG00000116894 | AC154473.1 | 15.420 | 4.16E-05 | 3.69E-02 |
| ENSMUSG00000114752 | Gm35110 | -15.234 | 4.38E-05 | 3.81E-02 |
| ENSMUSG00000027483 | Bpifa1 | -23.704 | 5.70E-05 | 4.86E-02 |

**Supplementary Table S3.**

**List of DEGs between 0.5% and 2% at 6h**

| Ensembl_ID | SYMBOL | og2FoldChang | pvalue | padj |
|---|---|---|---|---|
| ENSMUSG00000029378 | Areg | -20.996 | 6.85E-24 | 2.40E-19 |
| ENSMUSG00000078922 | Tgtp1 | -21.151 | 2.55E-22 | 4.47E-18 |
| ENSMUSG00000114422 | Gm30411 | -30.000 | 7.15E-22 | 8.37E-18 |
| ENSMUSG00000030366 | Ceacam12 | -29.999 | 2.54E-19 | 2.23E-15 |
| ENSMUSG00000104698 | Gm42602 | 17.633 | 1.14E-15 | 8.02E-12 |
| ENSMUSG00000091923 | Gm8267 | -29.998 | 1.71E-15 | 9.99E-12 |
| ENSMUSG00000114157 | Gm47870 | -21.558 | 8.32E-15 | 4.18E-11 |
| ENSMUSG00000058976 | Usp17lc | -19.183 | 1.60E-14 | 7.03E-11 |
| ENSMUSG00000102676 | Gm37435 | -29.102 | 1.84E-14 | 7.20E-11 |
| ENSMUSG00000116884 | AC127341.5 | -29.106 | 3.63E-14 | 1.27E-10 |
| ENSMUSG00000035780 | Ugt2a3 | -28.837 | 4.38E-14 | 1.40E-10 |
| ENSMUSG00000109031 | Gm45201 | -19.679 | 6.23E-14 | 1.82E-10 |
| ENSMUSG00000099906 | Gm28653 | -25.991 | 3.85E-13 | 1.04E-09 |
| ENSMUSG00000028750 | Pla2g2c | -21.518 | 1.08E-12 | 2.70E-09 |
| ENSMUSG00000104821 | Gm42481 | -30.000 | 1.39E-12 | 3.24E-09 |
| ENSMUSG00000079356 | Mettl4-ps1 | -29.999 | 2.27E-12 | 4.98E-09 |
| ENSMUSG00000057710 | 9630041A04 | 17.041 | 2.86E-12 | 5.91E-09 |
| ENSMUSG00000021620 | Acot12 | 17.441 | 3.49E-12 | 6.82E-09 |
| ENSMUSG00000058114 | Olfr127 | -30.000 | 3.77E-12 | 6.97E-09 |
| ENSMUSG00000020395 | Itk | -16.480 | 4.08E-12 | 7.16E-09 |
| ENSMUSG00000038257 | Glra3 | 15.712 | 5.63E-12 | 9.41E-09 |
| ENSMUSG00000089854 | Gm16133 | 17.642 | 6.13E-12 | 9.78E-09 |
| ENSMUSG00000002341 | Ncan | 19.337 | 8.72E-12 | 1.33E-08 |
| ENSMUSG00000063394 | Olfr293 | -29.998 | 1.30E-11 | 1.90E-08 |
| ENSMUSG00000075105 | Olfr1218 | -29.156 | 2.24E-11 | 3.02E-08 |
| ENSMUSG00000109299 | Gm45164 | -17.450 | 2.16E-11 | 3.02E-08 |
| ENSMUSG00000088789 | Scarna13 | -17.769 | 2.77E-11 | 3.61E-08 |
| ENSMUSG00000077711 | AF357399 | -29.194 | 2.94E-11 | 3.68E-08 |
| ENSMUSG00000106543 | Gm43378 | -18.791 | 3.16E-11 | 3.83E-08 |
| ENSMUSG00000097444 | Gm10489 | -19.366 | 3.37E-11 | 3.95E-08 |
| ENSMUSG00000104903 | Gm43707 | -17.295 | 3.64E-11 | 4.12E-08 |
| ENSMUSG00000067813 | Xkr9 | -18.604 | 4.35E-11 | 4.63E-08 |
| ENSMUSG00000109446 | Gm9195 | -17.004 | 4.32E-11 | 4.63E-08 |
| ENSMUSG00000075020 | Mir670hg | -18.159 | 5.01E-11 | 5.17E-08 |
| ENSMUSG00000025804 | Ccr1 | -23.703 | 1.32E-10 | 1.32E-07 |
| ENSMUSG00000042812 | Foxf1 | 17.683 | 4.15E-10 | 4.05E-07 |
| ENSMUSG00000030669 | Calca | -15.845 | 7.41E-10 | 6.85E-07 |
| ENSMUSG00000105820 | Gm43400 | -15.171 | 7.38E-10 | 6.85E-07 |
| ENSMUSG00000115564 | Gm49199 | -29.997 | 1.15E-09 | 1.03E-06 |
| ENSMUSG00000021070 | Bdkrb2 | 16.661 | 1.49E-09 | 1.31E-06 |
| ENSMUSG00000068048 | Rhox9 | -17.346 | 2.51E-09 | 2.15E-06 |
| ENSMUSG00000103772 | Gm36933 | -18.025 | 3.92E-09 | 3.27E-06 |

| | | | | |
|---|---|---|---|---|
| ENSMUSG00000115330 | Gm49221 | 15.380 | 4.00E-09 | 3.27E-06 |
| ENSMUSG00000046717 | Igbp1b | -16.686 | 4.13E-09 | 3.30E-06 |
| ENSMUSG00000047108 | Dnajb7 | -18.940 | 4.83E-09 | 3.77E-06 |
| ENSMUSG00000103877 | Gm21358 | -17.633 | 8.56E-09 | 6.54E-06 |
| ENSMUSG00000101268 | 2010310C07 | -16.948 | 9.42E-09 | 7.04E-06 |
| ENSMUSG00000104406 | Gm38014 | 16.109 | 1.37E-08 | 1.00E-05 |
| ENSMUSG00000097934 | 6720483E21 | 16.380 | 1.98E-08 | 1.42E-05 |
| ENSMUSG00000081357 | Xlr4e-ps | -16.438 | 2.45E-08 | 1.72E-05 |
| ENSMUSG00000111312 | Gm47205 | -21.274 | 3.36E-08 | 2.31E-05 |
| ENSMUSG00000024803 | Ankrd1 | -8.461 | 4.22E-08 | 2.85E-05 |
| ENSMUSG00000090304 | Vmn2r99 | 14.851 | 7.63E-08 | 5.06E-05 |
| ENSMUSG00000066688 | Gm13083 | -18.310 | 9.47E-08 | 6.16E-05 |
| ENSMUSG00000087622 | Gm12290 | -13.641 | 9.86E-08 | 6.30E-05 |
| ENSMUSG00000111065 | Gm47676 | -16.716 | 1.25E-07 | 7.82E-05 |
| ENSMUSG00000110738 | Gm47092 | -26.797 | 1.92E-07 | 1.18E-04 |
| ENSMUSG00000022878 | Adipoq | 14.611 | 2.09E-07 | 1.27E-04 |
| ENSMUSG00000086061 | Gm13075 | -17.170 | 2.89E-07 | 1.72E-04 |
| ENSMUSG00000027857 | Tshb | 15.772 | 3.90E-07 | 2.25E-04 |
| ENSMUSG00000053977 | Cd8a | -16.891 | 3.93E-07 | 2.25E-04 |
| ENSMUSG00000084899 | Gm15344 | 12.356 | 3.98E-07 | 2.25E-04 |
| ENSMUSG00000102713 | Gm29994 | -16.990 | 4.31E-07 | 2.40E-04 |
| ENSMUSG00000027801 | Tm4sf4 | 15.496 | 4.51E-07 | 2.47E-04 |
| ENSMUSG00000110363 | Gm45576 | -29.999 | 4.57E-07 | 2.47E-04 |
| ENSMUSG00000115686 | Gm49310 | -21.460 | 5.29E-07 | 2.81E-04 |
| ENSMUSG00000103159 | F830112A20 | 13.784 | 6.67E-07 | 3.50E-04 |
| ENSMUSG00000070704 | Ugt2b36 | 15.787 | 7.99E-07 | 4.13E-04 |
| ENSMUSG00000070077 | Mir491 | -13.636 | 8.23E-07 | 4.19E-04 |
| ENSMUSG00000113746 | Gm48242 | -17.910 | 1.12E-06 | 5.60E-04 |
| ENSMUSG00000096655 | 1700065D16 | 14.934 | 1.21E-06 | 5.97E-04 |
| ENSMUSG00000111874 | Gm36539 | -16.282 | 1.25E-06 | 6.11E-04 |
| ENSMUSG00000110702 | Gm45767 | 17.581 | 1.45E-06 | 6.99E-04 |
| ENSMUSG00000070501 | Ifi214 | 14.348 | 1.53E-06 | 7.25E-04 |
| ENSMUSG00000096690 | Olfr1296-ps1 | -19.863 | 1.91E-06 | 8.93E-04 |
| ENSMUSG00000097023 | Mir9-3hg | -15.790 | 2.49E-06 | 1.15E-03 |
| ENSMUSG00000033208 | S100b | 4.577 | 3.47E-06 | 1.58E-03 |
| ENSMUSG00000029915 | Clec5a | 15.702 | 3.65E-06 | 1.65E-03 |
| ENSMUSG00000114752 | Gm35110 | -16.962 | 3.70E-06 | 1.65E-03 |
| ENSMUSG00000108228 | 6430584L05 | 14.166 | 4.50E-06 | 1.97E-03 |
| ENSMUSG00000025431 | Crisp1 | -16.488 | 4.66E-06 | 2.02E-03 |
| ENSMUSG00000110618 | Gm39822 | 14.031 | 5.10E-06 | 2.18E-03 |
| ENSMUSG00000114253 | Gm47798 | -16.716 | 5.68E-06 | 2.37E-03 |
| ENSMUSG00000114382 | Gm4814 | 12.367 | 5.67E-06 | 2.37E-03 |
| ENSMUSG00000048764 | Tmprss11f | 13.739 | 6.00E-06 | 2.48E-03 |
| ENSMUSG00000030827 | Fgf21 | -6.230 | 7.41E-06 | 3.03E-03 |
| ENSMUSG00000024548 | Setbp1 | 4.930 | 8.97E-06 | 3.62E-03 |
| ENSMUSG00000020057 | Dram1 | 5.593 | 1.18E-05 | 4.65E-03 |
| ENSMUSG00000083228 | Gm11779 | 16.614 | 1.17E-05 | 4.65E-03 |

| | | | | |
|---|---|---|---|---|
| ENSMUSG00000004837 | Grap | -17.021 | 1.27E-05 | 4.90E-03 |
| ENSMUSG00000099891 | Gm5575 | -3.388 | 1.27E-05 | 4.90E-03 |
| ENSMUSG00000056600 | Olfr90 | -25.676 | 1.30E-05 | 4.97E-03 |
| ENSMUSG00000053916 | Nanp | -3.515 | 1.51E-05 | 5.71E-03 |
| ENSMUSG00000086118 | Gm14169 | -16.235 | 1.66E-05 | 6.18E-03 |
| ENSMUSG00000034839 | Larp6 | -3.695 | 2.25E-05 | 8.33E-03 |
| ENSMUSG00000096773 | Olfr675 | 24.748 | 2.41E-05 | 8.81E-03 |
| ENSMUSG00000109548 | Gm45066 | 24.652 | 2.59E-05 | 9.39E-03 |
| ENSMUSG00000055159 | 4930583K01 | -15.143 | 2.63E-05 | 9.43E-03 |
| ENSMUSG00000047363 | Cstad | 4.596 | 2.91E-05 | 1.03E-02 |
| ENSMUSG00000078921 | Tgtp2 | -5.910 | 2.96E-05 | 1.04E-02 |
| ENSMUSG00000085385 | Snhg17 | -1.536 | 3.17E-05 | 1.10E-02 |
| ENSMUSG00000095248 | Olfr681 | -24.326 | 3.44E-05 | 1.18E-02 |
| ENSMUSG00000022875 | Kng1 | 11.899 | 3.75E-05 | 1.28E-02 |
| ENSMUSG00000020154 | Ptprb | -4.054 | 3.80E-05 | 1.28E-02 |
| ENSMUSG00000104527 | Gm37332 | 19.351 | 4.41E-05 | 1.46E-02 |
| ENSMUSG00000107140 | Gm43811 | -20.954 | 4.38E-05 | 1.46E-02 |
| ENSMUSG00000102597 | Gm38382 | -24.220 | 4.66E-05 | 1.53E-02 |
| ENSMUSG00000058400 | Qrfpr | -23.782 | 5.37E-05 | 1.74E-02 |
| ENSMUSG00000055109 | Gm15155 | 12.532 | 5.83E-05 | 1.88E-02 |
| ENSMUSG00000094745 | Olfr954 | -17.018 | 6.85E-05 | 2.19E-02 |
| ENSMUSG00000105701 | Gm42587 | -23.665 | 6.93E-05 | 2.19E-02 |
| ENSMUSG00000085379 | 2310058D17 | -2.771 | 7.36E-05 | 2.31E-02 |
| ENSMUSG00000063779 | Chil4 | -19.337 | 7.97E-05 | 2.48E-02 |
| ENSMUSG00000070306 | Ccdc153 | -15.982 | 9.16E-05 | 2.82E-02 |
| ENSMUSG00000047225 | Olfr684 | -14.433 | 9.71E-05 | 2.97E-02 |
| ENSMUSG00000027499 | Pkia | -2.477 | 1.03E-04 | 3.11E-02 |
| ENSMUSG00000073902 | Gm1966 | -19.142 | 1.13E-04 | 3.36E-02 |
| ENSMUSG00000104937 | Gm43057 | -17.974 | 1.13E-04 | 3.36E-02 |
| ENSMUSG00000105811 | Gm42707 | -16.594 | 1.14E-04 | 3.36E-02 |
| ENSMUSG00000061410 | Zcchc14 | 1.183 | 1.35E-04 | 3.94E-02 |
| ENSMUSG00000109147 | 4930431P19 | -1.890 | 1.36E-04 | 3.96E-02 |
| ENSMUSG00000109733 | Gm33940 | -22.621 | 1.43E-04 | 4.11E-02 |
| ENSMUSG00000067642 | Adgrf3 | 14.033 | 1.47E-04 | 4.18E-02 |
| ENSMUSG00000095903 | Olfr968 | -18.814 | 1.49E-04 | 4.21E-02 |
| ENSMUSG00000112139 | Gm47431 | -22.543 | 1.51E-04 | 4.23E-02 |
| ENSMUSG00000100315 | 1700031P21 | -22.526 | 1.52E-04 | 4.25E-02 |
| ENSMUSG00000085763 | Smc2os | 3.802 | 1.57E-04 | 4.34E-02 |
| ENSMUSG00000037390 | Muc3 | 14.296 | 1.63E-04 | 4.46E-02 |
| ENSMUSG00000036052 | Dnajb5 | -3.482 | 1.66E-04 | 4.51E-02 |
| ENSMUSG00000075107 | Olfr1216 | -22.353 | 1.71E-04 | 4.62E-02 |
| ENSMUSG00000104938 | Gm42840 | -18.058 | 1.73E-04 | 4.64E-02 |
| ENSMUSG00000062082 | Cd200r4 | -22.306 | 1.77E-04 | 4.66E-02 |
| ENSMUSG00000086675 | Plxna4os2 | -22.314 | 1.76E-04 | 4.66E-02 |
| ENSMUSG00000022129 | Dct | -18.275 | 1.79E-04 | 4.70E-02 |
| ENSMUSG00000042433 | Pih1h3b | 21.942 | 1.81E-04 | 4.71E-02 |

**Supplementary Table S4.**

**List of DEGs  between 3h-24h in 0.5%**

| Ensembl_ID | SYMBOL | og2FoldChang | pvalue | padj |
|---|---|---|---|---|
| ENSMUSG00000069917 | Hba-a2 | -29.630 | 6.30E-07 | 1.20E-04 |
| ENSMUSG00000024027 | Glp1r | -21.106 | 2.44E-10 | 2.32E-07 |
| ENSMUSG00000073940 | Hbb-bt | -20.113 | 7.21E-04 | 1.98E-02 |
| ENSMUSG00000042433 | Pih1h3b | -17.736 | 2.74E-03 | 4.74E-02 |
| ENSMUSG00000096606 | Tpbgl | -16.707 | 1.95E-03 | 3.77E-02 |
| ENSMUSG00000021680 | Crhbp | -8.674 | 8.27E-07 | 1.42E-04 |
| ENSMUSG00000022296 | Baalc | -8.581 | 7.84E-08 | 2.34E-05 |
| ENSMUSG00000020911 | Krt19 | -8.326 | 1.83E-04 | 7.81E-03 |
| ENSMUSG00000115620 | Gm45924 | -7.793 | 4.85E-04 | 1.51E-02 |
| ENSMUSG00000001622 | Csn3 | -7.643 | 8.69E-04 | 2.25E-02 |
| ENSMUSG00000062345 | Serpinb2 | -7.162 | 1.10E-05 | 9.77E-04 |
| ENSMUSG00000031635 | Anxa10 | -7.141 | 1.81E-03 | 3.61E-02 |
| ENSMUSG00000085183 | Wincr1 | -6.700 | 2.86E-04 | 1.06E-02 |
| ENSMUSG00000068697 | Myoz1 | -6.657 | 6.76E-04 | 1.90E-02 |
| ENSMUSG00000030890 | Ilk | -6.601 | 1.88E-03 | 3.69E-02 |
| ENSMUSG00000028031 | Dkk2 | -6.525 | 4.20E-05 | 2.70E-03 |
| ENSMUSG00000070645 | Ren1 | -6.464 | 1.04E-03 | 2.48E-02 |
| ENSMUSG00000028328 | Tmod1 | -6.436 | 1.00E-04 | 5.07E-03 |
| ENSMUSG00000028172 | Tacr3 | -6.400 | 5.47E-05 | 3.27E-03 |
| ENSMUSG00000050359 | Sprr1a | -6.364 | 9.58E-09 | 4.86E-06 |
| ENSMUSG00000111467 | Gm47775 | -6.081 | 2.50E-04 | 9.58E-03 |
| ENSMUSG00000032487 | Ptgs2 | -6.057 | 1.20E-06 | 1.88E-04 |
| ENSMUSG00000050541 | Adra1b | -6.044 | 4.76E-05 | 2.94E-03 |
| ENSMUSG00000007946 | Phox2a | -5.896 | 2.49E-03 | 4.46E-02 |
| ENSMUSG00000033730 | Egr3 | -5.789 | 3.37E-05 | 2.30E-03 |
| ENSMUSG00000029304 | Spp1 | -5.643 | 4.53E-08 | 1.45E-05 |
| ENSMUSG00000022602 | Arc | -5.494 | 3.32E-05 | 2.28E-03 |
| ENSMUSG00000021403 | Serpinb9b | -5.372 | 1.13E-07 | 3.23E-05 |
| ENSMUSG00000047638 | Nr1h4 | -5.138 | 1.45E-06 | 2.17E-04 |
| ENSMUSG00000097084 | Foxl1 | -5.092 | 1.10E-03 | 2.57E-02 |
| ENSMUSG00000046223 | Plaur | -4.973 | 2.34E-12 | 3.93E-09 |
| ENSMUSG00000033174 | Mgll | -4.960 | 3.39E-05 | 2.31E-03 |
| ENSMUSG00000022805 | Maats1 | -4.901 | 1.36E-05 | 1.16E-03 |
| ENSMUSG00000032394 | Igdcc3 | -4.854 | 2.57E-03 | 4.54E-02 |
| ENSMUSG00000030854 | Ptpn5 | -4.793 | 4.77E-07 | 9.55E-05 |
| ENSMUSG00000050335 | Lgals3 | -4.733 | 3.05E-13 | 9.52E-10 |
| ENSMUSG00000027068 | Dhrs9 | -4.654 | 1.71E-03 | 3.47E-02 |
| ENSMUSG00000024912 | Fosl1 | -4.628 | 1.84E-05 | 1.47E-03 |
| ENSMUSG00000097746 | Gm6225 | -4.624 | 2.87E-03 | 4.89E-02 |
| ENSMUSG00000047171 | Helt | -4.551 | 1.54E-03 | 3.21E-02 |
| ENSMUSG00000074199 | Krtdap | -4.442 | 5.39E-05 | 3.24E-03 |
| ENSMUSG00000010797 | Wnt2 | -4.404 | 1.22E-03 | 2.76E-02 |

| | | | | |
|---|---|---|---|---|
| ENSMUSG00000050578 | Mmp13 | -4.400 | 1.99E-03 | 3.84E-02 |
| ENSMUSG00000015652 | Steap1 | -4.389 | 2.14E-11 | 2.74E-08 |
| ENSMUSG00000030827 | Fgf21 | -4.371 | 1.20E-04 | 5.75E-03 |
| ENSMUSG00000037868 | Egr2 | -4.306 | 1.07E-04 | 5.29E-03 |
| ENSMUSG00000003283 | Hck | -4.214 | 2.48E-06 | 3.24E-04 |
| ENSMUSG00000063531 | Sema3e | -4.170 | 1.71E-03 | 3.46E-02 |
| ENSMUSG00000031297 | Slc7a3 | -4.143 | 1.49E-06 | 2.20E-04 |
| ENSMUSG00000031383 | Dusp9 | -4.097 | 4.34E-04 | 1.39E-02 |
| ENSMUSG00000007655 | Cav1 | -4.089 | 2.34E-11 | 2.83E-08 |
| ENSMUSG00000023905 | Tnfrsf12a | -4.012 | 8.10E-13 | 1.61E-09 |
| ENSMUSG00000115009 | G930009F23 | -3.986 | 2.33E-03 | 4.26E-02 |
| ENSMUSG00000091020 | Gm5828 | -3.964 | 1.37E-03 | 2.98E-02 |
| ENSMUSG00000028128 | F3 | -3.963 | 1.25E-07 | 3.42E-05 |
| ENSMUSG00000112134 | Gm40723 | -3.924 | 1.20E-03 | 2.73E-02 |
| ENSMUSG00000005148 | Klf5 | -3.920 | 9.95E-05 | 5.05E-03 |
| ENSMUSG00000066270 | Gm10157 | -3.906 | 2.38E-03 | 4.33E-02 |
| ENSMUSG00000029752 | Asns | -3.883 | 5.62E-12 | 8.75E-09 |
| ENSMUSG00000055407 | Map6 | -3.855 | 5.77E-04 | 1.70E-02 |
| ENSMUSG00000056031 | 9330154J02I | -3.850 | 1.92E-03 | 3.74E-02 |
| ENSMUSG00000028776 | Tinagl1 | -3.826 | 1.91E-10 | 1.90E-07 |
| ENSMUSG00000026479 | Lamc2 | -3.822 | 1.33E-06 | 2.05E-04 |
| ENSMUSG00000115243 | Gm5207 | -3.808 | 4.43E-06 | 5.17E-04 |
| ENSMUSG00000024640 | Psat1 | -3.741 | 2.77E-11 | 3.14E-08 |
| ENSMUSG00000042734 | Ttc9 | -3.738 | 2.08E-03 | 3.95E-02 |
| ENSMUSG00000039126 | Prune2 | -3.732 | 1.76E-09 | 1.07E-06 |
| ENSMUSG00000028480 | Glipr2 | -3.722 | 9.05E-04 | 2.30E-02 |
| ENSMUSG00000019960 | Dusp6 | -3.681 | 4.24E-08 | 1.40E-05 |
| ENSMUSG00000055301 | Adh7 | -3.667 | 9.77E-04 | 2.39E-02 |
| ENSMUSG00000022686 | B3gnt5 | -3.635 | 2.27E-03 | 4.20E-02 |
| ENSMUSG00000019997 | Ctgf | -3.626 | 4.36E-07 | 9.14E-05 |
| ENSMUSG00000074743 | Thbd | -3.612 | 2.22E-03 | 4.13E-02 |
| ENSMUSG00000020256 | Aldh1l2 | -3.611 | 6.19E-05 | 3.56E-03 |
| ENSMUSG00000035799 | Twist1 | -3.592 | 1.74E-03 | 3.51E-02 |
| ENSMUSG00000039462 | Col10a1 | -3.568 | 2.77E-03 | 4.78E-02 |
| ENSMUSG00000073274 | Gm14636 | -3.568 | 3.98E-04 | 1.31E-02 |
| ENSMUSG00000027313 | Chac1 | -3.549 | 1.30E-06 | 2.01E-04 |
| ENSMUSG00000023046 | Igfbp6 | -3.470 | 1.72E-14 | 1.25E-10 |
| ENSMUSG00000029334 | Prkg2 | -3.469 | 2.16E-08 | 8.56E-06 |
| ENSMUSG00000032715 | Trib3 | -3.469 | 6.53E-06 | 6.75E-04 |
| ENSMUSG00000031530 | Dusp4 | -3.436 | 1.49E-04 | 6.78E-03 |
| ENSMUSG00000078249 | Hmga1b | -3.402 | 3.88E-07 | 8.55E-05 |
| ENSMUSG00000047139 | Cd24a | -3.398 | 8.73E-04 | 2.25E-02 |
| ENSMUSG00000000058 | Cav2 | -3.371 | 6.55E-09 | 3.48E-06 |
| ENSMUSG00000024990 | Rbp4 | -3.362 | 2.14E-04 | 8.70E-03 |
| ENSMUSG00000031367 | Ap1s2 | -3.305 | 3.80E-04 | 1.29E-02 |
| ENSMUSG00000037946 | Fgd3 | -3.253 | 7.31E-04 | 2.00E-02 |
| ENSMUSG00000051111 | Sv2c | -3.197 | 5.29E-04 | 1.60E-02 |

| | | | | |
|---|---|---|---|---|
| ENSMUSG00000048482 | Bdnf | -3.153 | 1.40E-03 | 3.00E-02 |
| ENSMUSG00000027611 | Procr | -3.138 | 2.89E-03 | 4.92E-02 |
| ENSMUSG00000024907 | Gal | -3.126 | 1.29E-04 | 6.07E-03 |
| ENSMUSG00000028195 | Cyr61 | -3.082 | 2.31E-07 | 5.74E-05 |
| ENSMUSG00000024883 | Rin1 | -3.062 | 3.92E-04 | 1.31E-02 |
| ENSMUSG00000031355 | Arhgap6 | -3.049 | 1.69E-03 | 3.44E-02 |
| ENSMUSG00000032501 | Trib1 | -3.041 | 2.30E-03 | 4.22E-02 |
| ENSMUSG00000024587 | Nars | -3.035 | 1.14E-09 | 7.33E-07 |
| ENSMUSG00000017144 | Rnd3 | -3.033 | 2.19E-06 | 2.95E-04 |
| ENSMUSG00000021835 | Bmp4 | -3.032 | 1.61E-07 | 4.23E-05 |
| ENSMUSG00000023043 | Krt18 | -3.018 | 3.02E-05 | 2.10E-03 |
| ENSMUSG00000026547 | Tagln2 | -2.996 | 2.49E-04 | 9.55E-03 |
| ENSMUSG00000029161 | Cgref1 | -2.984 | 7.55E-06 | 7.36E-04 |
| ENSMUSG00000021822 | Plau | -2.971 | 3.99E-04 | 1.31E-02 |
| ENSMUSG00000037855 | Zfp365 | -2.949 | 2.70E-05 | 1.95E-03 |
| ENSMUSG00000055737 | Ghr | -2.937 | 2.26E-08 | 8.80E-06 |
| ENSMUSG00000050222 | Il17d | -2.937 | 5.58E-06 | 6.12E-04 |
| ENSMUSG00000022241 | Tars | -2.931 | 2.06E-08 | 8.49E-06 |
| ENSMUSG00000034271 | Jdp2 | -2.927 | 2.15E-05 | 1.65E-03 |
| ENSMUSG00000046711 | Hmga1 | -2.907 | 2.97E-05 | 2.09E-03 |
| ENSMUSG00000005124 | Wisp1 | -2.906 | 2.58E-05 | 1.88E-03 |
| ENSMUSG00000022180 | Slc7a8 | -2.901 | 2.48E-03 | 4.44E-02 |
| ENSMUSG00000031574 | Star | -2.898 | 1.52E-07 | 4.10E-05 |
| ENSMUSG00000047821 | Trim16 | -2.894 | 2.81E-03 | 4.82E-02 |
| ENSMUSG00000025140 | Pycr1 | -2.867 | 9.24E-04 | 2.31E-02 |
| ENSMUSG00000028680 | Plk3 | -2.848 | 2.49E-05 | 1.83E-03 |
| ENSMUSG00000028811 | Yars | -2.841 | 2.39E-07 | 5.80E-05 |
| ENSMUSG00000089812 | Gm15867 | -2.839 | 1.29E-03 | 2.87E-02 |
| ENSMUSG00000038418 | Egr1 | -2.829 | 1.31E-03 | 2.90E-02 |
| ENSMUSG00000026204 | Ptprn | -2.818 | 4.24E-04 | 1.37E-02 |
| ENSMUSG00000083512 | Gm12749 | -2.816 | 2.93E-04 | 1.08E-02 |
| ENSMUSG00000019970 | Sgk1 | -2.812 | 3.57E-05 | 2.42E-03 |
| ENSMUSG00000101431 | Gm7901 | -2.798 | 2.82E-05 | 2.02E-03 |
| ENSMUSG00000029446 | Psph | -2.792 | 1.43E-14 | 1.25E-10 |
| ENSMUSG00000062661 | Ncs1 | -2.776 | 4.53E-06 | 5.23E-04 |
| ENSMUSG00000020312 | Shc2 | -2.746 | 1.07E-03 | 2.53E-02 |
| ENSMUSG00000072944 | Nup62cl | -2.735 | 3.66E-05 | 2.47E-03 |
| ENSMUSG00000073016 | Uprt | -2.729 | 2.75E-03 | 4.76E-02 |
| ENSMUSG00000106390 | Gm5551 | -2.724 | 7.74E-05 | 4.17E-03 |
| ENSMUSG00000081729 | Hspa9-ps1 | -2.694 | 3.61E-04 | 1.24E-02 |
| ENSMUSG00000052837 | Junb | -2.671 | 2.55E-03 | 4.53E-02 |
| ENSMUSG00000025403 | Shmt2 | -2.670 | 7.08E-09 | 3.68E-06 |
| ENSMUSG00000031434 | Morc4 | -2.652 | 1.02E-04 | 5.14E-03 |
| ENSMUSG00000086711 | Gm15482 | -2.645 | 2.81E-03 | 4.82E-02 |
| ENSMUSG00000096740 | Lbhd1 | -2.642 | 1.23E-03 | 2.77E-02 |
| ENSMUSG00000006442 | Srm | -2.637 | 7.18E-08 | 2.18E-05 |
| ENSMUSG00000081819 | Gm12722 | -2.625 | 2.36E-03 | 4.29E-02 |

| | | | | |
|---|---|---|---|---|
| ENSMUSG00000010755 | Cars | -2.619 | 9.92E-12 | 1.44E-08 |
| ENSMUSG00000029777 | Gars | -2.618 | 7.06E-10 | 4.97E-07 |
| ENSMUSG00000005667 | Mthfd2 | -2.612 | 1.16E-07 | 3.25E-05 |
| ENSMUSG00000026749 | Nek6 | -2.606 | 6.52E-07 | 1.22E-04 |
| ENSMUSG00000037960 | Card19 | -2.603 | 3.89E-08 | 1.35E-05 |
| ENSMUSG00000041324 | Inhba | -2.599 | 8.90E-04 | 2.28E-02 |
| ENSMUSG00000032802 | Srxn1 | -2.598 | 4.48E-05 | 2.81E-03 |
| ENSMUSG00000020325 | Fstl3 | -2.583 | 2.39E-03 | 4.34E-02 |
| ENSMUSG00000057604 | Lmcd1 | -2.573 | 2.93E-04 | 1.08E-02 |
| ENSMUSG00000034839 | Larp6 | -2.561 | 2.44E-03 | 4.40E-02 |
| ENSMUSG00000003541 | Ier3 | -2.560 | 5.09E-06 | 5.75E-04 |
| ENSMUSG00000037060 | Cavin3 | -2.554 | 1.39E-03 | 2.99E-02 |
| ENSMUSG00000043336 | Filip1l | -2.540 | 5.70E-04 | 1.69E-02 |
| ENSMUSG00000083465 | Gm11652 | -2.517 | 9.80E-06 | 8.88E-04 |
| ENSMUSG00000003348 | Mob3a | -2.515 | 5.10E-05 | 3.10E-03 |
| ENSMUSG00000097180 | 2700038G22 | -2.500 | 2.38E-08 | 9.12E-06 |
| ENSMUSG00000022557 | Bop1 | -2.457 | 1.91E-06 | 2.70E-04 |
| ENSMUSG00000037465 | Klf10 | -2.445 | 1.04E-05 | 9.32E-04 |
| ENSMUSG00000113769 | 5033406O09 | -2.437 | 8.99E-05 | 4.74E-03 |
| ENSMUSG00000030717 | Nupr1 | -2.433 | 2.74E-03 | 4.74E-02 |
| ENSMUSG00000060950 | Trmt61a | -2.430 | 9.29E-05 | 4.81E-03 |
| ENSMUSG00000097000 | Gm17435 | -2.405 | 6.51E-04 | 1.85E-02 |
| ENSMUSG00000039419 | Cntnap2 | -2.388 | 2.49E-06 | 3.24E-04 |
| ENSMUSG00000040322 | Slc25a24 | -2.385 | 3.25E-06 | 4.03E-04 |
| ENSMUSG00000029096 | Htra3 | -2.383 | 1.36E-03 | 2.96E-02 |
| ENSMUSG00000025007 | Aldh18a1 | -2.373 | 6.90E-07 | 1.27E-04 |
| ENSMUSG00000090394 | 4930523C07 | -2.367 | 9.28E-05 | 4.81E-03 |
| ENSMUSG00000030222 | Rerg | -2.362 | 1.04E-04 | 5.19E-03 |
| ENSMUSG00000038335 | Tsr1 | -2.345 | 4.71E-13 | 1.17E-09 |
| ENSMUSG00000038279 | Nop2 | -2.334 | 3.23E-10 | 2.61E-07 |
| ENSMUSG00000050017 | Pitpnb | -2.321 | 2.20E-05 | 1.68E-03 |
| ENSMUSG00000102048 | Gm6075 | -2.315 | 1.83E-04 | 7.80E-03 |
| ENSMUSG00000053398 | Phgdh | -2.313 | 5.60E-07 | 1.09E-04 |
| ENSMUSG00000028069 | Gpatch4 | -2.305 | 5.12E-09 | 2.80E-06 |
| ENSMUSG00000028063 | Lmna | -2.295 | 9.13E-05 | 4.78E-03 |
| ENSMUSG00000025287 | Acot9 | -2.273 | 2.03E-07 | 5.14E-05 |
| ENSMUSG00000022114 | Spry2 | -2.266 | 5.19E-04 | 1.58E-02 |
| ENSMUSG00000045763 | Basp1 | -2.247 | 8.51E-06 | 8.01E-04 |
| ENSMUSG00000068739 | Sars | -2.239 | 1.02E-06 | 1.70E-04 |
| ENSMUSG00000104560 | Gm8115 | -2.237 | 6.56E-04 | 1.86E-02 |
| ENSMUSG00000085666 | Gm9855 | -2.234 | 3.56E-04 | 1.22E-02 |
| ENSMUSG00000026784 | Pdss1 | -2.234 | 8.56E-10 | 5.83E-07 |
| ENSMUSG00000000561 | Wdr77 | -2.224 | 4.22E-07 | 9.04E-05 |
| ENSMUSG00000041506 | Rrp9 | -2.223 | 2.39E-07 | 5.80E-05 |
| ENSMUSG00000032051 | Fdx1 | -2.214 | 7.57E-05 | 4.11E-03 |
| ENSMUSG00000051223 | Bzw1 | -2.203 | 2.59E-07 | 6.00E-05 |
| ENSMUSG00000027610 | Gss | -2.196 | 6.57E-07 | 1.22E-04 |

| | | | | |
|---|---|---|---|---|
| ENSMUSG00000024493 | Lars | -2.194 | 2.88E-11 | 3.14E-08 |
| ENSMUSG00000024037 | Wdr4 | -2.173 | 5.63E-06 | 6.15E-04 |
| ENSMUSG00000028896 | Rcc1 | -2.162 | 2.53E-06 | 3.27E-04 |
| ENSMUSG00000024381 | Bin1 | -2.161 | 1.60E-04 | 7.15E-03 |
| ENSMUSG00000053746 | Ptrh1 | -2.160 | 3.95E-05 | 2.59E-03 |
| ENSMUSG00000070167 | Snora57 | -2.159 | 4.00E-04 | 1.31E-02 |
| ENSMUSG00000013089 | Etv5 | -2.152 | 9.96E-05 | 5.05E-03 |
| ENSMUSG00000032265 | Fam46a | -2.137 | 1.08E-03 | 2.54E-02 |
| ENSMUSG00000056501 | Cebpb | -2.137 | 5.17E-04 | 1.58E-02 |
| ENSMUSG00000010067 | Rassf1 | -2.136 | 6.87E-05 | 3.84E-03 |
| ENSMUSG00000020303 | Stc2 | -2.129 | 1.78E-05 | 1.44E-03 |
| ENSMUSG00000010048 | Ifrd2 | -2.122 | 4.77E-04 | 1.49E-02 |
| ENSMUSG00000025511 | Tspan4 | -2.113 | 1.44E-06 | 2.17E-04 |
| ENSMUSG00000032185 | Carm1 | -2.105 | 1.11E-03 | 2.57E-02 |
| ENSMUSG00000042608 | Stk40 | -2.102 | 2.97E-04 | 1.09E-02 |
| ENSMUSG00000031700 | Gpt2 | -2.091 | 1.05E-04 | 5.19E-03 |
| ENSMUSG00000042515 | Mum1l1 | -2.089 | 1.11E-06 | 1.81E-04 |
| ENSMUSG00000025132 | Arhgdia | -2.060 | 1.36E-03 | 2.96E-02 |
| ENSMUSG00000053560 | Ier2 | -2.058 | 7.60E-04 | 2.04E-02 |
| ENSMUSG00000081302 | Gm12020 | -2.042 | 5.05E-04 | 1.55E-02 |
| ENSMUSG00000025171 | Ubtd1 | -2.027 | 5.99E-04 | 1.75E-02 |
| ENSMUSG00000091898 | Tnnc1 | -2.008 | 6.10E-05 | 3.53E-03 |
| ENSMUSG00000029171 | Pgm1 | -2.002 | 8.35E-05 | 4.45E-03 |
| ENSMUSG00000041360 | Pum3 | -1.997 | 4.71E-06 | 5.35E-04 |
| ENSMUSG00000000916 | Nsun5 | -1.991 | 6.65E-06 | 6.85E-04 |
| ENSMUSG00000029599 | Ddx54 | -1.989 | 9.43E-04 | 2.34E-02 |
| ENSMUSG00000038552 | Fndc4 | -1.975 | 2.79E-03 | 4.79E-02 |
| ENSMUSG00000028010 | Gar1 | -1.968 | 1.16E-08 | 5.76E-06 |
| ENSMUSG00000032966 | Fkbp1a | -1.967 | 2.07E-05 | 1.60E-03 |
| ENSMUSG00000110126 | Gm9347 | -1.956 | 1.72E-03 | 3.47E-02 |
| ENSMUSG00000063229 | Ldha | -1.937 | 6.90E-05 | 3.85E-03 |
| ENSMUSG00000038894 | Irs2 | -1.937 | 4.07E-04 | 1.33E-02 |
| ENSMUSG00000024312 | Wdr46 | -1.932 | 4.67E-05 | 2.90E-03 |
| ENSMUSG00000029033 | Acap3 | -1.927 | 8.31E-04 | 2.19E-02 |
| ENSMUSG00000031490 | Eif4ebp1 | -1.923 | 2.85E-04 | 1.06E-02 |
| ENSMUSG00000078862 | Gm14326 | -1.920 | 1.38E-03 | 2.98E-02 |
| ENSMUSG00000023988 | Bysl | -1.913 | 5.44E-06 | 6.01E-04 |
| ENSMUSG00000015312 | Gadd45b | -1.910 | 2.37E-04 | 9.19E-03 |
| ENSMUSG00000040354 | Mars | -1.909 | 1.16E-06 | 1.84E-04 |
| ENSMUSG00000024772 | Ehd1 | -1.908 | 3.96E-04 | 1.31E-02 |
| ENSMUSG00000041774 | Ydjc | -1.904 | 4.45E-04 | 1.41E-02 |
| ENSMUSG00000002343 | Armc6 | -1.896 | 1.45E-03 | 3.07E-02 |
| ENSMUSG00000003500 | Impdh1 | -1.893 | 1.74E-05 | 1.43E-03 |
| ENSMUSG00000039405 | Prss23 | -1.879 | 1.10E-03 | 2.57E-02 |
| ENSMUSG00000095567 | Noc2l | -1.876 | 3.51E-06 | 4.31E-04 |
| ENSMUSG00000053801 | Grwd1 | -1.873 | 8.40E-07 | 1.43E-04 |
| ENSMUSG00000011179 | Odc1 | -1.869 | 2.89E-07 | 6.63E-05 |

| | | | | |
|---|---|---|---|---|
| ENSMUSG00000060477 | Irak2 | -1.868 | 8.88E-04 | 2.28E-02 |
| ENSMUSG00000026637 | Traf5 | -1.855 | 1.17E-03 | 2.68E-02 |
| ENSMUSG00000027405 | Nop56 | -1.852 | 1.49E-08 | 6.79E-06 |
| ENSMUSG00000048261 | Gm4879 | -1.848 | 1.07E-04 | 5.29E-03 |
| ENSMUSG00000001305 | Rrp15 | -1.838 | 3.05E-06 | 3.85E-04 |
| ENSMUSG00000018932 | Map2k3 | -1.835 | 4.47E-04 | 1.42E-02 |
| ENSMUSG00000025855 | Prkar1b | -1.835 | 2.37E-04 | 9.19E-03 |
| ENSMUSG00000047565 | Acot10 | -1.835 | 1.04E-03 | 2.48E-02 |
| ENSMUSG00000033706 | Smyd5 | -1.834 | 1.95E-05 | 1.52E-03 |
| ENSMUSG00000025485 | Ric8a | -1.828 | 3.89E-05 | 2.57E-03 |
| ENSMUSG00000025869 | Nop16 | -1.821 | 2.30E-05 | 1.74E-03 |
| ENSMUSG00000046865 | Fbl | -1.816 | 1.43E-05 | 1.20E-03 |
| ENSMUSG00000055044 | Pdlim1 | -1.815 | 1.91E-03 | 3.73E-02 |
| ENSMUSG00000003848 | Nob1 | -1.813 | 6.79E-05 | 3.83E-03 |
| ENSMUSG00000020205 | Phlda1 | -1.808 | 7.98E-06 | 7.64E-04 |
| ENSMUSG00000034765 | Dusp5 | -1.808 | 1.92E-04 | 8.06E-03 |
| ENSMUSG00000021595 | Nsun2 | -1.803 | 1.35E-05 | 1.16E-03 |
| ENSMUSG00000074280 | Gm6166 | -1.801 | 5.65E-04 | 1.68E-02 |
| ENSMUSG00000028495 | Rps6 | -1.790 | 4.00E-05 | 2.60E-03 |
| ENSMUSG00000026019 | Wdr12 | -1.790 | 2.48E-06 | 3.24E-04 |
| ENSMUSG00000085385 | Snhg17 | -1.789 | 1.18E-06 | 1.87E-04 |
| ENSMUSG00000038539 | Atf5 | -1.776 | 3.44E-04 | 1.20E-02 |
| ENSMUSG00000037601 | Nme1 | -1.774 | 5.45E-06 | 6.01E-04 |
| ENSMUSG00000042354 | Gnl3 | -1.772 | 5.75E-06 | 6.22E-04 |
| ENSMUSG00000026234 | Ncl | -1.772 | 6.31E-10 | 4.59E-07 |
| ENSMUSG00000058355 | Abce1 | -1.768 | 7.21E-06 | 7.26E-04 |
| ENSMUSG00000062937 | Mtap | -1.767 | 8.98E-06 | 8.30E-04 |
| ENSMUSG00000081732 | Gm15495 | -1.765 | 5.32E-04 | 1.61E-02 |
| ENSMUSG00000060647 | Gm7099 | -1.765 | 1.22E-03 | 2.75E-02 |
| ENSMUSG00000042406 | Atf4 | -1.751 | 2.38E-05 | 1.79E-03 |
| ENSMUSG00000001131 | Timp1 | -1.748 | 2.06E-04 | 8.51E-03 |
| ENSMUSG00000059325 | Hopx | -1.747 | 1.27E-03 | 2.82E-02 |
| ENSMUSG00000022369 | Mtbp | -1.732 | 2.42E-04 | 9.32E-03 |
| ENSMUSG00000029364 | Wsb2 | -1.723 | 8.72E-04 | 2.25E-02 |
| ENSMUSG00000057541 | Pus7 | -1.720 | 1.35E-05 | 1.16E-03 |
| ENSMUSG00000106664 | Gm17936 | -1.719 | 2.23E-04 | 8.89E-03 |
| ENSMUSG00000031352 | Hccs | -1.718 | 3.74E-06 | 4.48E-04 |
| ENSMUSG00000057561 | Eif1a | -1.715 | 5.38E-07 | 1.06E-04 |
| ENSMUSG00000028670 | Lypla2 | -1.713 | 6.78E-04 | 1.90E-02 |
| ENSMUSG00000024190 | Dusp1 | -1.710 | 1.35E-03 | 2.96E-02 |
| ENSMUSG00000083477 | Gm5555 | -1.706 | 6.16E-04 | 1.78E-02 |
| ENSMUSG00000081999 | Gm13461 | -1.703 | 6.59E-07 | 1.22E-04 |
| ENSMUSG00000078789 | Dph1 | -1.700 | 9.95E-05 | 5.05E-03 |
| ENSMUSG00000097769 | Snhg4 | -1.694 | 2.54E-04 | 9.70E-03 |
| ENSMUSG00000021196 | Pfkp | -1.693 | 1.77E-04 | 7.66E-03 |
| ENSMUSG00000005103 | Wdr1 | -1.692 | 4.66E-04 | 1.46E-02 |
| ENSMUSG00000055612 | Cdca7 | -1.690 | 8.72E-07 | 1.46E-04 |

| | | | | |
|---|---|---|---|---|
| ENSMUSG00000070284 | Gmppb | -1.686 | 1.36E-04 | 6.33E-03 |
| ENSMUSG00000032231 | Anxa2 | -1.682 | 9.22E-05 | 4.80E-03 |
| ENSMUSG00000052688 | Rab7b | -1.681 | 5.67E-04 | 1.68E-02 |
| ENSMUSG00000024999 | Noc3l | -1.679 | 1.08E-04 | 5.30E-03 |
| ENSMUSG00000008384 | Sertad1 | -1.679 | 1.33E-04 | 6.22E-03 |
| ENSMUSG00000020547 | Bzw2 | -1.677 | 7.32E-04 | 2.00E-02 |
| ENSMUSG00000048612 | Myof | -1.673 | 8.47E-04 | 2.22E-02 |
| ENSMUSG00000052151 | Plpp2 | -1.673 | 1.01E-05 | 9.09E-04 |
| ENSMUSG00000051557 | Pusl1 | -1.668 | 3.42E-04 | 1.20E-02 |
| ENSMUSG00000058809 | Hspd1-ps3 | -1.649 | 4.23E-05 | 2.71E-03 |
| ENSMUSG00000043192 | Gm1840 | -1.641 | 1.16E-03 | 2.67E-02 |
| ENSMUSG00000024841 | Eif1ad | -1.636 | 2.53E-07 | 5.98E-05 |
| ENSMUSG00000040463 | Mybbp1a | -1.625 | 1.73E-08 | 7.69E-06 |
| ENSMUSG00000030609 | Aen | -1.617 | 1.26E-03 | 2.81E-02 |
| ENSMUSG00000078812 | Eif5a | -1.616 | 2.90E-04 | 1.07E-02 |
| ENSMUSG00000048285 | Frmd6 | -1.616 | 4.78E-04 | 1.49E-02 |
| ENSMUSG00000031578 | Mak16 | -1.614 | 1.35E-06 | 2.07E-04 |
| ENSMUSG00000032078 | Zpr1 | -1.614 | 1.62E-04 | 7.22E-03 |
| ENSMUSG00000031960 | Aars | -1.611 | 1.86E-05 | 1.47E-03 |
| ENSMUSG00000006498 | Ptbp1 | -1.611 | 2.63E-03 | 4.62E-02 |
| ENSMUSG00000063316 | Rpl27 | -1.609 | 5.29E-07 | 1.05E-04 |
| ENSMUSG00000057614 | Gnai1 | -1.609 | 5.67E-04 | 1.68E-02 |
| ENSMUSG00000028729 | Ebna1bp2 | -1.607 | 1.20E-07 | 3.31E-05 |
| ENSMUSG00000025364 | Pa2g4 | -1.606 | 1.16E-03 | 2.66E-02 |
| ENSMUSG00000040675 | Mthfd1l | -1.603 | 1.66E-09 | 1.03E-06 |
| ENSMUSG00000026020 | Nop58 | -1.597 | 3.09E-10 | 2.59E-07 |
| ENSMUSG00000061024 | Rrs1 | -1.595 | 7.41E-07 | 1.34E-04 |
| ENSMUSG00000112825 | Gm9118 | -1.594 | 1.99E-03 | 3.83E-02 |
| ENSMUSG00000024697 | Gna14 | -1.591 | 5.14E-04 | 1.58E-02 |
| ENSMUSG00000026617 | Bpnt1 | -1.591 | 2.04E-08 | 8.49E-06 |
| ENSMUSG00000019082 | Slc25a22 | -1.590 | 5.38E-06 | 5.99E-04 |
| ENSMUSG00000043424 | Eif3j2 | -1.588 | 2.76E-10 | 2.48E-07 |
| ENSMUSG00000026305 | Lrrfip1 | -1.588 | 1.58E-03 | 3.26E-02 |
| ENSMUSG00000031657 | Heatr3 | -1.587 | 1.50E-06 | 2.20E-04 |
| ENSMUSG00000031568 | Rwdd4a | -1.579 | 6.51E-06 | 6.75E-04 |
| ENSMUSG00000020116 | Pno1 | -1.579 | 4.16E-08 | 1.40E-05 |
| ENSMUSG00000003355 | Fkbp11 | -1.574 | 1.58E-03 | 3.27E-02 |
| ENSMUSG00000006732 | Mettl1 | -1.572 | 2.19E-06 | 2.95E-04 |
| ENSMUSG00000019849 | Prep | -1.571 | 5.01E-05 | 3.06E-03 |
| ENSMUSG00000024785 | Rcl1 | -1.566 | 7.90E-04 | 2.11E-02 |
| ENSMUSG00000002825 | Qtrt1 | -1.564 | 9.70E-04 | 2.37E-02 |
| ENSMUSG00000029447 | Cct6a | -1.559 | 2.27E-06 | 3.04E-04 |
| ENSMUSG00000048007 | Timm8a1 | -1.557 | 2.31E-04 | 9.08E-03 |
| ENSMUSG00000029610 | Aimp2 | -1.555 | 3.68E-06 | 4.46E-04 |
| ENSMUSG00000022148 | Fyb | -1.552 | 2.10E-03 | 3.98E-02 |
| ENSMUSG00000020075 | Ddx21 | -1.547 | 4.26E-10 | 3.32E-07 |
| ENSMUSG00000037295 | Ldlrap1 | -1.541 | 5.39E-04 | 1.63E-02 |

| | | | | |
|---|---|---|---|---|
| ENSMUSG00000024359 | Hspa9 | -1.540 | 1.14E-09 | 7.33E-07 |
| ENSMUSG00000008450 | Nutf2 | -1.539 | 7.36E-06 | 7.30E-04 |
| ENSMUSG00000049957 | Ccdc137 | -1.539 | 4.62E-04 | 1.46E-02 |
| ENSMUSG00000073775 | Kti12 | -1.538 | 4.44E-04 | 1.41E-02 |
| ENSMUSG00000020869 | Lrrc59 | -1.538 | 1.25E-04 | 5.92E-03 |
| ENSMUSG00000026509 | Capn2 | -1.538 | 1.69E-03 | 3.44E-02 |
| ENSMUSG00000079614 | Seh1l | -1.533 | 4.87E-04 | 1.51E-02 |
| ENSMUSG00000062014 | Gmfb | -1.527 | 3.26E-04 | 1.17E-02 |
| ENSMUSG00000031792 | Usb1 | -1.521 | 1.15E-03 | 2.65E-02 |
| ENSMUSG00000021288 | Klc1 | -1.519 | 3.18E-04 | 1.15E-02 |
| ENSMUSG00000082121 | Gm8199 | -1.519 | 1.57E-03 | 3.25E-02 |
| ENSMUSG00000066839 | Ecsit | -1.516 | 7.21E-04 | 1.98E-02 |
| ENSMUSG00000105987 | AI506816 | -1.516 | 2.33E-03 | 4.26E-02 |
| ENSMUSG00000106943 | Dancr | -1.514 | 1.82E-03 | 3.62E-02 |
| ENSMUSG00000018446 | C1qbp | -1.513 | 3.73E-08 | 1.34E-05 |
| ENSMUSG00000030079 | Ruvbl1 | -1.513 | 7.53E-06 | 7.36E-04 |
| ENSMUSG00000027804 | Ppid | -1.504 | 2.11E-06 | 2.92E-04 |
| ENSMUSG00000018433 | Nol11 | -1.504 | 3.65E-04 | 1.24E-02 |
| ENSMUSG00000114419 | 5430414B19 | -1.496 | 8.93E-04 | 2.28E-02 |
| ENSMUSG00000056076 | Eif3b | -1.487 | 2.44E-04 | 9.37E-03 |
| ENSMUSG00000015176 | Nolc1 | -1.487 | 2.61E-05 | 1.90E-03 |
| ENSMUSG00000039910 | Cited2 | -1.484 | 2.75E-04 | 1.03E-02 |
| ENSMUSG00000071359 | Tbpl1 | -1.480 | 1.42E-04 | 6.53E-03 |
| ENSMUSG00000028179 | Cth | -1.480 | 2.29E-03 | 4.22E-02 |
| ENSMUSG00000026245 | Farsb | -1.480 | 3.00E-04 | 1.09E-02 |
| ENSMUSG00000005481 | Ddx39 | -1.476 | 2.44E-05 | 1.80E-03 |
| ENSMUSG00000028741 | Mrto4 | -1.475 | 1.61E-06 | 2.33E-04 |
| ENSMUSG00000039682 | Lap3 | -1.474 | 5.44E-05 | 3.26E-03 |
| ENSMUSG00000063888 | Rpl7l1 | -1.473 | 1.45E-04 | 6.63E-03 |
| ENSMUSG00000027533 | Fabp5 | -1.473 | 2.71E-03 | 4.70E-02 |
| ENSMUSG00000029507 | Pus1 | -1.472 | 1.14E-04 | 5.47E-03 |
| ENSMUSG00000001627 | Ifrd1 | -1.470 | 1.10E-04 | 5.33E-03 |
| ENSMUSG00000028318 | Polr1e | -1.463 | 7.61E-05 | 4.12E-03 |
| ENSMUSG00000014873 | Surf2 | -1.463 | 1.34E-03 | 2.94E-02 |
| ENSMUSG00000021877 | Arf4 | -1.461 | 1.28E-03 | 2.85E-02 |
| ENSMUSG00000024535 | Snx24 | -1.461 | 2.85E-03 | 4.87E-02 |
| ENSMUSG00000027808 | Serp1 | -1.459 | 3.10E-06 | 3.89E-04 |
| ENSMUSG00000029500 | Pgam5 | -1.453 | 3.83E-04 | 1.29E-02 |
| ENSMUSG00000074656 | Eif2s2 | -1.450 | 2.14E-06 | 2.93E-04 |
| ENSMUSG00000021149 | Gtpbp4 | -1.448 | 4.78E-09 | 2.68E-06 |
| ENSMUSG00000026615 | Eprs | -1.448 | 1.63E-07 | 4.23E-05 |
| ENSMUSG00000000078 | Klf6 | -1.445 | 6.40E-04 | 1.83E-02 |
| ENSMUSG00000024805 | Pcgf5 | -1.444 | 2.30E-05 | 1.74E-03 |
| ENSMUSG00000038510 | Rpf2 | -1.442 | 1.66E-04 | 7.32E-03 |
| ENSMUSG00000031917 | Nip7 | -1.441 | 1.61E-07 | 4.23E-05 |
| ENSMUSG00000030432 | Rpl28 | -1.429 | 6.39E-05 | 3.65E-03 |
| ENSMUSG00000030738 | Eif3c | -1.425 | 2.89E-06 | 3.66E-04 |

| | | | | |
|---|---|---|---|---|
| ENSMUSG00000030268 | Bcat1 | -1.424 | 9.37E-04 | 2.33E-02 |
| ENSMUSG00000026192 | Atic | -1.423 | 2.19E-04 | 8.81E-03 |
| ENSMUSG00000025980 | Hspd1 | -1.418 | 1.55E-12 | 2.81E-09 |
| ENSMUSG00000028180 | Zranb2 | -1.411 | 6.10E-06 | 6.44E-04 |
| ENSMUSG00000025981 | Coq10b | -1.410 | 1.40E-04 | 6.47E-03 |
| ENSMUSG00000028683 | Eif2b3 | -1.406 | 7.59E-07 | 1.35E-04 |
| ENSMUSG00000030189 | Ybx3 | -1.401 | 8.57E-04 | 2.23E-02 |
| ENSMUSG00000056131 | Pgm3 | -1.400 | 1.38E-03 | 2.98E-02 |
| ENSMUSG00000026670 | Uap1 | -1.399 | 9.33E-06 | 8.52E-04 |
| ENSMUSG00000032435 | Dync1li1 | -1.392 | 1.24E-03 | 2.79E-02 |
| ENSMUSG00000019814 | Ltv1 | -1.392 | 8.11E-05 | 4.34E-03 |
| ENSMUSG00000064264 | Zfp428 | -1.389 | 3.90E-06 | 4.65E-04 |
| ENSMUSG00000018848 | Rars | -1.388 | 5.39E-05 | 3.24E-03 |
| ENSMUSG00000024360 | Etf1 | -1.387 | 8.72E-04 | 2.25E-02 |
| ENSMUSG00000040688 | Tbl3 | -1.387 | 4.65E-04 | 1.46E-02 |
| ENSMUSG00000038736 | Nudcd1 | -1.387 | 3.03E-05 | 2.10E-03 |
| ENSMUSG00000028953 | Abcf2 | -1.386 | 6.73E-05 | 3.82E-03 |
| ENSMUSG00000067367 | Lyar | -1.384 | 1.78E-05 | 1.44E-03 |
| ENSMUSG00000068922 | Msto1 | -1.384 | 6.33E-04 | 1.82E-02 |
| ENSMUSG00000038388 | Mpp6 | -1.379 | 1.38E-03 | 2.98E-02 |
| ENSMUSG00000016554 | Eif3d | -1.376 | 4.60E-07 | 9.38E-05 |
| ENSMUSG00000021116 | Eif2s1 | -1.371 | 1.94E-05 | 1.52E-03 |
| ENSMUSG00000001785 | Pwp1 | -1.360 | 3.80E-05 | 2.53E-03 |
| ENSMUSG00000039356 | Exosc2 | -1.360 | 2.07E-04 | 8.53E-03 |
| ENSMUSG00000030759 | Far1 | -1.359 | 4.57E-07 | 9.38E-05 |
| ENSMUSG00000020691 | Mettl2 | -1.358 | 5.97E-04 | 1.75E-02 |
| ENSMUSG00000040028 | Elavl1 | -1.358 | 1.60E-03 | 3.29E-02 |
| ENSMUSG00000003868 | Ruvbl2 | -1.354 | 4.65E-05 | 2.89E-03 |
| ENSMUSG00000027944 | Hax1 | -1.353 | 1.05E-06 | 1.74E-04 |
| ENSMUSG00000057236 | Rbbp4 | -1.352 | 1.23E-03 | 2.78E-02 |
| ENSMUSG00000030983 | Bccip | -1.351 | 2.79E-03 | 4.81E-02 |
| ENSMUSG00000041057 | Wdr43 | -1.350 | 1.32E-05 | 1.15E-03 |
| ENSMUSG00000020780 | Srp68 | -1.348 | 3.66E-04 | 1.24E-02 |
| ENSMUSG00000011114 | Tbrg1 | -1.345 | 1.54E-04 | 6.94E-03 |
| ENSMUSG00000025995 | Wdr75 | -1.344 | 4.16E-05 | 2.69E-03 |
| ENSMUSG00000034343 | Ube2f | -1.343 | 5.38E-08 | 1.65E-05 |
| ENSMUSG00000025591 | Tma16 | -1.342 | 1.91E-06 | 2.70E-04 |
| ENSMUSG00000029430 | Ran | -1.342 | 7.67E-07 | 1.35E-04 |
| ENSMUSG00000027940 | Tpm3 | -1.341 | 9.58E-04 | 2.36E-02 |
| ENSMUSG00000068039 | Tcp1 | -1.338 | 2.10E-08 | 8.50E-06 |
| ENSMUSG00000042215 | Bag2 | -1.336 | 4.20E-05 | 2.70E-03 |
| ENSMUSG00000063802 | Hspbp1 | -1.336 | 1.05E-03 | 2.50E-02 |
| ENSMUSG00000001380 | Hars | -1.334 | 8.53E-04 | 2.23E-02 |
| ENSMUSG00000031432 | Prps1 | -1.333 | 6.97E-04 | 1.93E-02 |
| ENSMUSG00000020358 | Hnrnpab | -1.331 | 1.23E-04 | 5.86E-03 |
| ENSMUSG00000023110 | Prmt5 | -1.329 | 2.56E-04 | 9.74E-03 |
| ENSMUSG00000036427 | Gpi1 | -1.326 | 9.12E-04 | 2.31E-02 |

| | | | | |
|---|---|---|---|---|
| ENSMUSG00000068856 | Sf3b4 | -1.325 | 1.05E-03 | 2.50E-02 |
| ENSMUSG00000025240 | Sacm1l | -1.321 | 7.65E-04 | 2.05E-02 |
| ENSMUSG00000035530 | Eif1 | -1.313 | 3.95E-08 | 1.35E-05 |
| ENSMUSG00000062070 | Pgk1 | -1.311 | 3.98E-04 | 1.31E-02 |
| ENSMUSG00000034120 | Srsf2 | -1.310 | 3.82E-07 | 8.50E-05 |
| ENSMUSG00000026492 | Tfb2m | -1.309 | 2.32E-03 | 4.26E-02 |
| ENSMUSG00000052825 | Gm9892 | -1.309 | 3.26E-04 | 1.17E-02 |
| ENSMUSG00000026356 | Dars | -1.299 | 1.30E-04 | 6.13E-03 |
| ENSMUSG00000029415 | Sdad1 | -1.298 | 9.24E-05 | 4.80E-03 |
| ENSMUSG00000035726 | Supt16 | -1.295 | 1.77E-04 | 7.66E-03 |
| ENSMUSG00000035828 | Pim3 | -1.293 | 1.20E-03 | 2.73E-02 |
| ENSMUSG00000047649 | Cd3eap | -1.292 | 1.88E-05 | 1.48E-03 |
| ENSMUSG00000030888 | Rrp8 | -1.292 | 2.12E-04 | 8.60E-03 |
| ENSMUSG00000036693 | Nop14 | -1.290 | 2.51E-07 | 5.98E-05 |
| ENSMUSG00000078877 | Gm14295 | -1.289 | 3.76E-05 | 2.52E-03 |
| ENSMUSG00000096403 | Gm9825 | -1.288 | 1.07E-03 | 2.52E-02 |
| ENSMUSG00000037376 | Trmt6 | -1.287 | 1.37E-05 | 1.17E-03 |
| ENSMUSG00000053289 | Ddx10 | -1.283 | 1.89E-04 | 7.99E-03 |
| ENSMUSG00000034667 | Xpot | -1.280 | 7.66E-04 | 2.05E-02 |
| ENSMUSG00000081603 | Gm14681 | -1.279 | 4.75E-05 | 2.93E-03 |
| ENSMUSG00000006699 | Cdc42 | -1.279 | 2.50E-04 | 9.58E-03 |
| ENSMUSG00000026520 | Pycr2 | -1.276 | 2.68E-03 | 4.69E-02 |
| ENSMUSG00000027624 | Epb41l1 | -1.276 | 4.29E-04 | 1.38E-02 |
| ENSMUSG00000030057 | Cnbp | -1.274 | 4.28E-05 | 2.71E-03 |
| ENSMUSG00000041488 | Stx3 | -1.271 | 2.98E-05 | 2.09E-03 |
| ENSMUSG00000063785 | Utp14a | -1.269 | 2.54E-03 | 4.51E-02 |
| ENSMUSG00000040472 | Rabggta | -1.268 | 1.92E-04 | 8.06E-03 |
| ENSMUSG00000053907 | Mat2a | -1.263 | 6.29E-14 | 3.43E-10 |
| ENSMUSG00000067150 | Xpo5 | -1.259 | 9.15E-04 | 2.31E-02 |
| ENSMUSG00000020089 | Ppa1 | -1.258 | 1.10E-07 | 3.19E-05 |
| ENSMUSG00000032557 | Uba5 | -1.257 | 5.73E-05 | 3.39E-03 |
| ENSMUSG00000031278 | Acsl4 | -1.257 | 6.83E-05 | 3.83E-03 |
| ENSMUSG00000031485 | Plpbp | -1.256 | 7.61E-04 | 2.04E-02 |
| ENSMUSG00000019487 | Trip10 | -1.255 | 1.02E-03 | 2.45E-02 |
| ENSMUSG00000027236 | Eif3j1 | -1.255 | 6.25E-07 | 1.20E-04 |
| ENSMUSG00000024732 | Ccdc86 | -1.254 | 1.83E-04 | 7.81E-03 |
| ENSMUSG00000000339 | Rtca | -1.252 | 7.39E-04 | 2.01E-02 |
| ENSMUSG00000031167 | Rbm3 | -1.250 | 1.10E-04 | 5.33E-03 |
| ENSMUSG00000016319 | Slc25a5 | -1.248 | 2.51E-05 | 1.84E-03 |
| ENSMUSG00000020925 | Ccdc43 | -1.247 | 7.81E-05 | 4.19E-03 |
| ENSMUSG00000026377 | Nifk | -1.238 | 8.35E-06 | 7.88E-04 |
| ENSMUSG00000022234 | Cct5 | -1.226 | 1.54E-05 | 1.28E-03 |
| ENSMUSG00000022184 | Fbxo4 | -1.224 | 4.14E-04 | 1.35E-02 |
| ENSMUSG00000046027 | Stard5 | -1.223 | 4.40E-04 | 1.41E-02 |
| ENSMUSG00000021282 | Eif5 | -1.223 | 1.41E-05 | 1.19E-03 |
| ENSMUSG00000054000 | Tusc1 | -1.222 | 2.45E-03 | 4.41E-02 |
| ENSMUSG00000036285 | Noa1 | -1.221 | 1.31E-04 | 6.13E-03 |

| | | | | |
|---|---|---|---|---|
| ENSMUSG00000040550 | Otud6b | -1.217 | 1.17E-03 | 2.69E-02 |
| ENSMUSG00000015363 | Trabd | -1.216 | 2.18E-03 | 4.07E-02 |
| ENSMUSG00000038612 | Mcl1 | -1.210 | 4.00E-04 | 1.31E-02 |
| ENSMUSG00000028277 | Ube2j1 | -1.210 | 2.02E-03 | 3.86E-02 |
| ENSMUSG00000017999 | Ddx27 | -1.206 | 7.22E-06 | 7.26E-04 |
| ENSMUSG00000010095 | Slc3a2 | -1.204 | 1.53E-05 | 1.28E-03 |
| ENSMUSG00000039114 | Nrn1 | -1.203 | 2.15E-03 | 4.03E-02 |
| ENSMUSG00000022744 | Cldnd1 | -1.197 | 3.81E-05 | 2.53E-03 |
| ENSMUSG00000001707 | Eef1e1 | -1.197 | 2.34E-05 | 1.77E-03 |
| ENSMUSG00000022407 | Adsl | -1.195 | 5.17E-04 | 1.58E-02 |
| ENSMUSG00000002881 | Nab1 | -1.193 | 7.18E-05 | 3.96E-03 |
| ENSMUSG00000027854 | Sike1 | -1.190 | 1.61E-03 | 3.31E-02 |
| ENSMUSG00000026806 | Ddx31 | -1.190 | 3.29E-04 | 1.17E-02 |
| ENSMUSG00000004127 | Trmt10a | -1.190 | 5.86E-04 | 1.72E-02 |
| ENSMUSG00000025264 | Tsr2 | -1.190 | 1.74E-04 | 7.58E-03 |
| ENSMUSG00000020250 | Txnrd1 | -1.189 | 2.80E-03 | 4.81E-02 |
| ENSMUSG00000063694 | Cycs | -1.189 | 6.59E-04 | 1.86E-02 |
| ENSMUSG00000032042 | Srpr | -1.188 | 1.68E-03 | 3.43E-02 |
| ENSMUSG00000028330 | Ncbp1 | -1.187 | 3.41E-04 | 1.20E-02 |
| ENSMUSG00000029036 | Atad3a | -1.182 | 8.55E-04 | 2.23E-02 |
| ENSMUSG00000020534 | Shmt1 | -1.182 | 2.57E-03 | 4.54E-02 |
| ENSMUSG00000039841 | Zfp800 | -1.182 | 2.71E-04 | 1.02E-02 |
| ENSMUSG00000002984 | Tomm40 | -1.179 | 1.01E-03 | 2.44E-02 |
| ENSMUSG00000041747 | Utp15 | -1.176 | 3.70E-06 | 4.46E-04 |
| ENSMUSG00000020922 | Lsm12 | -1.176 | 1.13E-06 | 1.83E-04 |
| ENSMUSG00000034274 | Thoc5 | -1.175 | 2.61E-03 | 4.60E-02 |
| ENSMUSG00000024501 | Dpysl3 | -1.174 | 3.60E-04 | 1.24E-02 |
| ENSMUSG00000018697 | Aatf | -1.172 | 5.23E-05 | 3.17E-03 |
| ENSMUSG00000031948 | Kars | -1.171 | 1.35E-03 | 2.96E-02 |
| ENSMUSG00000061477 | Rps7 | -1.170 | 1.21E-03 | 2.74E-02 |
| ENSMUSG00000027531 | Impa1 | -1.169 | 9.05E-04 | 2.30E-02 |
| ENSMUSG00000039033 | Tasp1 | -1.169 | 1.32E-03 | 2.92E-02 |
| ENSMUSG00000017801 | Mlx | -1.167 | 2.44E-05 | 1.80E-03 |
| ENSMUSG00000030327 | Necap1 | -1.167 | 4.06E-04 | 1.33E-02 |
| ENSMUSG00000025613 | Cct8 | -1.166 | 7.96E-07 | 1.38E-04 |
| ENSMUSG00000024436 | Mrps18b | -1.165 | 5.96E-05 | 3.49E-03 |
| ENSMUSG00000017561 | Crlf3 | -1.164 | 2.11E-04 | 8.60E-03 |
| ENSMUSG00000020430 | Pes1 | -1.162 | 1.05E-04 | 5.21E-03 |
| ENSMUSG00000047714 | Ppp1r2 | -1.160 | 2.47E-03 | 4.43E-02 |
| ENSMUSG00000001774 | Chordc1 | -1.160 | 3.01E-08 | 1.10E-05 |
| ENSMUSG00000058126 | Tpm3-rs7 | -1.160 | 1.87E-03 | 3.68E-02 |
| ENSMUSG00000111877 | Gm6477 | -1.158 | 1.65E-04 | 7.27E-03 |
| ENSMUSG00000030505 | Prmt3 | -1.155 | 2.19E-03 | 4.09E-02 |
| ENSMUSG00000031146 | Plp2 | -1.155 | 2.86E-03 | 4.88E-02 |
| ENSMUSG00000050697 | Prkaa1 | -1.151 | 1.69E-04 | 7.43E-03 |
| ENSMUSG00000021692 | Dimt1 | -1.150 | 2.05E-04 | 8.49E-03 |
| ENSMUSG00000037563 | Rps16 | -1.146 | 4.07E-04 | 1.33E-02 |

| | | | | |
|---|---|---|---|---|
| ENSMUSG00000042508 | Dmtf1 | -1.146 | 2.29E-04 | 9.04E-03 |
| ENSMUSG00000022538 | Lsg1 | -1.146 | 2.64E-04 | 9.99E-03 |
| ENSMUSG00000002458 | Rgs19 | -1.144 | 1.70E-03 | 3.45E-02 |
| ENSMUSG00000042660 | Wdr55 | -1.142 | 2.20E-03 | 4.10E-02 |
| ENSMUSG00000006599 | Gtf2h1 | -1.142 | 8.27E-04 | 2.18E-02 |
| ENSMUSG00000004264 | Phb2 | -1.141 | 9.06E-04 | 2.30E-02 |
| ENSMUSG00000036902 | Neto2 | -1.140 | 1.84E-04 | 7.81E-03 |
| ENSMUSG00000022663 | Atg3 | -1.138 | 6.45E-05 | 3.67E-03 |
| ENSMUSG00000025134 | Alyref | -1.138 | 2.76E-05 | 1.98E-03 |
| ENSMUSG00000030007 | Cct7 | -1.138 | 3.53E-04 | 1.22E-02 |
| ENSMUSG00000028982 | Slc25a33 | -1.137 | 1.45E-03 | 3.06E-02 |
| ENSMUSG00000042225 | Ammecr1 | -1.134 | 2.13E-05 | 1.64E-03 |
| ENSMUSG00000034681 | Rnps1 | -1.132 | 4.90E-04 | 1.52E-02 |
| ENSMUSG00000027076 | Timm10 | -1.131 | 1.42E-03 | 3.01E-02 |
| ENSMUSG00000057113 | Npm1 | -1.124 | 7.15E-06 | 7.26E-04 |
| ENSMUSG00000033554 | Dph5 | -1.123 | 3.05E-04 | 1.11E-02 |
| ENSMUSG00000028990 | Lzic | -1.117 | 3.25E-04 | 1.17E-02 |
| ENSMUSG00000035960 | Apex1 | -1.115 | 2.52E-03 | 4.49E-02 |
| ENSMUSG00000014226 | Cacybp | -1.114 | 1.84E-03 | 3.64E-02 |
| ENSMUSG00000063524 | Eno1 | -1.113 | 2.29E-03 | 4.22E-02 |
| ENSMUSG00000033285 | Wdr3 | -1.112 | 6.23E-05 | 3.58E-03 |
| ENSMUSG00000059796 | Eif4a1 | -1.111 | 6.37E-05 | 3.65E-03 |
| ENSMUSG00000038046 | Mrm3 | -1.110 | 3.44E-04 | 1.20E-02 |
| ENSMUSG00000032279 | Idh3a | -1.110 | 1.55E-06 | 2.26E-04 |
| ENSMUSG00000024966 | Stip1 | -1.110 | 5.70E-04 | 1.69E-02 |
| ENSMUSG00000025190 | Got1 | -1.105 | 2.41E-05 | 1.80E-03 |
| ENSMUSG00000079435 | Rpl36a | -1.102 | 2.03E-03 | 3.88E-02 |
| ENSMUSG00000030662 | Ipo5 | -1.097 | 2.16E-06 | 2.94E-04 |
| ENSMUSG00000061079 | Zfp143 | -1.097 | 1.10E-03 | 2.57E-02 |
| ENSMUSG00000003070 | Efna2 | -1.097 | 2.57E-03 | 4.54E-02 |
| ENSMUSG00000003808 | Farsa | -1.095 | 6.19E-04 | 1.79E-02 |
| ENSMUSG00000039660 | Spout1 | -1.095 | 2.24E-03 | 4.16E-02 |
| ENSMUSG00000021958 | Pinx1 | -1.094 | 1.57E-04 | 7.04E-03 |
| ENSMUSG00000032714 | Syde1 | -1.091 | 2.91E-03 | 4.94E-02 |
| ENSMUSG00000027671 | Actl6a | -1.090 | 6.11E-05 | 3.53E-03 |
| ENSMUSG00000007029 | Vars | -1.085 | 9.53E-04 | 2.36E-02 |
| ENSMUSG00000020484 | Xbp1 | -1.082 | 1.42E-03 | 3.01E-02 |
| ENSMUSG00000070319 | Eif3g | -1.081 | 1.41E-06 | 2.14E-04 |
| ENSMUSG00000022752 | Tomm70a | -1.080 | 3.95E-04 | 1.31E-02 |
| ENSMUSG00000042369 | Rbm45 | -1.077 | 1.68E-04 | 7.38E-03 |
| ENSMUSG00000024528 | Srfbp1 | -1.077 | 4.62E-04 | 1.46E-02 |
| ENSMUSG00000031403 | Dkc1 | -1.073 | 8.40E-05 | 4.46E-03 |
| ENSMUSG00000021578 | Ccdc127 | -1.071 | 5.89E-06 | 6.30E-04 |
| ENSMUSG00000007739 | Cct4 | -1.068 | 1.24E-04 | 5.89E-03 |
| ENSMUSG00000015748 | Prpf3 | -1.066 | 2.70E-03 | 4.70E-02 |
| ENSMUSG00000028252 | Ccnc | -1.063 | 1.05E-03 | 2.49E-02 |
| ENSMUSG00000025630 | Hprt | -1.063 | 1.25E-03 | 2.81E-02 |

| | | | | |
|---|---|---|---|---|
| ENSMUSG00000033781 | Asb13 | -1.063 | 5.51E-04 | 1.65E-02 |
| ENSMUSG00000034203 | Chchd4 | -1.061 | 1.53E-05 | 1.28E-03 |
| ENSMUSG00000056962 | Jmjd6 | -1.060 | 1.91E-03 | 3.73E-02 |
| ENSMUSG00000027429 | Sec23b | -1.057 | 1.46E-04 | 6.67E-03 |
| ENSMUSG00000001783 | Rtcb | -1.051 | 3.50E-04 | 1.21E-02 |
| ENSMUSG00000030045 | Mrpl19 | -1.050 | 1.24E-03 | 2.79E-02 |
| ENSMUSG00000030521 | Mphosph10 | -1.050 | 1.39E-03 | 2.99E-02 |
| ENSMUSG00000028224 | Nbn | -1.047 | 1.09E-03 | 2.56E-02 |
| ENSMUSG00000022336 | Eif3e | -1.046 | 2.13E-03 | 4.01E-02 |
| ENSMUSG00000038482 | Tfdp1 | -1.038 | 2.84E-05 | 2.02E-03 |
| ENSMUSG00000006998 | Psmd2 | -1.037 | 1.96E-03 | 3.80E-02 |
| ENSMUSG00000071172 | Srsf3 | -1.035 | 1.69E-03 | 3.45E-02 |
| ENSMUSG00000097195 | Snhg5 | -1.034 | 2.73E-03 | 4.72E-02 |
| ENSMUSG00000030357 | Fkbp4 | -1.034 | 2.40E-03 | 4.36E-02 |
| ENSMUSG00000027122 | Arl14ep | -1.033 | 1.39E-03 | 2.99E-02 |
| ENSMUSG00000030942 | Thumpd1 | -1.028 | 7.75E-05 | 4.17E-03 |
| ENSMUSG00000021621 | Zcchc9 | -1.028 | 1.54E-03 | 3.21E-02 |
| ENSMUSG00000047945 | Marcksl1 | -1.025 | 2.68E-03 | 4.69E-02 |
| ENSMUSG00000023883 | Phf10 | -1.021 | 5.61E-04 | 1.67E-02 |
| ENSMUSG00000028790 | Khdrbs1 | -1.019 | 1.11E-03 | 2.58E-02 |
| ENSMUSG00000063884 | Ptcd3 | -1.016 | 9.93E-05 | 5.05E-03 |
| ENSMUSG00000034424 | Gcsh | -1.015 | 1.25E-03 | 2.80E-02 |
| ENSMUSG00000064181 | Rab3ip | -1.014 | 2.44E-03 | 4.40E-02 |
| ENSMUSG00000001674 | Ddx18 | -1.012 | 1.25E-04 | 5.92E-03 |
| ENSMUSG00000019782 | Rwdd1 | -1.011 | 1.91E-04 | 8.06E-03 |
| ENSMUSG00000030264 | Thumpd3 | -1.009 | 1.79E-04 | 7.70E-03 |
| ENSMUSG00000028869 | Gnl2 | -1.007 | 5.71E-04 | 1.69E-02 |
| ENSMUSG00000035575 | Utp6 | -1.006 | 5.94E-04 | 1.74E-02 |
| ENSMUSG00000022814 | Umps | -1.006 | 3.15E-05 | 2.17E-03 |
| ENSMUSG00000025393 | Atp5b | -1.003 | 3.46E-04 | 1.21E-02 |
| ENSMUSG00000008373 | Prpf31 | -1.000 | 5.59E-04 | 1.67E-02 |
| ENSMUSG00000013701 | Timm23 | -0.999 | 7.54E-05 | 4.11E-03 |
| ENSMUSG00000030881 | Arfip2 | -0.998 | 6.41E-04 | 1.83E-02 |
| ENSMUSG00000032026 | Rexo2 | -0.995 | 1.52E-03 | 3.19E-02 |
| ENSMUSG00000040124 | Gorab | -0.990 | 1.21E-03 | 2.74E-02 |
| ENSMUSG00000041733 | Coq5 | -0.988 | 2.76E-03 | 4.77E-02 |
| ENSMUSG00000022800 | Fyttd1 | -0.988 | 1.80E-03 | 3.60E-02 |
| ENSMUSG00000030224 | Strap | -0.987 | 5.51E-04 | 1.65E-02 |
| ENSMUSG00000026159 | Agfg1 | -0.985 | 5.87E-05 | 3.45E-03 |
| ENSMUSG00000042719 | Naa25 | -0.984 | 2.20E-04 | 8.83E-03 |
| ENSMUSG00000021018 | Polr2h | -0.981 | 7.64E-06 | 7.41E-04 |
| ENSMUSG00000021737 | Psmd6 | -0.980 | 2.95E-07 | 6.72E-05 |
| ENSMUSG00000023147 | Wrb | -0.979 | 6.75E-05 | 3.83E-03 |
| ENSMUSG00000028333 | Anp32b | -0.974 | 1.82E-05 | 1.46E-03 |
| ENSMUSG00000018326 | Ywhab | -0.973 | 3.39E-04 | 1.20E-02 |
| ENSMUSG00000017781 | Pitpna | -0.973 | 1.85E-03 | 3.66E-02 |
| ENSMUSG00000002319 | Ipo4 | -0.969 | 2.70E-03 | 4.70E-02 |

| | | | | |
|---|---|---|---|---|
| ENSMUSG00000001891 | Ugp2 | -0.969 | 3.28E-04 | 1.17E-02 |
| ENSMUSG00000022466 | Rpap3 | -0.965 | 7.03E-04 | 1.94E-02 |
| ENSMUSG00000022035 | Ccdc25 | -0.962 | 1.42E-04 | 6.53E-03 |
| ENSMUSG00000038827 | Fam206a | -0.962 | 4.12E-06 | 4.83E-04 |
| ENSMUSG00000067722 | BC003965 | -0.960 | 1.72E-03 | 3.47E-02 |
| ENSMUSG00000030061 | Uba3 | -0.959 | 1.02E-04 | 5.14E-03 |
| ENSMUSG00000020088 | Sar1a | -0.958 | 6.92E-04 | 1.92E-02 |
| ENSMUSG00000028034 | Fubp1 | -0.956 | 9.58E-04 | 2.36E-02 |
| ENSMUSG00000030603 | Psmc4 | -0.956 | 1.64E-04 | 7.25E-03 |
| ENSMUSG00000032058 | Ppp2r1b | -0.955 | 5.22E-04 | 1.59E-02 |
| ENSMUSG00000049751 | Rpl36al | -0.937 | 1.70E-03 | 3.46E-02 |
| ENSMUSG00000049517 | Rps23 | -0.936 | 3.32E-04 | 1.18E-02 |
| ENSMUSG00000078652 | Psme3 | -0.936 | 9.63E-05 | 4.97E-03 |
| ENSMUSG00000040738 | Ints8 | -0.935 | 2.05E-04 | 8.51E-03 |
| ENSMUSG00000033931 | Rbm34 | -0.935 | 4.23E-04 | 1.37E-02 |
| ENSMUSG00000028719 | Cmpk1 | -0.935 | 2.72E-03 | 4.72E-02 |
| ENSMUSG00000062203 | Gspt1 | -0.933 | 4.10E-04 | 1.34E-02 |
| ENSMUSG00000004788 | Eif2b2 | -0.930 | 2.35E-03 | 4.29E-02 |
| ENSMUSG00000032041 | Tirap | -0.924 | 2.85E-03 | 4.88E-02 |
| ENSMUSG00000033953 | Ppp3r1 | -0.919 | 2.33E-04 | 9.14E-03 |
| ENSMUSG00000026240 | Cops7b | -0.918 | 2.41E-03 | 4.38E-02 |
| ENSMUSG00000027162 | Lin7c | -0.917 | 9.40E-04 | 2.34E-02 |
| ENSMUSG00000038845 | Phb | -0.917 | 1.32E-03 | 2.92E-02 |
| ENSMUSG00000024081 | Cebpz | -0.914 | 6.29E-04 | 1.81E-02 |
| ENSMUSG00000020224 | Llph | -0.911 | 1.04E-03 | 2.48E-02 |
| ENSMUSG00000027357 | Crls1 | -0.910 | 4.84E-05 | 2.97E-03 |
| ENSMUSG00000020720 | Psmd12 | -0.907 | 3.47E-04 | 1.21E-02 |
| ENSMUSG00000030298 | Sec13 | -0.904 | 9.44E-04 | 2.34E-02 |
| ENSMUSG00000032458 | Copb2 | -0.899 | 2.90E-03 | 4.93E-02 |
| ENSMUSG00000004268 | Emg1 | -0.895 | 9.16E-05 | 4.78E-03 |
| ENSMUSG00000025439 | Clns1a | -0.895 | 1.21E-03 | 2.74E-02 |
| ENSMUSG00000043998 | Mgat2 | -0.893 | 9.03E-05 | 4.75E-03 |
| ENSMUSG00000001525 | Tubb5 | -0.892 | 8.45E-04 | 2.22E-02 |
| ENSMUSG00000044763 | Trmt10c | -0.889 | 1.04E-03 | 2.48E-02 |
| ENSMUSG00000028322 | Exosc3 | -0.885 | 1.77E-04 | 7.66E-03 |
| ENSMUSG00000021276 | Cinp | -0.884 | 1.36E-03 | 2.96E-02 |
| ENSMUSG00000049755 | Zfp672 | -0.882 | 2.52E-03 | 4.49E-02 |
| ENSMUSG00000031575 | Ash2l | -0.882 | 7.61E-04 | 2.04E-02 |
| ENSMUSG00000021810 | Ecd | -0.881 | 3.92E-04 | 1.31E-02 |
| ENSMUSG00000018965 | Ywhah | -0.881 | 2.92E-03 | 4.95E-02 |
| ENSMUSG00000021660 | Btf3 | -0.879 | 7.07E-04 | 1.95E-02 |
| ENSMUSG00000014504 | Srp19 | -0.875 | 2.93E-03 | 4.96E-02 |
| ENSMUSG00000024878 | Cbwd1 | -0.871 | 3.73E-05 | 2.51E-03 |
| ENSMUSG00000027170 | Eif3m | -0.870 | 3.49E-04 | 1.21E-02 |
| ENSMUSG00000061613 | U2af1 | -0.868 | 2.02E-03 | 3.87E-02 |
| ENSMUSG00000024712 | Rfk | -0.866 | 6.36E-04 | 1.82E-02 |
| ENSMUSG00000055531 | Cpsf6 | -0.865 | 1.26E-03 | 2.81E-02 |

| | | | | |
|---|---|---|---|---|
| ENSMUSG00000061207 | Stk19 | -0.863 | 1.36E-03 | 2.96E-02 |
| ENSMUSG00000038822 | Hace1 | -0.863 | 2.62E-03 | 4.61E-02 |
| ENSMUSG00000028187 | Rpf1 | -0.862 | 8.24E-04 | 2.18E-02 |
| ENSMUSG00000020349 | Ppp2ca | -0.862 | 9.20E-04 | 2.31E-02 |
| ENSMUSG00000024231 | Cul2 | -0.861 | 1.67E-03 | 3.42E-02 |
| ENSMUSG00000027823 | Gmps | -0.860 | 5.99E-04 | 1.75E-02 |
| ENSMUSG00000036398 | Ppp1r11 | -0.860 | 1.88E-03 | 3.68E-02 |
| ENSMUSG00000028907 | Utp11 | -0.859 | 2.36E-04 | 9.19E-03 |
| ENSMUSG00000038274 | Fau | -0.858 | 1.82E-04 | 7.80E-03 |
| ENSMUSG00000034321 | Exosc1 | -0.857 | 1.13E-03 | 2.61E-02 |
| ENSMUSG00000029326 | Enoph1 | -0.856 | 2.97E-03 | 5.00E-02 |
| ENSMUSG00000021546 | Hnrnpk | -0.848 | 2.12E-03 | 4.01E-02 |
| ENSMUSG00000006057 | Atp5g1 | -0.840 | 9.08E-04 | 2.30E-02 |
| ENSMUSG00000020248 | Nfyb | -0.837 | 3.39E-04 | 1.20E-02 |
| ENSMUSG00000001416 | Cct3 | -0.836 | 7.45E-05 | 4.07E-03 |
| ENSMUSG00000026036 | Nif3l1 | -0.834 | 1.10E-03 | 2.57E-02 |
| ENSMUSG00000022052 | Ppp2r2a | -0.831 | 2.13E-03 | 4.01E-02 |
| ENSMUSG00000030880 | Polr3e | -0.822 | 6.86E-04 | 1.91E-02 |
| ENSMUSG00000027905 | Ddx20 | -0.819 | 1.13E-04 | 5.45E-03 |
| ENSMUSG00000021868 | Ppif | -0.817 | 1.70E-03 | 3.45E-02 |
| ENSMUSG00000022285 | Ywhaz | -0.817 | 1.04E-04 | 5.19E-03 |
| ENSMUSG00000007338 | Mrpl49 | -0.817 | 4.62E-04 | 1.46E-02 |
| ENSMUSG00000036371 | Serbp1 | -0.816 | 1.56E-04 | 7.03E-03 |
| ENSMUSG00000047879 | Usp14 | -0.816 | 9.00E-04 | 2.30E-02 |
| ENSMUSG00000015290 | Ubl4a | -0.814 | 1.51E-04 | 6.84E-03 |
| ENSMUSG00000026869 | Psmd5 | -0.812 | 6.84E-04 | 1.91E-02 |
| ENSMUSG00000035021 | Baz1a | -0.811 | 4.45E-04 | 1.41E-02 |
| ENSMUSG00000028889 | Yrdc | -0.810 | 2.95E-03 | 4.98E-02 |
| ENSMUSG00000030879 | Mrpl17 | -0.808 | 2.69E-04 | 1.01E-02 |
| ENSMUSG00000061315 | Naca | -0.804 | 1.33E-03 | 2.94E-02 |
| ENSMUSG00000027601 | Mtfr1 | -0.803 | 3.03E-04 | 1.10E-02 |
| ENSMUSG00000061950 | Ppp4r1 | -0.799 | 2.73E-03 | 4.72E-02 |
| ENSMUSG00000034024 | Cct2 | -0.798 | 2.55E-07 | 5.98E-05 |
| ENSMUSG00000019795 | Pcmt1 | -0.797 | 2.63E-05 | 1.91E-03 |
| ENSMUSG00000040618 | Pck2 | -0.796 | 1.47E-03 | 3.08E-02 |
| ENSMUSG00000027981 | Rnpc3 | -0.794 | 2.19E-04 | 8.81E-03 |
| ENSMUSG00000027714 | Exosc9 | -0.793 | 1.17E-03 | 2.69E-02 |
| ENSMUSG00000055762 | Eef1d | -0.791 | 2.28E-04 | 9.04E-03 |
| ENSMUSG00000074797 | Itpa | -0.790 | 9.81E-04 | 2.39E-02 |
| ENSMUSG00000020775 | Mrpl38 | -0.789 | 1.73E-04 | 7.54E-03 |
| ENSMUSG00000036323 | Srp72 | -0.788 | 9.39E-04 | 2.34E-02 |
| ENSMUSG00000033813 | Tcea1 | -0.785 | 1.78E-03 | 3.57E-02 |
| ENSMUSG00000028902 | Sf3a3 | -0.780 | 1.40E-04 | 6.47E-03 |
| ENSMUSG00000028988 | Ctnnbip1 | -0.775 | 1.31E-03 | 2.90E-02 |
| ENSMUSG00000014195 | Dnajc7 | -0.773 | 7.44E-04 | 2.02E-02 |
| ENSMUSG00000032563 | Mrpl3 | -0.771 | 4.29E-05 | 2.71E-03 |
| ENSMUSG00000025580 | Eif4a3 | -0.753 | 1.93E-03 | 3.76E-02 |

| | | | | |
|---|---|---|---|---|
| ENSMUSG00000031422 | Morf4l2 | -0.741 | 2.78E-04 | 1.03E-02 |
| ENSMUSG00000031256 | Cstf2 | -0.734 | 9.86E-05 | 5.04E-03 |
| ENSMUSG00000029551 | Psmg3 | -0.732 | 8.46E-04 | 2.22E-02 |
| ENSMUSG00000070697 | Utp3 | -0.732 | 1.44E-03 | 3.04E-02 |
| ENSMUSG00000038446 | Cdc40 | -0.732 | 2.13E-03 | 4.01E-02 |
| ENSMUSG00000020708 | Psmc5 | -0.727 | 1.38E-03 | 2.98E-02 |
| ENSMUSG00000021131 | Erh | -0.723 | 2.88E-03 | 4.89E-02 |
| ENSMUSG00000038762 | Abcf1 | -0.720 | 1.56E-03 | 3.24E-02 |
| ENSMUSG00000074781 | Ube2n | -0.718 | 1.81E-03 | 3.61E-02 |
| ENSMUSG00000018565 | Elp5 | -0.707 | 5.99E-05 | 3.49E-03 |
| ENSMUSG00000016018 | Skiv2l2 | -0.706 | 1.29E-03 | 2.86E-02 |
| ENSMUSG00000025917 | Cops5 | -0.700 | 5.28E-04 | 1.60E-02 |
| ENSMUSG00000020180 | Snrpd3 | -0.696 | 1.14E-03 | 2.64E-02 |
| ENSMUSG00000063480 | Snu13 | -0.694 | 2.19E-04 | 8.81E-03 |
| ENSMUSG00000042229 | Rabif | -0.692 | 2.46E-03 | 4.42E-02 |
| ENSMUSG00000024580 | Grpel2 | -0.691 | 3.15E-04 | 1.14E-02 |
| ENSMUSG00000030754 | Copb1 | -0.678 | 9.24E-04 | 2.31E-02 |
| ENSMUSG00000022774 | Ncbp2 | -0.671 | 1.92E-03 | 3.74E-02 |
| ENSMUSG00000026229 | Psmd1 | -0.653 | 1.82E-03 | 3.62E-02 |
| ENSMUSG00000032002 | Dcun1d5 | -0.634 | 9.07E-04 | 2.30E-02 |
| ENSMUSG00000039640 | Mrpl12 | -0.634 | 6.18E-04 | 1.79E-02 |
| ENSMUSG00000030471 | Zdhhc13 | -0.627 | 2.65E-03 | 4.66E-02 |
| ENSMUSG00000046434 | Hnrnpa1 | -0.619 | 2.57E-03 | 4.54E-02 |
| ENSMUSG00000015120 | Ube2i | -0.618 | 1.55E-03 | 3.22E-02 |
| ENSMUSG00000029038 | Ssu72 | -0.616 | 1.01E-03 | 2.45E-02 |
| ENSMUSG00000005683 | Cs | -0.600 | 2.86E-03 | 4.89E-02 |
| ENSMUSG00000024668 | Sdhaf2 | -0.595 | 2.96E-04 | 1.09E-02 |
| ENSMUSG00000044221 | Grsf1 | -0.579 | 2.48E-03 | 4.44E-02 |
| ENSMUSG00000020952 | Scfd1 | -0.572 | 1.44E-03 | 3.04E-02 |
| ENSMUSG00000022204 | Ngdn | -0.526 | 2.19E-03 | 4.08E-02 |
| ENSMUSG00000004364 | Cul3 | -0.501 | 1.81E-03 | 3.61E-02 |
| ENSMUSG00000027104 | Atf2 | 0.579 | 1.42E-03 | 3.01E-02 |
| ENSMUSG00000009030 | Pdcl | 0.601 | 2.16E-03 | 4.05E-02 |
| ENSMUSG00000024248 | Cox7a2l | 0.699 | 2.21E-04 | 8.83E-03 |
| ENSMUSG00000030750 | Nsmce1 | 0.712 | 6.00E-05 | 3.49E-03 |
| ENSMUSG00000027304 | Rtf1 | 0.722 | 1.16E-03 | 2.67E-02 |
| ENSMUSG00000032398 | Snapc5 | 0.725 | 6.80E-04 | 1.91E-02 |
| ENSMUSG00000078570 | 1110065P20 | 0.741 | 2.36E-03 | 4.30E-02 |
| ENSMUSG00000020738 | Sumo2 | 0.747 | 2.36E-03 | 4.29E-02 |
| ENSMUSG00000019942 | Cdk1 | 0.750 | 1.37E-03 | 2.98E-02 |
| ENSMUSG00000025102 | 3110040N11 | 0.751 | 7.78E-05 | 4.18E-03 |
| ENSMUSG00000066724 | Gm10175 | 0.757 | 2.05E-03 | 3.90E-02 |
| ENSMUSG00000021109 | Hif1a | 0.772 | 1.40E-03 | 3.00E-02 |
| ENSMUSG00000031360 | Ctps2 | 0.785 | 1.60E-05 | 1.33E-03 |
| ENSMUSG00000032599 | Ip6k2 | 0.797 | 2.00E-03 | 3.84E-02 |
| ENSMUSG00000019370 | Calm3 | 0.808 | 1.21E-03 | 2.74E-02 |
| ENSMUSG00000026568 | Mpc2 | 0.813 | 5.91E-04 | 1.74E-02 |

| | | | | |
|---|---|---|---|---|
| ENSMUSG00000050732 | Vamp8 | 0.834 | 1.50E-03 | 3.14E-02 |
| ENSMUSG00000039159 | Ube2h | 0.845 | 1.28E-03 | 2.85E-02 |
| ENSMUSG00000050552 | Lamtor4 | 0.865 | 1.94E-03 | 3.76E-02 |
| ENSMUSG00000028066 | Pmf1 | 0.867 | 2.13E-05 | 1.64E-03 |
| ENSMUSG00000031023 | Akip1 | 0.868 | 6.11E-06 | 6.44E-04 |
| ENSMUSG00000017286 | Glod4 | 0.888 | 4.36E-04 | 1.39E-02 |
| ENSMUSG00000028062 | Lamtor2 | 0.890 | 9.07E-04 | 2.30E-02 |
| ENSMUSG00000031534 | Smim19 | 0.890 | 2.65E-04 | 9.99E-03 |
| ENSMUSG00000036639 | Nudt1 | 0.905 | 3.37E-04 | 1.19E-02 |
| ENSMUSG00000033545 | Znrf1 | 0.907 | 1.28E-03 | 2.86E-02 |
| ENSMUSG00000031641 | Cbr4 | 0.914 | 7.54E-04 | 2.04E-02 |
| ENSMUSG00000034744 | Nagk | 0.929 | 2.18E-03 | 4.07E-02 |
| ENSMUSG00000040767 | Snrnp25 | 0.942 | 2.87E-03 | 4.89E-02 |
| ENSMUSG00000030613 | Ccdc90b | 0.943 | 1.10E-04 | 5.33E-03 |
| ENSMUSG00000022092 | Ppp3cc | 0.955 | 9.31E-04 | 2.32E-02 |
| ENSMUSG00000056394 | Lig1 | 0.972 | 1.46E-03 | 3.07E-02 |
| ENSMUSG00000016940 | Kctd2 | 0.979 | 2.66E-03 | 4.66E-02 |
| ENSMUSG00000052915 | Msl1 | 0.986 | 1.51E-03 | 3.17E-02 |
| ENSMUSG00000078566 | Bnip3 | 0.987 | 1.37E-03 | 2.98E-02 |
| ENSMUSG00000045103 | Dmd | 0.989 | 1.76E-03 | 3.53E-02 |
| ENSMUSG00000030301 | Ccdc91 | 0.993 | 2.70E-03 | 4.70E-02 |
| ENSMUSG00000053929 | Cyhr1 | 1.009 | 1.93E-06 | 2.71E-04 |
| ENSMUSG00000049881 | 2810025M15 | 1.016 | 4.00E-05 | 2.60E-03 |
| ENSMUSG00000060147 | Serpinb6a | 1.017 | 1.76E-03 | 3.54E-02 |
| ENSMUSG00000040771 | Oard1 | 1.031 | 1.33E-04 | 6.20E-03 |
| ENSMUSG00000043411 | Usp48 | 1.033 | 8.72E-04 | 2.25E-02 |
| ENSMUSG00000006021 | Kptn | 1.034 | 1.11E-03 | 2.58E-02 |
| ENSMUSG00000030982 | 9030624J02I | 1.035 | 2.77E-03 | 4.78E-02 |
| ENSMUSG00000042364 | Snx18 | 1.038 | 2.23E-04 | 8.89E-03 |
| ENSMUSG00000022817 | Itgb5 | 1.045 | 2.43E-03 | 4.39E-02 |
| ENSMUSG00000005233 | Spc25 | 1.046 | 5.99E-04 | 1.75E-02 |
| ENSMUSG00000027160 | Ccdc34 | 1.059 | 7.56E-05 | 4.11E-03 |
| ENSMUSG00000040841 | Six5 | 1.072 | 2.84E-04 | 1.06E-02 |
| ENSMUSG00000003778 | Brd8 | 1.075 | 3.88E-04 | 1.30E-02 |
| ENSMUSG00000061887 | Ssbp3 | 1.078 | 2.95E-03 | 4.98E-02 |
| ENSMUSG00000033111 | 3830406C13 | 1.080 | 1.35E-05 | 1.16E-03 |
| ENSMUSG00000029059 | Fam213b | 1.084 | 1.19E-03 | 2.72E-02 |
| ENSMUSG00000035901 | Dennd5a | 1.090 | 3.79E-04 | 1.28E-02 |
| ENSMUSG00000044201 | Cdc25c | 1.102 | 5.02E-04 | 1.55E-02 |
| ENSMUSG00000042293 | Gm5617 | 1.111 | 1.57E-03 | 3.26E-02 |
| ENSMUSG00000049225 | Pdp1 | 1.119 | 7.24E-06 | 7.26E-04 |
| ENSMUSG00000028969 | Cdk5 | 1.121 | 2.14E-03 | 4.03E-02 |
| ENSMUSG00000032038 | St3gal4 | 1.133 | 3.86E-04 | 1.30E-02 |
| ENSMUSG00000035637 | Grhpr | 1.134 | 1.82E-04 | 7.80E-03 |
| ENSMUSG00000030213 | Atf7ip | 1.136 | 2.83E-05 | 2.02E-03 |
| ENSMUSG00000019689 | Fmc1 | 1.137 | 9.68E-04 | 2.37E-02 |
| ENSMUSG00000044252 | Osbpl1a | 1.146 | 2.21E-04 | 8.83E-03 |

| | | | | |
|---|---|---|---|---|
| ENSMUSG00000021033 | Gstz1 | 1.173 | 7.41E-06 | 7.31E-04 |
| ENSMUSG00000027459 | Fam110a | 1.173 | 2.36E-03 | 4.29E-02 |
| ENSMUSG00000040549 | Ckap5 | 1.175 | 2.99E-04 | 1.09E-02 |
| ENSMUSG00000022051 | Bnip3l | 1.176 | 4.01E-06 | 4.73E-04 |
| ENSMUSG00000078695 | Cisd3 | 1.179 | 1.63E-03 | 3.35E-02 |
| ENSMUSG00000052139 | Babam2 | 1.180 | 2.08E-04 | 8.54E-03 |
| ENSMUSG00000060538 | Tmem219 | 1.180 | 1.65E-03 | 3.38E-02 |
| ENSMUSG00000001403 | Ube2c | 1.180 | 6.11E-05 | 3.53E-03 |
| ENSMUSG00000002580 | Mien1 | 1.187 | 1.19E-05 | 1.04E-03 |
| ENSMUSG00000006717 | Acot13 | 1.187 | 8.59E-06 | 8.04E-04 |
| ENSMUSG00000028832 | Stmn1 | 1.188 | 8.61E-04 | 2.24E-02 |
| ENSMUSG00000039795 | Zfand1 | 1.192 | 1.59E-04 | 7.12E-03 |
| ENSMUSG00000073481 | 44987 | 1.195 | 5.12E-06 | 5.75E-04 |
| ENSMUSG00000022314 | Rad21 | 1.195 | 6.22E-04 | 1.80E-02 |
| ENSMUSG00000035992 | Fnip1 | 1.200 | 1.92E-03 | 3.74E-02 |
| ENSMUSG00000072772 | Grcc10 | 1.204 | 1.63E-04 | 7.25E-03 |
| ENSMUSG00000030990 | Pgap2 | 1.206 | 9.15E-05 | 4.78E-03 |
| ENSMUSG00000029177 | Cenpa | 1.209 | 3.43E-06 | 4.22E-04 |
| ENSMUSG00000033713 | Foxn3 | 1.210 | 1.86E-03 | 3.66E-02 |
| ENSMUSG00000028070 | Naxe | 1.220 | 8.46E-07 | 1.43E-04 |
| ENSMUSG00000019842 | Traf3ip2 | 1.226 | 1.83E-03 | 3.63E-02 |
| ENSMUSG00000073633 | Fbxo36 | 1.229 | 1.61E-03 | 3.32E-02 |
| ENSMUSG00000002804 | Nudt14 | 1.230 | 9.18E-04 | 2.31E-02 |
| ENSMUSG00000021215 | Net1 | 1.231 | 2.03E-03 | 3.88E-02 |
| ENSMUSG00000037499 | Nenf | 1.234 | 1.91E-03 | 3.73E-02 |
| ENSMUSG00000024253 | Dync2li1 | 1.237 | 5.94E-05 | 3.48E-03 |
| ENSMUSG00000033685 | Ucp2 | 1.257 | 3.49E-04 | 1.21E-02 |
| ENSMUSG00000057103 | Nat8f1 | 1.258 | 1.43E-03 | 3.04E-02 |
| ENSMUSG00000023094 | Msrb2 | 1.274 | 7.48E-04 | 2.03E-02 |
| ENSMUSG00000025268 | Maged2 | 1.278 | 1.53E-03 | 3.20E-02 |
| ENSMUSG00000034449 | Dhrs11 | 1.280 | 2.43E-03 | 4.40E-02 |
| ENSMUSG00000024392 | Bag6 | 1.283 | 3.84E-04 | 1.30E-02 |
| ENSMUSG00000042712 | Tceal9 | 1.285 | 2.00E-03 | 3.84E-02 |
| ENSMUSG00000022323 | Rida | 1.287 | 1.98E-03 | 3.83E-02 |
| ENSMUSG00000035048 | Anapc13 | 1.296 | 1.09E-03 | 2.56E-02 |
| ENSMUSG00000006715 | Gmnn | 1.296 | 6.03E-04 | 1.75E-02 |
| ENSMUSG00000027165 | B230118H07 | 1.302 | 1.10E-04 | 5.33E-03 |
| ENSMUSG00000023055 | Calcoco1 | 1.317 | 8.90E-04 | 2.28E-02 |
| ENSMUSG00000033429 | Mcee | 1.317 | 4.25E-05 | 2.71E-03 |
| ENSMUSG00000034354 | Mtmr3 | 1.325 | 6.51E-04 | 1.85E-02 |
| ENSMUSG00000023966 | Rsph9 | 1.326 | 2.82E-06 | 3.59E-04 |
| ENSMUSG00000027332 | Ivd | 1.326 | 1.10E-04 | 5.33E-03 |
| ENSMUSG00000048922 | Cdca2 | 1.332 | 4.05E-04 | 1.33E-02 |
| ENSMUSG00000031644 | Nek1 | 1.336 | 2.96E-03 | 4.99E-02 |
| ENSMUSG00000036644 | Tbc1d9b | 1.340 | 2.61E-04 | 9.90E-03 |
| ENSMUSG00000002058 | Unc119 | 1.342 | 9.83E-05 | 5.04E-03 |
| ENSMUSG00000046341 | Gm11223 | 1.345 | 8.22E-04 | 2.18E-02 |

| | | | | |
|---|---|---|---|---|
| ENSMUSG00000097059 | Fam120aos | 1.350 | 2.29E-03 | 4.22E-02 |
| ENSMUSG00000044080 | S100a1 | 1.350 | 9.62E-04 | 2.36E-02 |
| ENSMUSG00000026495 | Efcab2 | 1.359 | 2.10E-03 | 3.98E-02 |
| ENSMUSG00000030727 | Rabep2 | 1.360 | 1.04E-03 | 2.48E-02 |
| ENSMUSG00000045211 | Nudt18 | 1.361 | 1.81E-04 | 7.79E-03 |
| ENSMUSG00000044475 | Ascc1 | 1.362 | 1.34E-04 | 6.23E-03 |
| ENSMUSG00000027454 | Gins1 | 1.372 | 2.81E-03 | 4.82E-02 |
| ENSMUSG00000051817 | Sox12 | 1.372 | 1.90E-04 | 8.01E-03 |
| ENSMUSG00000022661 | Cd200 | 1.373 | 1.98E-03 | 3.82E-02 |
| ENSMUSG00000036782 | Klhl13 | 1.373 | 1.74E-03 | 3.51E-02 |
| ENSMUSG00000004951 | Hspb1 | 1.374 | 1.24E-03 | 2.79E-02 |
| ENSMUSG00000031099 | Smarca1 | 1.376 | 9.29E-04 | 2.32E-02 |
| ENSMUSG00000036764 | Dnajc12 | 1.376 | 7.46E-04 | 2.02E-02 |
| ENSMUSG00000020914 | Top2a | 1.380 | 1.26E-03 | 2.81E-02 |
| ENSMUSG00000022982 | Sod1 | 1.388 | 1.03E-03 | 2.47E-02 |
| ENSMUSG00000025466 | Fuom | 1.394 | 4.76E-04 | 1.49E-02 |
| ENSMUSG00000061461 | Smim20 | 1.394 | 3.86E-04 | 1.30E-02 |
| ENSMUSG00000028470 | Hint2 | 1.397 | 3.87E-05 | 2.56E-03 |
| ENSMUSG00000074749 | Kiz | 1.401 | 1.81E-05 | 1.46E-03 |
| ENSMUSG00000062866 | Phactr2 | 1.402 | 3.63E-04 | 1.24E-02 |
| ENSMUSG00000035142 | Nubpl | 1.402 | 6.93E-05 | 3.86E-03 |
| ENSMUSG00000034032 | Rpap1 | 1.406 | 7.08E-04 | 1.95E-02 |
| ENSMUSG00000000876 | Pxmp4 | 1.431 | 5.50E-04 | 1.65E-02 |
| ENSMUSG00000062031 | Pgghg | 1.432 | 2.34E-03 | 4.28E-02 |
| ENSMUSG00000037628 | Cdkn3 | 1.434 | 3.97E-04 | 1.31E-02 |
| ENSMUSG00000040128 | Pnrc1 | 1.434 | 1.50E-06 | 2.20E-04 |
| ENSMUSG00000028795 | Ccdc28b | 1.436 | 1.82E-08 | 7.95E-06 |
| ENSMUSG00000038286 | Bphl | 1.441 | 4.13E-05 | 2.68E-03 |
| ENSMUSG00000001666 | Ddt | 1.446 | 8.78E-04 | 2.26E-02 |
| ENSMUSG00000098090 | 2700099C18 | 1.447 | 1.67E-03 | 3.41E-02 |
| ENSMUSG00000029022 | Miip | 1.451 | 4.45E-04 | 1.41E-02 |
| ENSMUSG00000028549 | Itgb3bp | 1.453 | 8.86E-05 | 4.68E-03 |
| ENSMUSG00000041308 | Sntb2 | 1.453 | 3.09E-04 | 1.12E-02 |
| ENSMUSG00000045312 | Lhfpl2 | 1.458 | 1.18E-03 | 2.70E-02 |
| ENSMUSG00000032175 | Tyk2 | 1.470 | 2.11E-04 | 8.60E-03 |
| ENSMUSG00000038718 | Pbx3 | 1.483 | 3.25E-05 | 2.23E-03 |
| ENSMUSG00000033577 | Myo6 | 1.485 | 9.98E-04 | 2.42E-02 |
| ENSMUSG00000026211 | Obsl1 | 1.491 | 3.94E-05 | 2.59E-03 |
| ENSMUSG00000051343 | Rab11fip5 | 1.495 | 2.93E-03 | 4.96E-02 |
| ENSMUSG00000097415 | AU020206 | 1.497 | 5.04E-04 | 1.55E-02 |
| ENSMUSG00000022550 | Adck5 | 1.500 | 2.26E-04 | 9.00E-03 |
| ENSMUSG00000066456 | Hmgn3 | 1.515 | 4.05E-05 | 2.63E-03 |
| ENSMUSG00000032295 | Man2c1 | 1.517 | 1.10E-03 | 2.57E-02 |
| ENSMUSG00000002043 | Trappc6a | 1.521 | 3.80E-05 | 2.53E-03 |
| ENSMUSG00000043644 | 0610009L18 | 1.530 | 2.27E-03 | 4.20E-02 |
| ENSMUSG00000045257 | Morn2 | 1.531 | 3.42E-04 | 1.20E-02 |
| ENSMUSG00000033222 | Ttf2 | 1.535 | 1.04E-04 | 5.19E-03 |

| | | | | |
|---|---|---|---|---|
| ENSMUSG00000034522 | Zfp395 | 1.556 | 1.96E-05 | 1.52E-03 |
| ENSMUSG00000086587 | Gm11837 | 1.559 | 1.85E-03 | 3.66E-02 |
| ENSMUSG00000021775 | Nr1d2 | 1.559 | 6.30E-06 | 6.61E-04 |
| ENSMUSG00000028378 | Ptgr1 | 1.560 | 2.66E-03 | 4.66E-02 |
| ENSMUSG00000025404 | R3hdm2 | 1.569 | 1.25E-04 | 5.92E-03 |
| ENSMUSG00000029763 | Exoc4 | 1.572 | 3.48E-04 | 1.21E-02 |
| ENSMUSG00000026664 | Phyh | 1.580 | 3.61E-06 | 4.40E-04 |
| ENSMUSG00000035765 | Dym | 1.587 | 2.88E-05 | 2.05E-03 |
| ENSMUSG00000024866 | Acy3 | 1.589 | 2.34E-04 | 9.16E-03 |
| ENSMUSG00000033488 | Cryzl2 | 1.592 | 2.31E-04 | 9.08E-03 |
| ENSMUSG00000066705 | Fxyd6 | 1.596 | 2.02E-03 | 3.86E-02 |
| ENSMUSG00000032009 | Sesn3 | 1.598 | 2.73E-04 | 1.02E-02 |
| ENSMUSG00000022272 | Myo10 | 1.602 | 9.50E-04 | 2.35E-02 |
| ENSMUSG00000007837 | Prrg2 | 1.605 | 2.33E-03 | 4.26E-02 |
| ENSMUSG00000032121 | Tmem218 | 1.606 | 2.08E-04 | 8.53E-03 |
| ENSMUSG00000039745 | Htatip2 | 1.609 | 1.74E-05 | 1.43E-03 |
| ENSMUSG00000029385 | Ccng2 | 1.609 | 7.32E-07 | 1.33E-04 |
| ENSMUSG00000025993 | Slc40a1 | 1.617 | 4.72E-07 | 9.53E-05 |
| ENSMUSG00000002997 | Prkar2b | 1.631 | 1.10E-03 | 2.57E-02 |
| ENSMUSG00000026939 | Tmem141 | 1.638 | 5.66E-05 | 3.37E-03 |
| ENSMUSG00000066357 | Wdr6 | 1.643 | 2.00E-06 | 2.80E-04 |
| ENSMUSG00000039671 | Zmynd8 | 1.652 | 1.09E-03 | 2.56E-02 |
| ENSMUSG00000039686 | Zer1 | 1.664 | 2.13E-03 | 4.01E-02 |
| ENSMUSG00000032067 | Pts | 1.671 | 4.64E-08 | 1.45E-05 |
| ENSMUSG00000022587 | Ly6e | 1.673 | 3.97E-04 | 1.31E-02 |
| ENSMUSG00000028789 | Azin2 | 1.684 | 2.58E-03 | 4.55E-02 |
| ENSMUSG00000041126 | H2afv | 1.687 | 1.22E-08 | 5.90E-06 |
| ENSMUSG00000044986 | Tst | 1.688 | 1.07E-03 | 2.52E-02 |
| ENSMUSG00000024646 | Cyb5a | 1.696 | 9.20E-04 | 2.31E-02 |
| ENSMUSG00000020263 | Appl2 | 1.701 | 4.92E-04 | 1.52E-02 |
| ENSMUSG00000103962 | Gm38213 | 1.702 | 2.17E-03 | 4.06E-02 |
| ENSMUSG00000046329 | Slc25a23 | 1.707 | 8.70E-04 | 2.25E-02 |
| ENSMUSG00000058756 | Thra | 1.707 | 3.27E-04 | 1.17E-02 |
| ENSMUSG00000028497 | Hacd4 | 1.720 | 9.89E-04 | 2.41E-02 |
| ENSMUSG00000042675 | Ypel3 | 1.725 | 3.92E-09 | 2.25E-06 |
| ENSMUSG00000024975 | Pdcd4 | 1.726 | 1.02E-05 | 9.15E-04 |
| ENSMUSG00000073755 | 5730409E04 | 1.726 | 1.31E-03 | 2.90E-02 |
| ENSMUSG00000034908 | Sidt2 | 1.727 | 2.06E-03 | 3.92E-02 |
| ENSMUSG00000002055 | Spag5 | 1.740 | 1.20E-03 | 2.74E-02 |
| ENSMUSG00000021619 | Atg10 | 1.740 | 8.46E-04 | 2.22E-02 |
| ENSMUSG00000041219 | Arhgap11a | 1.742 | 1.38E-03 | 2.98E-02 |
| ENSMUSG00000048581 | E130311K13 | 1.744 | 2.54E-05 | 1.86E-03 |
| ENSMUSG00000067336 | Bmpr2 | 1.753 | 2.28E-03 | 4.22E-02 |
| ENSMUSG00000020898 | Ctc1 | 1.755 | 1.75E-05 | 1.43E-03 |
| ENSMUSG00000022390 | Zc3h7b | 1.761 | 5.92E-04 | 1.74E-02 |
| ENSMUSG00000038070 | Cntln | 1.770 | 2.05E-04 | 8.49E-03 |
| ENSMUSG00000013236 | Ptprs | 1.792 | 1.50E-03 | 3.15E-02 |

| | | | | |
|---|---|---|---|---|
| ENSMUSG00000023505 | Cdca3 | 1.798 | 1.29E-08 | 5.97E-06 |
| ENSMUSG00000069835 | Sat2 | 1.806 | 1.56E-03 | 3.23E-02 |
| ENSMUSG00000022217 | Emc9 | 1.807 | 5.08E-05 | 3.10E-03 |
| ENSMUSG00000028597 | Gpx7 | 1.810 | 2.45E-03 | 4.41E-02 |
| ENSMUSG00000035642 | Aamdc | 1.816 | 4.10E-07 | 8.95E-05 |
| ENSMUSG00000010362 | Rdm1 | 1.821 | 4.59E-07 | 9.38E-05 |
| ENSMUSG00000079598 | Clec2l | 1.823 | 1.43E-03 | 3.04E-02 |
| ENSMUSG00000058135 | Gstm1 | 1.831 | 1.36E-05 | 1.16E-03 |
| ENSMUSG00000083307 | AA414768 | 1.845 | 7.32E-04 | 2.00E-02 |
| ENSMUSG00000032218 | Ccnb2 | 1.848 | 2.41E-06 | 3.19E-04 |
| ENSMUSG00000085457 | 1110046J04I | 1.850 | 1.08E-04 | 5.30E-03 |
| ENSMUSG00000038248 | Sobp | 1.869 | 4.24E-04 | 1.37E-02 |
| ENSMUSG00000036523 | Greb1 | 1.870 | 2.13E-03 | 4.01E-02 |
| ENSMUSG00000107369 | Gstm2-ps1 | 1.887 | 8.09E-06 | 7.69E-04 |
| ENSMUSG00000029521 | Chek2 | 1.889 | 2.22E-05 | 1.69E-03 |
| ENSMUSG00000060985 | Tdrd5 | 1.898 | 4.01E-04 | 1.32E-02 |
| ENSMUSG00000039450 | Dcxr | 1.923 | 2.57E-08 | 9.66E-06 |
| ENSMUSG00000031453 | Rasa3 | 1.929 | 6.38E-06 | 6.67E-04 |
| ENSMUSG00000003849 | Nqo1 | 1.932 | 2.02E-03 | 3.86E-02 |
| ENSMUSG00000040557 | Mettl27 | 1.949 | 4.37E-08 | 1.42E-05 |
| ENSMUSG00000000538 | Tom1l2 | 1.955 | 1.42E-03 | 3.01E-02 |
| ENSMUSG00000112876 | Gm32443 | 1.958 | 1.08E-03 | 2.54E-02 |
| ENSMUSG00000027227 | Sord | 1.961 | 2.00E-03 | 3.84E-02 |
| ENSMUSG00000040296 | Ddx58 | 1.971 | 1.02E-03 | 2.45E-02 |
| ENSMUSG00000028199 | Cryz | 1.978 | 7.41E-05 | 4.07E-03 |
| ENSMUSG00000027654 | Fam83d | 1.985 | 1.15E-06 | 1.84E-04 |
| ENSMUSG00000035168 | Tanc1 | 1.988 | 6.27E-04 | 1.81E-02 |
| ENSMUSG00000094910 | D430019H16 | 1.990 | 5.24E-04 | 1.60E-02 |
| ENSMUSG00000068758 | Il3ra | 1.994 | 1.53E-03 | 3.20E-02 |
| ENSMUSG00000033715 | Akr1c14 | 1.994 | 1.57E-04 | 7.04E-03 |
| ENSMUSG00000038252 | Ncapd2 | 2.002 | 1.09E-04 | 5.31E-03 |
| ENSMUSG00000030319 | Cand2 | 2.005 | 6.83E-04 | 1.91E-02 |
| ENSMUSG00000025545 | Clybl | 2.034 | 7.90E-06 | 7.61E-04 |
| ENSMUSG00000020388 | Pdlim4 | 2.037 | 1.83E-05 | 1.47E-03 |
| ENSMUSG00000040249 | Lrp1 | 2.037 | 1.60E-04 | 7.15E-03 |
| ENSMUSG00000021065 | Fut8 | 2.038 | 1.77E-05 | 1.44E-03 |
| ENSMUSG00000021884 | Hacl1 | 2.038 | 1.16E-05 | 1.02E-03 |
| ENSMUSG00000087336 | Gm15860 | 2.060 | 3.19E-04 | 1.15E-02 |
| ENSMUSG00000085614 | 1700123M08 | 2.069 | 6.04E-04 | 1.76E-02 |
| ENSMUSG00000024940 | Ltbp3 | 2.075 | 1.46E-03 | 3.07E-02 |
| ENSMUSG00000086382 | Chrna1os | 2.087 | 1.95E-04 | 8.14E-03 |
| ENSMUSG00000046718 | Bst2 | 2.092 | 1.23E-04 | 5.87E-03 |
| ENSMUSG00000049086 | Bmyc | 2.103 | 2.10E-04 | 8.59E-03 |
| ENSMUSG00000020486 | 45173 | 2.106 | 6.68E-04 | 1.88E-02 |
| ENSMUSG00000027983 | Cyp2u1 | 2.110 | 6.92E-04 | 1.92E-02 |
| ENSMUSG00000059645 | Gm7361 | 2.113 | 7.55E-04 | 2.04E-02 |
| ENSMUSG00000041000 | Trim62 | 2.123 | 2.27E-04 | 9.03E-03 |

| | | | | |
|---|---|---|---|---|
| ENSMUSG00000010307 | Tmem86a | 2.125 | 7.01E-05 | 3.89E-03 |
| ENSMUSG00000022773 | Ypel1 | 2.137 | 4.19E-07 | 9.04E-05 |
| ENSMUSG00000026104 | Stat1 | 2.155 | 9.83E-04 | 2.39E-02 |
| ENSMUSG00000091561 | Gm6665 | 2.158 | 5.20E-06 | 5.82E-04 |
| ENSMUSG00000032224 | Fam81a | 2.159 | 1.40E-03 | 2.99E-02 |
| ENSMUSG00000004035 | Gstm7 | 2.164 | 1.44E-03 | 3.05E-02 |
| ENSMUSG00000057143 | Trim12c | 2.188 | 2.06E-03 | 3.92E-02 |
| ENSMUSG00000040562 | Gstm2 | 2.201 | 5.70E-06 | 6.19E-04 |
| ENSMUSG00000044167 | Foxo1 | 2.223 | 1.76E-04 | 7.64E-03 |
| ENSMUSG00000021094 | Dhrs7 | 2.228 | 1.53E-03 | 3.19E-02 |
| ENSMUSG00000010651 | Acaa1b | 2.244 | 2.19E-03 | 4.09E-02 |
| ENSMUSG00000031666 | Rbl2 | 2.254 | 1.27E-06 | 1.99E-04 |
| ENSMUSG00000052676 | Zmat1 | 2.265 | 2.42E-03 | 4.39E-02 |
| ENSMUSG00000110935 | Gm8834 | 2.278 | 1.99E-04 | 8.32E-03 |
| ENSMUSG00000027890 | Gstm4 | 2.289 | 6.74E-06 | 6.91E-04 |
| ENSMUSG00000060675 | Pla2g16 | 2.292 | 2.71E-06 | 3.48E-04 |
| ENSMUSG00000014164 | Klhl3 | 2.304 | 2.19E-04 | 8.81E-03 |
| ENSMUSG00000001313 | Rnd2 | 2.316 | 2.15E-03 | 4.03E-02 |
| ENSMUSG00000031310 | Zmym3 | 2.329 | 5.61E-05 | 3.34E-03 |
| ENSMUSG00000033610 | Pank1 | 2.333 | 1.31E-03 | 2.90E-02 |
| ENSMUSG00000032221 | Mns1 | 2.338 | 4.30E-05 | 2.71E-03 |
| ENSMUSG00000111080 | Gm20300 | 2.349 | 1.15E-05 | 1.02E-03 |
| ENSMUSG00000098318 | Lockd | 2.357 | 1.86E-05 | 1.47E-03 |
| ENSMUSG00000068551 | Zfp467 | 2.385 | 2.21E-03 | 4.10E-02 |
| ENSMUSG00000057176 | Ccdc189 | 2.385 | 1.04E-03 | 2.48E-02 |
| ENSMUSG00000054612 | Mgmt | 2.414 | 4.48E-10 | 3.37E-07 |
| ENSMUSG00000064215 | Ifi27 | 2.416 | 5.36E-13 | 1.17E-09 |
| ENSMUSG00000041476 | Smpx | 2.418 | 1.26E-05 | 1.10E-03 |
| ENSMUSG00000033420 | Antxr1 | 2.434 | 9.57E-04 | 2.36E-02 |
| ENSMUSG00000100155 | Gm2109 | 2.444 | 2.93E-03 | 4.96E-02 |
| ENSMUSG00000078350 | Smim1 | 2.483 | 6.45E-04 | 1.84E-02 |
| ENSMUSG00000079491 | H2-T10 | 2.490 | 5.10E-04 | 1.57E-02 |
| ENSMUSG00000051048 | P4ha3 | 2.499 | 5.37E-05 | 3.24E-03 |
| ENSMUSG00000105471 | A430073D23 | 2.517 | 2.65E-04 | 9.99E-03 |
| ENSMUSG00000073758 | Sh3d21 | 2.521 | 1.37E-04 | 6.34E-03 |
| ENSMUSG00000026875 | Traf1 | 2.523 | 1.35E-03 | 2.96E-02 |
| ENSMUSG00000020216 | Jsrp1 | 2.526 | 7.54E-06 | 7.36E-04 |
| ENSMUSG00000027253 | Lrp4 | 2.554 | 1.72E-03 | 3.47E-02 |
| ENSMUSG00000087380 | 2210408F21 | 2.556 | 1.04E-04 | 5.19E-03 |
| ENSMUSG00000027966 | Col11a1 | 2.592 | 9.85E-05 | 5.04E-03 |
| ENSMUSG00000027330 | Cdc25b | 2.613 | 8.91E-06 | 8.27E-04 |
| ENSMUSG00000028015 | Ctso | 2.619 | 7.38E-04 | 2.01E-02 |
| ENSMUSG00000001520 | Nrip2 | 2.634 | 1.31E-04 | 6.13E-03 |
| ENSMUSG00000062901 | Klhl24 | 2.649 | 1.59E-10 | 1.65E-07 |
| ENSMUSG00000055692 | Tmem191c | 2.662 | 8.11E-04 | 2.15E-02 |
| ENSMUSG00000085829 | Gm4285 | 2.662 | 8.06E-04 | 2.14E-02 |
| ENSMUSG00000002409 | Dyrk1b | 2.668 | 2.71E-04 | 1.02E-02 |

| | | | | |
|---|---|---|---|---|
| ENSMUSG00000034664 | Itga2b | 2.697 | 2.45E-03 | 4.41E-02 |
| ENSMUSG00000037440 | Vnn1 | 2.728 | 1.01E-03 | 2.44E-02 |
| ENSMUSG00000103839 | Gm37607 | 2.738 | 8.42E-04 | 2.21E-02 |
| ENSMUSG00000036545 | Adamts2 | 2.764 | 6.83E-05 | 3.83E-03 |
| ENSMUSG00000022389 | Tef | 2.822 | 7.99E-07 | 1.38E-04 |
| ENSMUSG00000085315 | A430018G15 | 2.824 | 5.52E-04 | 1.65E-02 |
| ENSMUSG00000035407 | Kank4 | 2.834 | 1.32E-03 | 2.92E-02 |
| ENSMUSG00000021416 | Eci3 | 2.843 | 1.01E-03 | 2.44E-02 |
| ENSMUSG00000040483 | Xaf1 | 2.864 | 2.88E-03 | 4.90E-02 |
| ENSMUSG00000041132 | N4bp2l1 | 2.884 | 7.00E-04 | 1.94E-02 |
| ENSMUSG00000019301 | Hsd17b1 | 2.910 | 1.99E-03 | 3.83E-02 |
| ENSMUSG00000097403 | 9230116N13 | 2.920 | 1.02E-03 | 2.45E-02 |
| ENSMUSG00000113031 | Gm34552 | 2.924 | 2.09E-03 | 3.97E-02 |
| ENSMUSG00000001663 | Gstt1 | 2.927 | 4.49E-06 | 5.21E-04 |
| ENSMUSG00000030793 | Pycard | 2.953 | 4.65E-06 | 5.32E-04 |
| ENSMUSG00000067698 | Gm10220 | 2.994 | 4.54E-05 | 2.83E-03 |
| ENSMUSG00000039533 | Mmd2 | 3.006 | 6.58E-04 | 1.86E-02 |
| ENSMUSG00000102979 | Gm37350 | 3.010 | 2.70E-03 | 4.70E-02 |
| ENSMUSG00000034706 | Dnaic2 | 3.018 | 1.97E-03 | 3.80E-02 |
| ENSMUSG00000020962 | Gtf2a1 | 3.037 | 1.81E-06 | 2.60E-04 |
| ENSMUSG00000112646 | Gm5654 | 3.054 | 2.07E-04 | 8.53E-03 |
| ENSMUSG00000040666 | Sh3bgr | 3.069 | 8.25E-04 | 2.18E-02 |
| ENSMUSG00000085776 | 5430402O13 | 3.088 | 1.05E-03 | 2.49E-02 |
| ENSMUSG00000002265 | Peg3 | 3.090 | 2.84E-03 | 4.87E-02 |
| ENSMUSG00000030380 | Mzf1 | 3.100 | 8.54E-04 | 2.23E-02 |
| ENSMUSG00000031808 | Slc27a1 | 3.100 | 1.58E-03 | 3.26E-02 |
| ENSMUSG00000067700 | Gm5862 | 3.112 | 1.16E-03 | 2.66E-02 |
| ENSMUSG00000044122 | Proca1 | 3.137 | 9.20E-04 | 2.31E-02 |
| ENSMUSG00000116097 | AL590144.2 | 3.143 | 9.45E-06 | 8.59E-04 |
| ENSMUSG00000028211 | Trp53inp1 | 3.146 | 8.11E-06 | 7.69E-04 |
| ENSMUSG00000036086 | Zranb3 | 3.157 | 9.20E-04 | 2.31E-02 |
| ENSMUSG00000035509 | Fbxl21 | 3.166 | 2.51E-03 | 4.48E-02 |
| ENSMUSG00000022235 | Cmbl | 3.172 | 1.16E-07 | 3.25E-05 |
| ENSMUSG00000027217 | Tspan18 | 3.179 | 3.74E-04 | 1.27E-02 |
| ENSMUSG00000089686 | Mnd1-ps | 3.190 | 2.02E-03 | 3.87E-02 |
| ENSMUSG00000003273 | Car11 | 3.204 | 3.03E-04 | 1.10E-02 |
| ENSMUSG00000001665 | Gstt3 | 3.208 | 7.92E-06 | 7.61E-04 |
| ENSMUSG00000024308 | Tapbp | 3.223 | 3.56E-04 | 1.22E-02 |
| ENSMUSG00000028264 | Spaca1 | 3.226 | 1.37E-03 | 2.98E-02 |
| ENSMUSG00000039543 | Cfap70 | 3.231 | 1.78E-03 | 3.57E-02 |
| ENSMUSG00000038776 | Ephx1 | 3.240 | 1.91E-03 | 3.73E-02 |
| ENSMUSG00000107689 | Gm44386 | 3.254 | 1.10E-06 | 1.80E-04 |
| ENSMUSG00000097354 | 2310001H17 | 3.273 | 7.83E-04 | 2.09E-02 |
| ENSMUSG00000066026 | Dhrs3 | 3.302 | 6.44E-04 | 1.84E-02 |
| ENSMUSG00000026817 | Ak1 | 3.325 | 9.24E-06 | 8.47E-04 |
| ENSMUSG00000022620 | Arsa | 3.341 | 2.60E-03 | 4.59E-02 |
| ENSMUSG00000054580 | Pla2r1 | 3.362 | 1.80E-03 | 3.60E-02 |

| | | | | |
|---|---|---|---|---|
| ENSMUSG00000030257 | Srgap3 | 3.381 | 2.87E-04 | 1.06E-02 |
| ENSMUSG00000034949 | Zfr2 | 3.444 | 2.09E-03 | 3.97E-02 |
| ENSMUSG00000027489 | Necab3 | 3.445 | 1.04E-03 | 2.48E-02 |
| ENSMUSG00000041536 | Serpina3a | 3.466 | 2.76E-04 | 1.03E-02 |
| ENSMUSG00000021708 | Rasgrf2 | 3.479 | 2.93E-03 | 4.96E-02 |
| ENSMUSG00000022438 | Parvb | 3.499 | 2.11E-04 | 8.60E-03 |
| ENSMUSG00000096727 | Psmb9 | 3.533 | 7.65E-07 | 1.35E-04 |
| ENSMUSG00000099354 | 1700124L16 | 3.561 | 9.55E-04 | 2.36E-02 |
| ENSMUSG00000086003 | B230206L02 | 3.602 | 1.41E-03 | 3.01E-02 |
| ENSMUSG00000074771 | Ankef1 | 3.614 | 1.92E-03 | 3.74E-02 |
| ENSMUSG00000073821 | 8030451A03 | 3.673 | 9.90E-04 | 2.41E-02 |
| ENSMUSG00000038453 | Srcin1 | 3.706 | 6.85E-04 | 1.91E-02 |
| ENSMUSG00000037709 | Fam13a | 3.721 | 6.34E-04 | 1.82E-02 |
| ENSMUSG00000086900 | Kcnab3os | 3.725 | 1.04E-03 | 2.48E-02 |
| ENSMUSG00000024818 | Slc25a45 | 3.732 | 1.71E-04 | 7.46E-03 |
| ENSMUSG00000027871 | Hsd3b1 | 3.734 | 1.26E-08 | 5.97E-06 |
| ENSMUSG00000026807 | Ak8 | 3.756 | 7.05E-04 | 1.95E-02 |
| ENSMUSG00000085354 | Gm2044 | 3.766 | 1.14E-03 | 2.64E-02 |
| ENSMUSG00000030708 | Dnajb13 | 3.770 | 2.69E-03 | 4.70E-02 |
| ENSMUSG00000028415 | Spink4 | 3.776 | 1.41E-03 | 3.01E-02 |
| ENSMUSG00000027870 | Hao2 | 3.790 | 6.12E-04 | 1.78E-02 |
| ENSMUSG00000110489 | Gm31659 | 3.808 | 2.32E-03 | 4.25E-02 |
| ENSMUSG00000085055 | Gm15958 | 3.817 | 5.45E-04 | 1.64E-02 |
| ENSMUSG00000086946 | Gm15527 | 3.838 | 2.69E-03 | 4.70E-02 |
| ENSMUSG00000026489 | Coq8a | 3.858 | 1.63E-13 | 5.91E-10 |
| ENSMUSG00000020096 | Tbata | 3.866 | 5.28E-04 | 1.60E-02 |
| ENSMUSG00000036585 | Fgf1 | 3.869 | 1.53E-03 | 3.19E-02 |
| ENSMUSG00000026831 | 1700007K13 | 3.885 | 7.79E-04 | 2.09E-02 |
| ENSMUSG00000019933 | Mrln | 3.909 | 1.95E-03 | 3.77E-02 |
| ENSMUSG00000048458 | Fam212b | 3.919 | 8.83E-04 | 2.27E-02 |
| ENSMUSG00000074218 | Cox7a1 | 3.921 | 2.03E-08 | 8.49E-06 |
| ENSMUSG00000053388 | Trim50 | 3.938 | 1.83E-03 | 3.64E-02 |
| ENSMUSG00000074968 | Ano3 | 3.941 | 2.46E-03 | 4.42E-02 |
| ENSMUSG00000086735 | Gm13977 | 3.946 | 2.30E-03 | 4.22E-02 |
| ENSMUSG00000025468 | Caly | 3.949 | 8.82E-05 | 4.67E-03 |
| ENSMUSG00000025934 | Gsta3 | 3.961 | 4.81E-13 | 1.17E-09 |
| ENSMUSG00000028931 | Kcnab2 | 3.987 | 1.22E-03 | 2.75E-02 |
| ENSMUSG00000112736 | Gm48718 | 3.991 | 7.43E-04 | 2.02E-02 |
| ENSMUSG00000028871 | Rspo1 | 3.996 | 5.84E-05 | 3.45E-03 |
| ENSMUSG00000038415 | Foxq1 | 4.012 | 1.19E-03 | 2.72E-02 |
| ENSMUSG00000068245 | Phf11d | 4.023 | 2.81E-03 | 4.82E-02 |
| ENSMUSG00000057948 | Unc13d | 4.025 | 3.96E-06 | 4.70E-04 |
| ENSMUSG00000022236 | Ropn1l | 4.043 | 3.46E-04 | 1.21E-02 |
| ENSMUSG00000114617 | Gm48766 | 4.046 | 2.69E-03 | 4.69E-02 |
| ENSMUSG00000072941 | Sod3 | 4.048 | 2.52E-04 | 9.62E-03 |
| ENSMUSG00000039217 | Il18 | 4.056 | 1.41E-03 | 3.01E-02 |
| ENSMUSG00000026785 | Pkn3 | 4.056 | 1.39E-03 | 2.99E-02 |

| | | | | |
|---|---|---|---|---|
| ENSMUSG00000074678 | Defb25 | 4.062 | 1.93E-03 | 3.75E-02 |
| ENSMUSG00000017453 | Pipox | 4.062 | 5.55E-04 | 1.66E-02 |
| ENSMUSG00000023935 | Spats1 | 4.072 | 3.13E-05 | 2.17E-03 |
| ENSMUSG00000086813 | Gm13657 | 4.092 | 2.64E-04 | 9.99E-03 |
| ENSMUSG00000032343 | Impg1 | 4.101 | 7.51E-04 | 2.03E-02 |
| ENSMUSG00000039110 | Mycbpap | 4.112 | 8.64E-04 | 2.24E-02 |
| ENSMUSG00000099583 | Hist1h3d | 4.143 | 8.02E-04 | 2.14E-02 |
| ENSMUSG00000108592 | Gm38973 | 4.174 | 2.53E-03 | 4.50E-02 |
| ENSMUSG00000058656 | Samd12 | 4.235 | 1.74E-03 | 3.51E-02 |
| ENSMUSG00000033849 | B3galt2 | 4.250 | 2.42E-03 | 4.38E-02 |
| ENSMUSG00000001985 | Grik3 | 4.259 | 4.15E-04 | 1.35E-02 |
| ENSMUSG00000027347 | Rasgrp1 | 4.259 | 1.11E-03 | 2.58E-02 |
| ENSMUSG00000086405 | 9330198N18 | 4.276 | 2.76E-03 | 4.76E-02 |
| ENSMUSG00000075410 | Prcd | 4.289 | 6.18E-07 | 1.19E-04 |
| ENSMUSG00000108415 | Gm30146 | 4.294 | 1.60E-03 | 3.29E-02 |
| ENSMUSG00000038264 | Sema7a | 4.317 | 1.65E-05 | 1.36E-03 |
| ENSMUSG00000033082 | Clec1a | 4.321 | 9.45E-04 | 2.34E-02 |
| ENSMUSG00000012819 | Cdh23 | 4.323 | 4.51E-05 | 2.82E-03 |
| ENSMUSG00000079017 | Ifi27l2a | 4.328 | 3.25E-06 | 4.03E-04 |
| ENSMUSG00000074595 | Wfdc6a | 4.357 | 5.80E-06 | 6.24E-04 |
| ENSMUSG00000040812 | Agbl2 | 4.359 | 7.05E-04 | 1.95E-02 |
| ENSMUSG00000058952 | Cfi | 4.360 | 9.65E-05 | 4.97E-03 |
| ENSMUSG00000038763 | Alpk3 | 4.363 | 1.37E-03 | 2.98E-02 |
| ENSMUSG00000022763 | Aifm3 | 4.393 | 2.43E-03 | 4.40E-02 |
| ENSMUSG00000043664 | Tmem221 | 4.408 | 1.74E-03 | 3.51E-02 |
| ENSMUSG00000032908 | Sgpp2 | 4.417 | 9.55E-04 | 2.36E-02 |
| ENSMUSG00000038246 | Fam50b | 4.425 | 1.89E-03 | 3.71E-02 |
| ENSMUSG00000026241 | Nppc | 4.436 | 2.56E-03 | 4.54E-02 |
| ENSMUSG00000036395 | Glb1l2 | 4.503 | 9.29E-04 | 2.32E-02 |
| ENSMUSG00000073413 | Ly6g6d | 4.508 | 3.27E-04 | 1.17E-02 |
| ENSMUSG00000101378 | Gm28086 | 4.513 | 6.79E-05 | 3.83E-03 |
| ENSMUSG00000055809 | Dnaaf3 | 4.527 | 1.41E-03 | 3.01E-02 |
| ENSMUSG00000030935 | Acsm3 | 4.547 | 5.81E-04 | 1.71E-02 |
| ENSMUSG00000021185 | Dglucy | 4.574 | 3.57E-07 | 8.03E-05 |
| ENSMUSG00000020963 | Tshr | 4.597 | 2.76E-03 | 4.76E-02 |
| ENSMUSG00000035948 | Acss3 | 4.611 | 8.64E-04 | 2.24E-02 |
| ENSMUSG00000027656 | Wisp2 | 4.623 | 1.65E-03 | 3.37E-02 |
| ENSMUSG00000030474 | Siglece | 4.628 | 1.89E-04 | 7.99E-03 |
| ENSMUSG00000086993 | Rsf1os2 | 4.637 | 1.17E-03 | 2.67E-02 |
| ENSMUSG00000041193 | Pla2g5 | 4.721 | 1.37E-03 | 2.98E-02 |
| ENSMUSG00000023153 | Tmem52 | 4.732 | 2.51E-03 | 4.48E-02 |
| ENSMUSG00000047591 | Mafa | 4.747 | 1.26E-03 | 2.81E-02 |
| ENSMUSG00000055632 | Hmcn2 | 4.762 | 2.39E-06 | 3.17E-04 |
| ENSMUSG00000085419 | Gm11734 | 4.796 | 7.16E-05 | 3.96E-03 |
| ENSMUSG00000084942 | 7330404K18 | 4.798 | 1.44E-03 | 3.04E-02 |
| ENSMUSG00000081670 | Gm15697 | 4.816 | 1.86E-03 | 3.67E-02 |
| ENSMUSG00000054763 | Defb42 | 4.822 | 2.00E-04 | 8.34E-03 |

| | | | | |
|---|---|---|---|---|
| ENSMUSG00000092592 | Gm20449 | 4.824 | 2.49E-03 | 4.46E-02 |
| ENSMUSG00000112765 | 9430078K24 | 4.882 | 3.52E-04 | 1.21E-02 |
| ENSMUSG00000089671 | Gm16537 | 4.896 | 7.10E-05 | 3.93E-03 |
| ENSMUSG00000042429 | Adora1 | 4.908 | 4.92E-04 | 1.52E-02 |
| ENSMUSG00000079363 | Gbp4 | 4.925 | 8.73E-04 | 2.25E-02 |
| ENSMUSG00000065715 | Snord7 | 4.934 | 1.48E-04 | 6.74E-03 |
| ENSMUSG00000028927 | Padi2 | 4.937 | 2.02E-03 | 3.86E-02 |
| ENSMUSG00000035916 | Ptprq | 4.939 | 4.87E-04 | 1.51E-02 |
| ENSMUSG00000071531 | Gprin2 | 4.945 | 7.29E-04 | 2.00E-02 |
| ENSMUSG00000054418 | 2900041M22 | 5.039 | 5.69E-04 | 1.69E-02 |
| ENSMUSG00000097459 | Gm26895 | 5.041 | 6.89E-04 | 1.92E-02 |
| ENSMUSG00000049625 | Tifab | 5.059 | 2.55E-03 | 4.52E-02 |
| ENSMUSG00000111329 | A830035O19 | 5.064 | 4.47E-04 | 1.42E-02 |
| ENSMUSG00000097399 | Gm26555 | 5.079 | 9.14E-04 | 2.31E-02 |
| ENSMUSG00000065952 | C330021F23 | 5.084 | 4.18E-04 | 1.35E-02 |
| ENSMUSG00000113771 | Gm36529 | 5.119 | 7.88E-04 | 2.11E-02 |
| ENSMUSG00000025329 | Padi1 | 5.122 | 2.20E-03 | 4.09E-02 |
| ENSMUSG00000107876 | Gm43936 | 5.125 | 2.31E-04 | 9.08E-03 |
| ENSMUSG00000084411 | Gm11510 | 5.128 | 6.67E-04 | 1.88E-02 |
| ENSMUSG00000030317 | Timp4 | 5.133 | 2.43E-05 | 1.80E-03 |
| ENSMUSG00000112833 | Gm36595 | 5.135 | 2.59E-03 | 4.58E-02 |
| ENSMUSG00000109770 | Gm30085 | 5.153 | 1.93E-03 | 3.75E-02 |
| ENSMUSG00000058966 | Fam57b | 5.171 | 2.98E-04 | 1.09E-02 |
| ENSMUSG00000027577 | Chrna4 | 5.177 | 9.64E-04 | 2.36E-02 |
| ENSMUSG00000108446 | Gm44997 | 5.180 | 1.08E-04 | 5.30E-03 |
| ENSMUSG00000073991 | Cnbd1 | 5.203 | 2.24E-03 | 4.15E-02 |
| ENSMUSG00000035686 | Thrsp | 5.218 | 4.35E-05 | 2.73E-03 |
| ENSMUSG00000087175 | Gm15133 | 5.225 | 6.72E-04 | 1.89E-02 |
| ENSMUSG00000103572 | Gm37115 | 5.233 | 8.25E-04 | 2.18E-02 |
| ENSMUSG00000034829 | Nxnl1 | 5.242 | 4.52E-04 | 1.43E-02 |
| ENSMUSG00000026222 | Sp100 | 5.246 | 1.86E-03 | 3.67E-02 |
| ENSMUSG00000048038 | Ccdc187 | 5.315 | 1.04E-04 | 5.19E-03 |
| ENSMUSG00000049555 | Tmie | 5.337 | 1.71E-04 | 7.46E-03 |
| ENSMUSG00000069814 | Ccdc92b | 5.339 | 2.30E-04 | 9.08E-03 |
| ENSMUSG00000111256 | Gm48094 | 5.358 | 2.50E-03 | 4.47E-02 |
| ENSMUSG00000016942 | Tmprss6 | 5.446 | 2.36E-04 | 9.19E-03 |
| ENSMUSG00000030217 | Art4 | 5.458 | 3.34E-05 | 2.29E-03 |
| ENSMUSG00000096997 | Gm26791 | 5.477 | 1.83E-03 | 3.63E-02 |
| ENSMUSG00000104011 | Gm32391 | 5.510 | 1.92E-04 | 8.06E-03 |
| ENSMUSG00000039264 | Gimap3 | 5.523 | 1.70E-03 | 3.45E-02 |
| ENSMUSG00000103062 | Gm37200 | 5.529 | 2.27E-03 | 4.20E-02 |
| ENSMUSG00000078998 | Bpifa6 | 5.538 | 1.82E-03 | 3.62E-02 |
| ENSMUSG00000113687 | AC123705.1 | 5.553 | 2.60E-03 | 4.59E-02 |
| ENSMUSG00000112220 | Gm36208 | 5.572 | 2.94E-05 | 2.08E-03 |
| ENSMUSG00000033182 | Kbtbd12 | 5.573 | 3.64E-04 | 1.24E-02 |
| ENSMUSG00000100198 | 1700030O20 | 5.615 | 6.29E-04 | 1.81E-02 |
| ENSMUSG00000105651 | 1700017M07 | 5.621 | 2.50E-03 | 4.46E-02 |

| | | | | |
|---|---|---|---|---|
| ENSMUSG00000116903 | AC161607.1 | 5.625 | 2.59E-04 | 9.83E-03 |
| ENSMUSG00000090164 | BC035044 | 5.626 | 3.87E-04 | 1.30E-02 |
| ENSMUSG00000038132 | Rbm24 | 5.651 | 2.20E-03 | 4.09E-02 |
| ENSMUSG00000087476 | Rap1gapos | 5.658 | 8.37E-04 | 2.20E-02 |
| ENSMUSG00000007279 | Scube2 | 5.670 | 4.97E-05 | 3.05E-03 |
| ENSMUSG00000078653 | Cntd1 | 5.680 | 3.24E-04 | 1.16E-02 |
| ENSMUSG00000021223 | Papln | 5.682 | 8.03E-04 | 2.14E-02 |
| ENSMUSG00000079013 | Serpina3j | 5.686 | 9.09E-06 | 8.37E-04 |
| ENSMUSG00000087612 | A230005M16 | 5.692 | 4.55E-06 | 5.23E-04 |
| ENSMUSG00000035000 | Dpp4 | 5.705 | 1.49E-03 | 3.13E-02 |
| ENSMUSG00000027220 | Syt13 | 5.705 | 9.80E-04 | 2.39E-02 |
| ENSMUSG00000048442 | Smim5 | 5.754 | 5.43E-04 | 1.64E-02 |
| ENSMUSG00000004359 | Spic | 5.781 | 4.16E-04 | 1.35E-02 |
| ENSMUSG00000018919 | Tm4sf5 | 5.791 | 6.60E-04 | 1.86E-02 |
| ENSMUSG00000109231 | Gm45737 | 5.797 | 1.64E-03 | 3.36E-02 |
| ENSMUSG00000006784 | Ttc25 | 5.807 | 3.61E-04 | 1.24E-02 |
| ENSMUSG00000085860 | 2410003L11 | 5.812 | 2.39E-04 | 9.23E-03 |
| ENSMUSG00000074623 | Gm826 | 5.812 | 4.28E-05 | 2.71E-03 |
| ENSMUSG00000079547 | H2-DMb1 | 5.828 | 1.10E-03 | 2.57E-02 |
| ENSMUSG00000060441 | Trim5 | 5.830 | 1.22E-04 | 5.84E-03 |
| ENSMUSG00000038233 | Fam198a | 5.856 | 2.38E-04 | 9.22E-03 |
| ENSMUSG00000030707 | Coro1a | 5.885 | 2.25E-03 | 4.17E-02 |
| ENSMUSG00000083827 | Gm15712 | 5.914 | 2.11E-03 | 3.99E-02 |
| ENSMUSG00000114150 | Gm46367 | 5.940 | 4.70E-04 | 1.47E-02 |
| ENSMUSG00000060791 | Gmfg | 5.941 | 7.31E-06 | 7.29E-04 |
| ENSMUSG00000102344 | 9430053O09 | 5.943 | 1.14E-03 | 2.63E-02 |
| ENSMUSG00000081058 | Hist2h3c2 | 5.970 | 5.04E-04 | 1.55E-02 |
| ENSMUSG00000102544 | Gm5103 | 5.970 | 4.64E-04 | 1.46E-02 |
| ENSMUSG00000069385 | Gm10267 | 5.997 | 1.06E-03 | 2.50E-02 |
| ENSMUSG00000085772 | D630024D03 | 6.009 | 1.50E-04 | 6.83E-03 |
| ENSMUSG00000097717 | 1700066J03I | 6.026 | 1.53E-04 | 6.90E-03 |
| ENSMUSG00000114939 | Gm48566 | 6.028 | 1.69E-03 | 3.44E-02 |
| ENSMUSG00000113974 | 4930442G10 | 6.050 | 2.40E-04 | 9.27E-03 |
| ENSMUSG00000002190 | Clgn | 6.093 | 4.29E-04 | 1.38E-02 |
| ENSMUSG00000024164 | C3 | 6.116 | 5.38E-04 | 1.63E-02 |
| ENSMUSG00000067704 | Wfdc13 | 6.130 | 1.34E-03 | 2.95E-02 |
| ENSMUSG00000035208 | Slfn8 | 6.163 | 2.29E-03 | 4.22E-02 |
| ENSMUSG00000048231 | H2-M10.4 | 6.169 | 4.23E-04 | 1.37E-02 |
| ENSMUSG00000043953 | Ccrl2 | 6.172 | 1.00E-03 | 2.43E-02 |
| ENSMUSG00000031089 | Slc6a14 | 6.205 | 7.01E-04 | 1.94E-02 |
| ENSMUSG00000035594 | Chrna5 | 6.230 | 3.90E-04 | 1.30E-02 |
| ENSMUSG00000072852 | 2310040G07 | 6.273 | 1.49E-04 | 6.79E-03 |
| ENSMUSG00000029228 | Lnx1 | 6.277 | 2.19E-04 | 8.81E-03 |
| ENSMUSG00000100815 | Gm29112 | 6.292 | 9.03E-04 | 2.30E-02 |
| ENSMUSG00000116288 | AC116487.2 | 6.403 | 1.82E-04 | 7.80E-03 |
| ENSMUSG00000108750 | Gm44750 | 6.549 | 1.86E-03 | 3.67E-02 |
| ENSMUSG00000062488 | Ifit3b | 6.555 | 1.06E-03 | 2.50E-02 |

| | | | | |
|---|---|---|---|---|
| ENSMUSG00000004892 | Bcan | 6.578 | 2.10E-07 | 5.27E-05 |
| ENSMUSG00000068463 | B630019A1[illegible] | 6.636 | 8.18E-05 | 4.36E-03 |
| ENSMUSG00000087132 | A930001C0[illegible] | 6.647 | 4.29E-07 | 9.08E-05 |
| ENSMUSG00000059625 | Sohlh1 | 6.657 | 3.00E-05 | 2.10E-03 |
| ENSMUSG00000041550 | Serpina5 | 6.704 | 7.40E-04 | 2.01E-02 |
| ENSMUSG00000029343 | Crybb1 | 6.750 | 3.74E-05 | 2.51E-03 |
| ENSMUSG00000047085 | Lrrc4b | 6.944 | 2.42E-05 | 1.80E-03 |
| ENSMUSG00000030732 | Chrdl2 | 7.003 | 2.91E-04 | 1.07E-02 |
| ENSMUSG00000104109 | Gm30292 | 7.038 | 1.14E-05 | 1.01E-03 |
| ENSMUSG00000037188 | Grhl3 | 7.097 | 2.92E-05 | 2.07E-03 |
| ENSMUSG00000090667 | Gm765 | 7.140 | 3.42E-04 | 1.20E-02 |
| ENSMUSG00000024245 | Tmem178 | 7.179 | 3.04E-09 | 1.79E-06 |
| ENSMUSG00000050700 | Emilin3 | 7.262 | 7.26E-05 | 3.99E-03 |
| ENSMUSG00000047343 | Mettl21c | 7.671 | 7.25E-06 | 7.26E-04 |
| ENSMUSG00000002324 | Rec8 | 7.790 | 8.74E-06 | 8.15E-04 |
| ENSMUSG00000085121 | 5730437C11 | 7.964 | 2.06E-06 | 2.86E-04 |
| ENSMUSG00000020928 | Higd1b | 17.753 | 2.64E-03 | 4.63E-02 |
| ENSMUSG00000070423 | Olfr558 | 18.332 | 1.80E-03 | 3.61E-02 |

**Supplementary Table S5.**

**List of DEGs between 3h-24h in 2%**

| Ensembl_ID | SYMBOL | og2FoldChang | pvalue | padj |
|---|---|---|---|---|
| ENSMUSG00000085183 | Wincr1 | -22.053 | 5.26E-28 | 1.02E-23 |
| ENSMUSG00000110163 | Gm45389 | -18.287 | 1.88E-03 | 3.87E-02 |
| ENSMUSG00000050578 | Mmp13 | -10.837 | 3.98E-10 | 2.65E-07 |
| ENSMUSG00000021680 | Crhbp | -10.495 | 3.66E-09 | 1.81E-06 |
| ENSMUSG00000115620 | Gm45924 | -8.719 | 9.48E-05 | 4.81E-03 |
| ENSMUSG00000050359 | Sprr1a | -8.526 | 7.92E-14 | 1.39E-10 |
| ENSMUSG00000020911 | Krt19 | -8.078 | 2.85E-04 | 1.04E-02 |
| ENSMUSG00000062345 | Serpinb2 | -8.021 | 1.23E-06 | 1.94E-04 |
| ENSMUSG00000047562 | Mmp10 | -7.771 | 3.01E-04 | 1.08E-02 |
| ENSMUSG00000021403 | Serpinb9b | -7.348 | 8.96E-13 | 1.44E-09 |
| ENSMUSG00000025582 | Nptx1 | -7.337 | 4.30E-04 | 1.37E-02 |
| ENSMUSG00000022296 | Baalc | -7.015 | 1.16E-05 | 9.65E-04 |
| ENSMUSG00000029819 | Npy | -6.829 | 5.84E-05 | 3.43E-03 |
| ENSMUSG00000109903 | Gm28710 | -6.728 | 3.10E-04 | 1.10E-02 |
| ENSMUSG00000024810 | Il33 | -6.671 | 4.17E-04 | 1.35E-02 |
| ENSMUSG00000112120 | Gm32255 | -6.607 | 9.34E-04 | 2.43E-02 |
| ENSMUSG00000021675 | F2rl2 | -6.603 | 2.05E-04 | 8.15E-03 |
| ENSMUSG00000111467 | Gm47775 | -6.472 | 2.83E-04 | 1.03E-02 |
| ENSMUSG00000029272 | Sult1e1 | -6.380 | 2.25E-03 | 4.43E-02 |
| ENSMUSG00000041552 | Ptchd1 | -6.373 | 1.74E-03 | 3.68E-02 |
| ENSMUSG00000044626 | Liph | -6.317 | 1.13E-04 | 5.40E-03 |
| ENSMUSG00000029304 | Spp1 | -5.985 | 6.48E-09 | 2.91E-06 |
| ENSMUSG00000029378 | Areg | -5.911 | 1.89E-03 | 3.87E-02 |
| ENSMUSG00000031297 | Slc7a3 | -5.791 | 5.43E-11 | 5.00E-08 |
| ENSMUSG00000026535 | Ifi202b | -5.760 | 1.76E-07 | 3.96E-05 |
| ENSMUSG00000026981 | Il1rn | -5.715 | 6.36E-05 | 3.61E-03 |
| ENSMUSG00000024912 | Fosl1 | -5.704 | 1.43E-07 | 3.47E-05 |
| ENSMUSG00000010797 | Wnt2 | -5.529 | 7.57E-05 | 4.12E-03 |
| ENSMUSG00000023905 | Tnfrsf12a | -5.512 | 1.13E-22 | 1.09E-18 |
| ENSMUSG00000019997 | Ctgf | -5.485 | 3.17E-14 | 6.12E-11 |
| ENSMUSG00000099974 | Bcl2a1d | -5.460 | 3.40E-04 | 1.17E-02 |
| ENSMUSG00000050335 | Lgals3 | -4.976 | 2.06E-14 | 4.97E-11 |
| ENSMUSG00000028172 | Tacr3 | -4.913 | 1.55E-03 | 3.44E-02 |
| ENSMUSG00000048482 | Bdnf | -4.881 | 1.69E-06 | 2.51E-04 |
| ENSMUSG00000029334 | Prkg2 | -4.816 | 2.45E-12 | 3.16E-09 |
| ENSMUSG00000105528 | Gm43519 | -4.781 | 8.48E-04 | 2.26E-02 |
| ENSMUSG00000041660 | Bbox1 | -4.720 | 8.28E-04 | 2.21E-02 |
| ENSMUSG00000115009 | G930009F23 | -4.716 | 1.18E-03 | 2.83E-02 |
| ENSMUSG00000063531 | Sema3e | -4.683 | 4.29E-04 | 1.37E-02 |
| ENSMUSG00000033730 | Egr3 | -4.679 | 6.88E-04 | 1.94E-02 |
| ENSMUSG00000015652 | Steap1 | -4.663 | 1.15E-12 | 1.71E-09 |
| ENSMUSG00000074199 | Krtdap | -4.603 | 4.17E-05 | 2.62E-03 |

| | | | | |
|---|---|---|---|---|
| ENSMUSG00000037946 | Fgd3 | -4.450 | 7.32E-06 | 6.97E-04 |
| ENSMUSG00000024640 | Psat1 | -4.404 | 4.81E-15 | 1.33E-11 |
| ENSMUSG00000032487 | Ptgs2 | -4.396 | 5.09E-04 | 1.55E-02 |
| ENSMUSG00000005148 | Klf5 | -4.393 | 1.63E-05 | 1.28E-03 |
| ENSMUSG00000055407 | Map6 | -4.354 | 1.25E-04 | 5.63E-03 |
| ENSMUSG00000029752 | Asns | -4.296 | 2.58E-14 | 5.53E-11 |
| ENSMUSG00000047171 | Helt | -4.277 | 2.70E-04 | 9.97E-03 |
| ENSMUSG00000032715 | Trib3 | -4.270 | 3.74E-08 | 1.25E-05 |
| ENSMUSG00000023043 | Krt18 | -4.259 | 4.04E-09 | 1.95E-06 |
| ENSMUSG00000021835 | Bmp4 | -4.259 | 1.46E-12 | 2.01E-09 |
| ENSMUSG00000042734 | Ttc9 | -4.060 | 9.48E-04 | 2.46E-02 |
| ENSMUSG00000028128 | F3 | -4.056 | 6.22E-08 | 1.79E-05 |
| ENSMUSG00000006445 | Epha2 | -4.041 | 1.73E-03 | 3.67E-02 |
| ENSMUSG00000046223 | Plaur | -4.031 | 1.28E-08 | 5.17E-06 |
| ENSMUSG00000115243 | Gm5207 | -4.015 | 3.00E-05 | 2.05E-03 |
| ENSMUSG00000007655 | Cav1 | -3.994 | 6.69E-11 | 5.87E-08 |
| ENSMUSG00000089812 | Gm15867 | -3.972 | 2.72E-05 | 1.90E-03 |
| ENSMUSG00000027313 | Chac1 | -3.963 | 7.08E-08 | 1.95E-05 |
| ENSMUSG00000029161 | Cgref1 | -3.941 | 4.37E-09 | 2.06E-06 |
| ENSMUSG00000023046 | Igfbp6 | -3.938 | 4.25E-18 | 2.74E-14 |
| ENSMUSG00000074813 | Gm14005 | -3.911 | 2.92E-05 | 2.00E-03 |
| ENSMUSG00000024907 | Gal | -3.880 | 2.67E-06 | 3.56E-04 |
| ENSMUSG00000031530 | Dusp4 | -3.822 | 9.00E-05 | 4.65E-03 |
| ENSMUSG00000028195 | Cyr61 | -3.802 | 2.29E-10 | 1.77E-07 |
| ENSMUSG00000003541 | Ier3 | -3.782 | 1.68E-11 | 1.80E-08 |
| ENSMUSG00000001021 | S100a3 | -3.736 | 3.09E-05 | 2.08E-03 |
| ENSMUSG00000027562 | Car2 | -3.732 | 1.08E-04 | 5.25E-03 |
| ENSMUSG00000078249 | Hmga1b | -3.727 | 2.73E-08 | 9.78E-06 |
| ENSMUSG00000022686 | B3gnt5 | -3.670 | 2.10E-03 | 4.20E-02 |
| ENSMUSG00000022805 | Maats1 | -3.655 | 1.04E-03 | 2.62E-02 |
| ENSMUSG00000028776 | Tinagl1 | -3.603 | 2.38E-09 | 1.25E-06 |
| ENSMUSG00000026479 | Lamc2 | -3.544 | 7.35E-06 | 6.97E-04 |
| ENSMUSG00000028480 | Glipr2 | -3.525 | 1.68E-03 | 3.59E-02 |
| ENSMUSG00000037855 | Zfp365 | -3.517 | 7.53E-07 | 1.30E-04 |
| ENSMUSG00000051111 | Sv2c | -3.494 | 1.71E-04 | 7.20E-03 |
| ENSMUSG00000020256 | Aldh1l2 | -3.473 | 2.26E-04 | 8.70E-03 |
| ENSMUSG00000037868 | Egr2 | -3.462 | 1.66E-03 | 3.58E-02 |
| ENSMUSG00000021822 | Plau | -3.360 | 6.29E-05 | 3.59E-03 |
| ENSMUSG00000005124 | Wisp1 | -3.310 | 1.78E-06 | 2.57E-04 |
| ENSMUSG00000019960 | Dusp6 | -3.299 | 9.27E-07 | 1.53E-04 |
| ENSMUSG00000030854 | Ptpn5 | -3.264 | 4.40E-04 | 1.39E-02 |
| ENSMUSG00000030827 | Fgf21 | -3.240 | 1.94E-03 | 3.94E-02 |
| ENSMUSG00000074657 | Kif5a | -3.230 | 1.13E-03 | 2.77E-02 |
| ENSMUSG00000046711 | Hmga1 | -3.222 | 3.71E-06 | 4.19E-04 |
| ENSMUSG00000017144 | Rnd3 | -3.212 | 5.39E-07 | 1.00E-04 |
| ENSMUSG00000042256 | Ptchd4 | -3.208 | 1.05E-03 | 2.64E-02 |
| ENSMUSG00000000058 | Cav2 | -3.176 | 4.61E-08 | 1.42E-05 |

| | | | | |
|---|---|---|---|---|
| ENSMUSG00000110126 | Gm9347 | -3.162 | 4.48E-06 | 4.81E-04 |
| ENSMUSG00000034271 | Jdp2 | -3.098 | 7.32E-06 | 6.97E-04 |
| ENSMUSG00000032068 | Plet1 | -3.077 | 2.62E-03 | 4.92E-02 |
| ENSMUSG00000057604 | Lmcd1 | -3.050 | 1.89E-05 | 1.42E-03 |
| ENSMUSG00000029446 | Psph | -2.990 | 1.79E-16 | 6.91E-13 |
| ENSMUSG00000025140 | Pycr1 | -2.976 | 6.06E-04 | 1.77E-02 |
| ENSMUSG00000030717 | Nupr1 | -2.905 | 3.49E-04 | 1.20E-02 |
| ENSMUSG00000113769 | 5033406O09 | -2.877 | 5.26E-06 | 5.50E-04 |
| ENSMUSG00000062661 | Ncs1 | -2.857 | 2.73E-06 | 3.61E-04 |
| ENSMUSG00000031434 | Morc4 | -2.779 | 4.66E-05 | 2.84E-03 |
| ENSMUSG00000026547 | Tagln2 | -2.777 | 6.84E-04 | 1.94E-02 |
| ENSMUSG00000115505 | Gm9247 | -2.751 | 2.46E-03 | 4.71E-02 |
| ENSMUSG00000003283 | Hck | -2.714 | 2.66E-03 | 4.98E-02 |
| ENSMUSG00000028811 | Yars | -2.711 | 8.32E-07 | 1.40E-04 |
| ENSMUSG00000042549 | Map2k3os | -2.706 | 1.29E-03 | 3.04E-02 |
| ENSMUSG00000028680 | Plk3 | -2.689 | 7.13E-05 | 3.95E-03 |
| ENSMUSG00000037960 | Card19 | -2.682 | 1.51E-08 | 5.94E-06 |
| ENSMUSG00000020142 | Slc1a4 | -2.661 | 1.15E-03 | 2.80E-02 |
| ENSMUSG00000039126 | Prune2 | -2.653 | 1.83E-05 | 1.39E-03 |
| ENSMUSG00000015312 | Gadd45b | -2.627 | 4.50E-07 | 8.98E-05 |
| ENSMUSG00000083512 | Gm12749 | -2.615 | 1.98E-03 | 4.02E-02 |
| ENSMUSG00000024587 | Nars | -2.610 | 1.65E-07 | 3.79E-05 |
| ENSMUSG00000029777 | Gars | -2.568 | 1.50E-09 | 9.15E-07 |
| ENSMUSG00000037060 | Cavin3 | -2.556 | 1.38E-03 | 3.18E-02 |
| ENSMUSG00000036856 | Wnt4 | -2.541 | 2.60E-03 | 4.90E-02 |
| ENSMUSG00000026784 | Pdss1 | -2.535 | 3.79E-12 | 4.58E-09 |
| ENSMUSG00000031574 | Star | -2.491 | 6.46E-06 | 6.33E-04 |
| ENSMUSG00000006442 | Srm | -2.471 | 4.47E-07 | 8.98E-05 |
| ENSMUSG00000052688 | Rab7b | -2.461 | 4.62E-07 | 9.11E-05 |
| ENSMUSG00000053746 | Ptrh1 | -2.450 | 3.48E-06 | 4.07E-04 |
| ENSMUSG00000073433 | Arhgdig | -2.423 | 6.07E-07 | 1.10E-04 |
| ENSMUSG00000022241 | Tars | -2.422 | 3.61E-06 | 4.13E-04 |
| ENSMUSG00000005667 | Mthfd2 | -2.414 | 9.69E-07 | 1.59E-04 |
| ENSMUSG00000091898 | Tnnc1 | -2.413 | 1.70E-06 | 2.51E-04 |
| ENSMUSG00000010755 | Cars | -2.412 | 3.62E-10 | 2.50E-07 |
| ENSMUSG00000025511 | Tspan4 | -2.400 | 4.67E-08 | 1.42E-05 |
| ENSMUSG00000082674 | Gm11914 | -2.390 | 1.07E-03 | 2.67E-02 |
| ENSMUSG00000011179 | Odc1 | -2.338 | 1.40E-10 | 1.13E-07 |
| ENSMUSG00000101431 | Gm7901 | -2.334 | 4.16E-04 | 1.35E-02 |
| ENSMUSG00000026749 | Nek6 | -2.330 | 8.78E-06 | 8.04E-04 |
| ENSMUSG00000055737 | Ghr | -2.329 | 9.24E-06 | 8.27E-04 |
| ENSMUSG00000060950 | Trmt61a | -2.329 | 1.81E-04 | 7.48E-03 |
| ENSMUSG00000019970 | Sgk1 | -2.325 | 6.25E-04 | 1.80E-02 |
| ENSMUSG00000031490 | Eif4ebp1 | -2.322 | 1.18E-05 | 9.74E-04 |
| ENSMUSG00000053398 | Phgdh | -2.321 | 5.32E-07 | 9.98E-05 |
| ENSMUSG00000027610 | Gss | -2.317 | 1.61E-07 | 3.76E-05 |
| ENSMUSG00000025007 | Aldh18a1 | -2.308 | 1.42E-06 | 2.19E-04 |

| | | | | |
|---|---|---|---|---|
| ENSMUSG00000001025 | S100a6 | -2.297 | 2.65E-03 | 4.96E-02 |
| ENSMUSG00000028063 | Lmna | -2.247 | 1.28E-04 | 5.73E-03 |
| ENSMUSG00000034765 | Dusp5 | -2.213 | 4.92E-06 | 5.17E-04 |
| ENSMUSG00000038539 | Atf5 | -2.197 | 1.08E-05 | 9.16E-04 |
| ENSMUSG00000032802 | Srxn1 | -2.188 | 5.89E-04 | 1.74E-02 |
| ENSMUSG00000018849 | Wwc1 | -2.167 | 1.37E-03 | 3.16E-02 |
| ENSMUSG00000050222 | Il17d | -2.164 | 7.53E-04 | 2.06E-02 |
| ENSMUSG00000038894 | Irs2 | -2.163 | 9.62E-05 | 4.84E-03 |
| ENSMUSG00000028069 | Gpatch4 | -2.155 | 4.60E-08 | 1.42E-05 |
| ENSMUSG00000083465 | Gm11652 | -2.148 | 9.03E-05 | 4.65E-03 |
| ENSMUSG00000042743 | Sgtb | -2.137 | 1.65E-03 | 3.58E-02 |
| ENSMUSG00000068739 | Sars | -2.130 | 3.33E-06 | 3.97E-04 |
| ENSMUSG00000025403 | Shmt2 | -2.047 | 9.18E-06 | 8.27E-04 |
| ENSMUSG00000025287 | Acot9 | -2.032 | 3.40E-06 | 4.00E-04 |
| ENSMUSG00000098274 | Rpl24 | -2.029 | 3.54E-04 | 1.21E-02 |
| ENSMUSG00000042515 | Mum1l1 | -2.025 | 2.35E-06 | 3.21E-04 |
| ENSMUSG00000041506 | Rrp9 | -2.002 | 3.31E-06 | 3.97E-04 |
| ENSMUSG00000025869 | Nop16 | -2.002 | 3.27E-06 | 3.97E-04 |
| ENSMUSG00000002343 | Armc6 | -1.999 | 8.49E-04 | 2.26E-02 |
| ENSMUSG00000032231 | Anxa2 | -1.982 | 4.07E-06 | 4.50E-04 |
| ENSMUSG00000045763 | Basp1 | -1.979 | 8.78E-05 | 4.61E-03 |
| ENSMUSG00000050017 | Pitpnb | -1.968 | 3.19E-04 | 1.11E-02 |
| ENSMUSG00000038335 | Tsr1 | -1.957 | 1.51E-09 | 9.15E-07 |
| ENSMUSG00000020205 | Phlda1 | -1.954 | 1.43E-06 | 2.19E-04 |
| ENSMUSG00000037601 | Nme1 | -1.949 | 5.86E-07 | 1.07E-04 |
| ENSMUSG00000038279 | Nop2 | -1.945 | 1.58E-07 | 3.72E-05 |
| ENSMUSG00000003348 | Mob3a | -1.928 | 1.78E-03 | 3.74E-02 |
| ENSMUSG00000056501 | Cebpb | -1.927 | 1.74E-03 | 3.68E-02 |
| ENSMUSG00000041774 | Ydjc | -1.923 | 3.95E-04 | 1.31E-02 |
| ENSMUSG00000028495 | Rps6 | -1.921 | 1.23E-05 | 1.00E-03 |
| ENSMUSG00000053560 | Ier2 | -1.920 | 1.68E-03 | 3.59E-02 |
| ENSMUSG00000029171 | Pgm1 | -1.909 | 1.76E-04 | 7.34E-03 |
| ENSMUSG00000020303 | Stc2 | -1.908 | 1.18E-04 | 5.51E-03 |
| ENSMUSG00000074220 | Zfp382 | -1.896 | 9.55E-04 | 2.47E-02 |
| ENSMUSG00000028896 | Rcc1 | -1.887 | 4.09E-05 | 2.58E-03 |
| ENSMUSG00000018932 | Map2k3 | -1.887 | 3.07E-04 | 1.09E-02 |
| ENSMUSG00000000561 | Wdr77 | -1.885 | 1.82E-05 | 1.39E-03 |
| ENSMUSG00000055044 | Pdlim1 | -1.882 | 1.29E-03 | 3.03E-02 |
| ENSMUSG00000085156 | Snhg15 | -1.877 | 3.59E-04 | 1.22E-02 |
| ENSMUSG00000063229 | Ldha | -1.872 | 1.20E-04 | 5.57E-03 |
| ENSMUSG00000085666 | Gm9855 | -1.871 | 2.44E-03 | 4.68E-02 |
| ENSMUSG00000021831 | Ero1l | -1.847 | 2.99E-05 | 2.04E-03 |
| ENSMUSG00000048612 | Myof | -1.825 | 2.80E-04 | 1.02E-02 |
| ENSMUSG00000007041 | Clic1 | -1.811 | 2.55E-05 | 1.84E-03 |
| ENSMUSG00000053801 | Grwd1 | -1.807 | 2.01E-06 | 2.83E-04 |
| ENSMUSG00000047565 | Acot10 | -1.807 | 1.38E-03 | 3.19E-02 |
| ENSMUSG00000039405 | Prss23 | -1.791 | 1.87E-03 | 3.86E-02 |

| | | | | |
|---|---|---|---|---|
| ENSMUSG00000027405 | Nop56 | -1.785 | 4.86E-08 | 1.42E-05 |
| ENSMUSG00000010067 | Rassf1 | -1.779 | 9.09E-04 | 2.38E-02 |
| ENSMUSG00000024037 | Wdr4 | -1.768 | 2.23E-04 | 8.66E-03 |
| ENSMUSG00000063316 | Rpl27 | -1.763 | 4.07E-08 | 1.33E-05 |
| ENSMUSG00000051223 | Bzw1 | -1.761 | 3.85E-05 | 2.46E-03 |
| ENSMUSG00000037465 | Klf10 | -1.759 | 1.51E-03 | 3.37E-02 |
| ENSMUSG00000101188 | Eif4a-ps4 | -1.754 | 5.30E-04 | 1.60E-02 |
| ENSMUSG00000043192 | Gm1840 | -1.744 | 7.56E-04 | 2.06E-02 |
| ENSMUSG00000002825 | Qtrt1 | -1.740 | 2.41E-04 | 9.19E-03 |
| ENSMUSG00000049957 | Ccdc137 | -1.736 | 7.83E-05 | 4.21E-03 |
| ENSMUSG00000032966 | Fkbp1a | -1.734 | 1.75E-04 | 7.33E-03 |
| ENSMUSG00000032051 | Fdx1 | -1.727 | 2.01E-03 | 4.07E-02 |
| ENSMUSG00000008450 | Nutf2 | -1.698 | 7.47E-07 | 1.30E-04 |
| ENSMUSG00000013089 | Etv5 | -1.692 | 2.28E-03 | 4.47E-02 |
| ENSMUSG00000000078 | Klf6 | -1.687 | 6.76E-05 | 3.78E-03 |
| ENSMUSG00000112903 | Gm5948 | -1.684 | 4.24E-04 | 1.36E-02 |
| ENSMUSG00000095567 | Noc2l | -1.669 | 3.69E-05 | 2.42E-03 |
| ENSMUSG00000078812 | Eif5a | -1.664 | 1.90E-04 | 7.77E-03 |
| ENSMUSG00000022184 | Fbxo4 | -1.656 | 1.81E-06 | 2.60E-04 |
| ENSMUSG00000052825 | Gm9892 | -1.655 | 5.45E-06 | 5.66E-04 |
| ENSMUSG00000001305 | Rrp15 | -1.649 | 2.81E-05 | 1.95E-03 |
| ENSMUSG00000024493 | Lars | -1.642 | 6.50E-07 | 1.16E-04 |
| ENSMUSG00000029500 | Pgam5 | -1.642 | 6.19E-05 | 3.57E-03 |
| ENSMUSG00000042406 | Atf4 | -1.639 | 7.61E-05 | 4.13E-03 |
| ENSMUSG00000028010 | Gar1 | -1.631 | 2.28E-06 | 3.12E-04 |
| ENSMUSG00000085385 | Snhg17 | -1.626 | 9.84E-06 | 8.61E-04 |
| ENSMUSG00000052151 | Plpp2 | -1.615 | 2.06E-05 | 1.53E-03 |
| ENSMUSG00000074280 | Gm6166 | -1.612 | 2.09E-03 | 4.19E-02 |
| ENSMUSG00000003848 | Nob1 | -1.602 | 4.32E-04 | 1.38E-02 |
| ENSMUSG00000067161 | Gm5560 | -1.600 | 1.78E-03 | 3.73E-02 |
| ENSMUSG00000033706 | Smyd5 | -1.594 | 2.08E-04 | 8.19E-03 |
| ENSMUSG00000064264 | Zfp428 | -1.591 | 1.19E-07 | 2.99E-05 |
| ENSMUSG00000024999 | Noc3l | -1.580 | 2.67E-04 | 9.93E-03 |
| ENSMUSG00000028466 | Creb3 | -1.561 | 9.62E-04 | 2.48E-02 |
| ENSMUSG00000046865 | Fbl | -1.558 | 2.07E-04 | 8.19E-03 |
| ENSMUSG00000000916 | Nsun5 | -1.557 | 4.34E-04 | 1.38E-02 |
| ENSMUSG00000040354 | Mars | -1.551 | 7.84E-05 | 4.21E-03 |
| ENSMUSG00000022554 | Hgh1 | -1.550 | 2.61E-07 | 5.49E-05 |
| ENSMUSG00000031960 | Aars | -1.547 | 3.97E-05 | 2.52E-03 |
| ENSMUSG00000048007 | Timm8a1 | -1.542 | 2.65E-04 | 9.88E-03 |
| ENSMUSG00000090243 | Gm16103 | -1.541 | 2.94E-04 | 1.05E-02 |
| ENSMUSG00000028729 | Ebna1bp2 | -1.540 | 3.92E-07 | 8.05E-05 |
| ENSMUSG00000005481 | Ddx39 | -1.540 | 1.07E-05 | 9.11E-04 |
| ENSMUSG00000048261 | Gm4879 | -1.531 | 1.07E-03 | 2.66E-02 |
| ENSMUSG00000021595 | Nsun2 | -1.529 | 2.25E-04 | 8.70E-03 |
| ENSMUSG00000026234 | Ncl | -1.522 | 1.10E-07 | 2.81E-05 |
| ENSMUSG00000029430 | Ran | -1.519 | 2.19E-08 | 8.29E-06 |

| | | | | |
|---|---|---|---|---|
| ENSMUSG00000028741 | Mrto4 | -1.510 | 9.00E-07 | 1.50E-04 |
| ENSMUSG00000051557 | Pusl1 | -1.508 | 1.17E-03 | 2.82E-02 |
| ENSMUSG00000057561 | Eif1a | -1.505 | 1.10E-05 | 9.34E-04 |
| ENSMUSG00000040463 | Mybbp1a | -1.504 | 1.82E-07 | 4.04E-05 |
| ENSMUSG00000029610 | Aimp2 | -1.498 | 8.28E-06 | 7.71E-04 |
| ENSMUSG00000032078 | Zpr1 | -1.497 | 4.65E-04 | 1.45E-02 |
| ENSMUSG00000024841 | Eif1ad | -1.496 | 2.42E-06 | 3.27E-04 |
| ENSMUSG00000026020 | Nop58 | -1.486 | 4.72E-09 | 2.17E-06 |
| ENSMUSG00000030079 | Ruvbl1 | -1.482 | 1.16E-05 | 9.65E-04 |
| ENSMUSG00000041360 | Pum3 | -1.478 | 7.05E-04 | 1.98E-02 |
| ENSMUSG00000031578 | Mak16 | -1.475 | 1.01E-05 | 8.69E-04 |
| ENSMUSG00000063524 | Eno1 | -1.475 | 5.40E-05 | 3.25E-03 |
| ENSMUSG00000018446 | C1qbp | -1.475 | 8.15E-08 | 2.19E-05 |
| ENSMUSG00000026019 | Wdr12 | -1.472 | 1.07E-04 | 5.23E-03 |
| ENSMUSG00000026670 | Uap1 | -1.469 | 3.34E-06 | 3.97E-04 |
| ENSMUSG00000006732 | Mettl1 | -1.468 | 9.71E-06 | 8.57E-04 |
| ENSMUSG00000026603 | Smyd2 | -1.463 | 1.09E-03 | 2.70E-02 |
| ENSMUSG00000040675 | Mthfd1l | -1.459 | 4.21E-08 | 1.36E-05 |
| ENSMUSG00000024312 | Wdr46 | -1.458 | 2.12E-03 | 4.23E-02 |
| ENSMUSG00000041057 | Wdr43 | -1.450 | 2.86E-06 | 3.72E-04 |
| ENSMUSG00000025485 | Ric8a | -1.449 | 1.12E-03 | 2.76E-02 |
| ENSMUSG00000042215 | Bag2 | -1.445 | 9.62E-06 | 8.53E-04 |
| ENSMUSG00000001380 | Hars | -1.445 | 3.05E-04 | 1.09E-02 |
| ENSMUSG00000038046 | Mrm3 | -1.444 | 3.17E-06 | 3.95E-04 |
| ENSMUSG00000058355 | Abce1 | -1.442 | 2.52E-04 | 9.48E-03 |
| ENSMUSG00000001627 | Ifrd1 | -1.440 | 1.50E-04 | 6.49E-03 |
| ENSMUSG00000028982 | Slc25a33 | -1.439 | 5.60E-05 | 3.32E-03 |
| ENSMUSG00000027944 | Hax1 | -1.438 | 2.06E-07 | 4.47E-05 |
| ENSMUSG00000062867 | Impdh2 | -1.436 | 1.10E-03 | 2.72E-02 |
| ENSMUSG00000039356 | Exosc2 | -1.428 | 9.87E-05 | 4.94E-03 |
| ENSMUSG00000031432 | Prps1 | -1.422 | 2.97E-04 | 1.07E-02 |
| ENSMUSG00000055612 | Cdca7 | -1.418 | 3.75E-05 | 2.43E-03 |
| ENSMUSG00000031146 | Plp2 | -1.416 | 2.60E-04 | 9.74E-03 |
| ENSMUSG00000022863 | Btg3 | -1.409 | 3.98E-04 | 1.31E-02 |
| ENSMUSG00000028318 | Polr1e | -1.405 | 1.45E-04 | 6.32E-03 |
| ENSMUSG00000029447 | Cct6a | -1.381 | 2.82E-05 | 1.95E-03 |
| ENSMUSG00000020116 | Pno1 | -1.380 | 1.64E-06 | 2.46E-04 |
| ENSMUSG00000026806 | Ddx31 | -1.379 | 3.10E-05 | 2.08E-03 |
| ENSMUSG00000067367 | Lyar | -1.375 | 2.03E-05 | 1.51E-03 |
| ENSMUSG00000025981 | Coq10b | -1.373 | 2.09E-04 | 8.21E-03 |
| ENSMUSG00000020691 | Mettl2 | -1.370 | 5.31E-04 | 1.60E-02 |
| ENSMUSG00000025980 | Hspd1 | -1.368 | 8.81E-12 | 1.00E-08 |
| ENSMUSG00000027395 | Polr1b | -1.367 | 1.13E-03 | 2.77E-02 |
| ENSMUSG00000029036 | Atad3a | -1.364 | 1.21E-04 | 5.57E-03 |
| ENSMUSG00000041488 | Stx3 | -1.363 | 7.15E-06 | 6.87E-04 |
| ENSMUSG00000042354 | Gnl3 | -1.362 | 4.88E-04 | 1.50E-02 |
| ENSMUSG00000020089 | Ppa1 | -1.360 | 9.61E-09 | 4.04E-06 |

| | | | | |
|---|---|---|---|---|
| ENSMUSG00000040158 | Tax1bp3 | -1.351 | 2.10E-03 | 4.20E-02 |
| ENSMUSG00000021149 | Gtpbp4 | -1.350 | 4.74E-08 | 1.42E-05 |
| ENSMUSG00000070284 | Gmppb | -1.350 | 2.28E-03 | 4.47E-02 |
| ENSMUSG00000031568 | Rwdd4a | -1.350 | 1.16E-04 | 5.48E-03 |
| ENSMUSG00000036693 | Nop14 | -1.347 | 7.03E-08 | 1.95E-05 |
| ENSMUSG00000062937 | Mtap | -1.345 | 7.26E-04 | 2.02E-02 |
| ENSMUSG00000038550 | Ciart | -1.341 | 3.01E-04 | 1.08E-02 |
| ENSMUSG00000071359 | Tbpl1 | -1.341 | 5.72E-04 | 1.70E-02 |
| ENSMUSG00000038510 | Rpf2 | -1.337 | 4.82E-04 | 1.49E-02 |
| ENSMUSG00000000339 | Rtca | -1.336 | 3.15E-04 | 1.11E-02 |
| ENSMUSG00000023988 | Bysl | -1.335 | 1.50E-03 | 3.36E-02 |
| ENSMUSG00000054836 | Elp6 | -1.329 | 2.23E-03 | 4.39E-02 |
| ENSMUSG00000020358 | Hnrnpab | -1.329 | 1.26E-04 | 5.67E-03 |
| ENSMUSG00000031352 | Hccs | -1.324 | 3.68E-04 | 1.24E-02 |
| ENSMUSG00000027804 | Ppid | -1.322 | 3.04E-05 | 2.06E-03 |
| ENSMUSG00000021282 | Eif5 | -1.318 | 2.87E-06 | 3.72E-04 |
| ENSMUSG00000025995 | Wdr75 | -1.315 | 6.04E-05 | 3.53E-03 |
| ENSMUSG00000025591 | Tma16 | -1.310 | 3.25E-06 | 3.97E-04 |
| ENSMUSG00000074656 | Eif2s2 | -1.310 | 1.84E-05 | 1.39E-03 |
| ENSMUSG00000038388 | Mpp6 | -1.308 | 2.39E-03 | 4.64E-02 |
| ENSMUSG00000026245 | Farsb | -1.306 | 1.41E-03 | 3.23E-02 |
| ENSMUSG00000031278 | Acsl4 | -1.306 | 3.55E-05 | 2.34E-03 |
| ENSMUSG00000027236 | Eif3j1 | -1.302 | 2.30E-07 | 4.93E-05 |
| ENSMUSG00000024360 | Etf1 | -1.302 | 1.78E-03 | 3.74E-02 |
| ENSMUSG00000097195 | Snhg5 | -1.297 | 1.72E-04 | 7.24E-03 |
| ENSMUSG00000062070 | Pgk1 | -1.295 | 5.09E-04 | 1.55E-02 |
| ENSMUSG00000004127 | Trmt10a | -1.294 | 1.87E-04 | 7.69E-03 |
| ENSMUSG00000020869 | Lrrc59 | -1.293 | 1.27E-03 | 3.00E-02 |
| ENSMUSG00000010095 | Slc3a2 | -1.287 | 3.79E-06 | 4.24E-04 |
| ENSMUSG00000024726 | Carnmt1 | -1.283 | 5.10E-04 | 1.55E-02 |
| ENSMUSG00000024359 | Hspa9 | -1.282 | 3.98E-07 | 8.09E-05 |
| ENSMUSG00000068039 | Tcp1 | -1.278 | 8.71E-08 | 2.31E-05 |
| ENSMUSG00000115219 | Eef1akmt4 | -1.268 | 5.87E-04 | 1.74E-02 |
| ENSMUSG00000056962 | Jmjd6 | -1.267 | 2.04E-04 | 8.13E-03 |
| ENSMUSG00000081999 | Gm13461 | -1.261 | 1.89E-04 | 7.75E-03 |
| ENSMUSG00000021694 | Ercc8 | -1.261 | 2.82E-05 | 1.95E-03 |
| ENSMUSG00000028683 | Eif2b3 | -1.260 | 9.19E-06 | 8.27E-04 |
| ENSMUSG00000007029 | Vars | -1.256 | 1.42E-04 | 6.23E-03 |
| ENSMUSG00000035530 | Eif1 | -1.251 | 1.68E-07 | 3.81E-05 |
| ENSMUSG00000020075 | Ddx21 | -1.248 | 4.72E-07 | 9.21E-05 |
| ENSMUSG00000031917 | Nip7 | -1.247 | 5.78E-06 | 5.88E-04 |
| ENSMUSG00000030057 | Cnbp | -1.246 | 6.22E-05 | 3.57E-03 |
| ENSMUSG00000061024 | Rrs1 | -1.244 | 1.13E-04 | 5.42E-03 |
| ENSMUSG00000108414 | Snhg1 | -1.240 | 2.08E-03 | 4.17E-02 |
| ENSMUSG00000029507 | Pus1 | -1.239 | 1.17E-03 | 2.82E-02 |
| ENSMUSG00000022234 | Cct5 | -1.237 | 1.31E-05 | 1.05E-03 |
| ENSMUSG00000028330 | Ncbp1 | -1.228 | 2.11E-04 | 8.25E-03 |

| | | | | |
|---|---|---|---|---|
| ENSMUSG00000024805 | Pcgf5 | -1.226 | 3.25E-04 | 1.13E-02 |
| ENSMUSG00000025264 | Tsr2 | -1.225 | 1.09E-04 | 5.29E-03 |
| ENSMUSG00000030007 | Cct7 | -1.224 | 1.21E-04 | 5.57E-03 |
| ENSMUSG00000026615 | Eprs | -1.218 | 1.05E-05 | 9.05E-04 |
| ENSMUSG00000063888 | Rpl7l1 | -1.217 | 1.70E-03 | 3.62E-02 |
| ENSMUSG00000026617 | Bpnt1 | -1.216 | 1.83E-05 | 1.39E-03 |
| ENSMUSG00000034343 | Ube2f | -1.209 | 9.86E-07 | 1.60E-04 |
| ENSMUSG00000020484 | Xbp1 | -1.208 | 3.67E-04 | 1.24E-02 |
| ENSMUSG00000001785 | Pwp1 | -1.207 | 2.56E-04 | 9.62E-03 |
| ENSMUSG00000038274 | Fau | -1.206 | 1.46E-07 | 3.49E-05 |
| ENSMUSG00000021116 | Eif2s1 | -1.204 | 1.76E-04 | 7.34E-03 |
| ENSMUSG00000046027 | Stard5 | -1.199 | 5.60E-04 | 1.67E-02 |
| ENSMUSG00000030888 | Rrp8 | -1.196 | 6.04E-04 | 1.77E-02 |
| ENSMUSG00000027613 | Eif6 | -1.195 | 1.66E-03 | 3.58E-02 |
| ENSMUSG00000024732 | Ccdc86 | -1.194 | 3.65E-04 | 1.23E-02 |
| ENSMUSG00000059796 | Eif4a1 | -1.190 | 1.83E-05 | 1.39E-03 |
| ENSMUSG00000034203 | Chchd4 | -1.190 | 1.23E-06 | 1.94E-04 |
| ENSMUSG00000034120 | Srsf2 | -1.188 | 4.10E-06 | 4.51E-04 |
| ENSMUSG00000041747 | Utp15 | -1.187 | 2.91E-06 | 3.74E-04 |
| ENSMUSG00000020648 | Dus4l | -1.187 | 8.60E-04 | 2.28E-02 |
| ENSMUSG00000023110 | Prmt5 | -1.184 | 1.13E-03 | 2.77E-02 |
| ENSMUSG00000041453 | Rpl21 | -1.183 | 7.22E-04 | 2.01E-02 |
| ENSMUSG00000070319 | Eif3g | -1.179 | 1.43E-07 | 3.47E-05 |
| ENSMUSG00000025134 | Alyref | -1.174 | 1.51E-05 | 1.19E-03 |
| ENSMUSG00000003868 | Ruvbl2 | -1.174 | 4.15E-04 | 1.35E-02 |
| ENSMUSG00000030738 | Eif3c | -1.166 | 1.28E-04 | 5.73E-03 |
| ENSMUSG00000014226 | Cacybp | -1.163 | 1.13E-03 | 2.77E-02 |
| ENSMUSG00000029229 | Chic2 | -1.155 | 4.16E-04 | 1.35E-02 |
| ENSMUSG00000016554 | Eif3d | -1.155 | 2.33E-05 | 1.70E-03 |
| ENSMUSG00000055723 | Rras2 | -1.146 | 1.90E-03 | 3.89E-02 |
| ENSMUSG00000022407 | Adsl | -1.134 | 9.92E-04 | 2.53E-02 |
| ENSMUSG00000015176 | Nolc1 | -1.132 | 1.37E-03 | 3.16E-02 |
| ENSMUSG00000025613 | Cct8 | -1.130 | 1.72E-06 | 2.51E-04 |
| ENSMUSG00000038975 | Rabggtb | -1.128 | 7.76E-05 | 4.20E-03 |
| ENSMUSG00000025858 | Get4 | -1.125 | 1.36E-03 | 3.16E-02 |
| ENSMUSG00000001707 | Eef1e1 | -1.123 | 7.23E-05 | 3.97E-03 |
| ENSMUSG00000063884 | Ptcd3 | -1.118 | 1.85E-05 | 1.40E-03 |
| ENSMUSG00000027669 | Gnb4 | -1.117 | 2.43E-03 | 4.67E-02 |
| ENSMUSG00000006699 | Cdc42 | -1.108 | 1.51E-03 | 3.37E-02 |
| ENSMUSG00000002881 | Nab1 | -1.107 | 2.30E-04 | 8.82E-03 |
| ENSMUSG00000024528 | Srfbp1 | -1.106 | 3.18E-04 | 1.11E-02 |
| ENSMUSG00000006599 | Gtf2h1 | -1.106 | 1.19E-03 | 2.85E-02 |
| ENSMUSG00000043424 | Eif3j2 | -1.102 | 8.41E-06 | 7.78E-04 |
| ENSMUSG00000001525 | Tubb5 | -1.101 | 3.80E-05 | 2.45E-03 |
| ENSMUSG00000028007 | Snx7 | -1.096 | 2.47E-03 | 4.72E-02 |
| ENSMUSG00000079435 | Rpl36a | -1.094 | 2.19E-03 | 4.32E-02 |
| ENSMUSG00000030264 | Thumpd3 | -1.093 | 4.87E-05 | 2.94E-03 |

| | | | | |
|---|---|---|---|---|
| ENSMUSG00000027808 | Serp1 | -1.091 | 4.89E-04 | 1.50E-02 |
| ENSMUSG00000063694 | Cycs | -1.090 | 1.80E-03 | 3.76E-02 |
| ENSMUSG00000016319 | Slc25a5 | -1.086 | 2.47E-04 | 9.35E-03 |
| ENSMUSG00000004264 | Phb2 | -1.079 | 1.69E-03 | 3.61E-02 |
| ENSMUSG00000013701 | Timm23 | -1.078 | 1.82E-05 | 1.39E-03 |
| ENSMUSG00000072235 | Tuba1a | -1.077 | 1.09E-03 | 2.70E-02 |
| ENSMUSG00000106988 | Tsg101-ps | -1.076 | 7.45E-04 | 2.05E-02 |
| ENSMUSG00000030759 | Far1 | -1.076 | 6.49E-05 | 3.67E-03 |
| ENSMUSG00000011114 | Tbrg1 | -1.075 | 2.49E-03 | 4.74E-02 |
| ENSMUSG00000037594 | BC022687 | -1.074 | 2.13E-03 | 4.24E-02 |
| ENSMUSG00000007739 | Cct4 | -1.071 | 1.19E-04 | 5.53E-03 |
| ENSMUSG00000081603 | Gm14681 | -1.071 | 6.55E-04 | 1.88E-02 |
| ENSMUSG00000032279 | Idh3a | -1.069 | 3.68E-06 | 4.18E-04 |
| ENSMUSG00000026356 | Dars | -1.069 | 1.66E-03 | 3.58E-02 |
| ENSMUSG00000001674 | Ddx18 | -1.061 | 5.65E-05 | 3.34E-03 |
| ENSMUSG00000022814 | Umps | -1.060 | 1.15E-05 | 9.65E-04 |
| ENSMUSG00000023883 | Phf10 | -1.060 | 3.45E-04 | 1.19E-02 |
| ENSMUSG00000034681 | Rnps1 | -1.059 | 1.11E-03 | 2.75E-02 |
| ENSMUSG00000030045 | Mrpl19 | -1.057 | 1.17E-03 | 2.82E-02 |
| ENSMUSG00000028889 | Yrdc | -1.053 | 1.10E-04 | 5.31E-03 |
| ENSMUSG00000025630 | Hprt | -1.052 | 1.41E-03 | 3.22E-02 |
| ENSMUSG00000031657 | Heatr3 | -1.047 | 1.49E-03 | 3.34E-02 |
| ENSMUSG00000001891 | Ugp2 | -1.047 | 1.04E-04 | 5.13E-03 |
| ENSMUSG00000002660 | Clpp | -1.040 | 2.35E-03 | 4.57E-02 |
| ENSMUSG00000021958 | Pinx1 | -1.040 | 3.24E-04 | 1.12E-02 |
| ENSMUSG00000025190 | Got1 | -1.039 | 7.23E-05 | 3.97E-03 |
| ENSMUSG00000017561 | Crlf3 | -1.034 | 1.00E-03 | 2.55E-02 |
| ENSMUSG00000071172 | Srsf3 | -1.032 | 1.75E-03 | 3.69E-02 |
| ENSMUSG00000003808 | Farsa | -1.027 | 1.32E-03 | 3.08E-02 |
| ENSMUSG00000022752 | Tomm70a | -1.027 | 7.51E-04 | 2.05E-02 |
| ENSMUSG00000090137 | Uba52 | -1.027 | 5.66E-06 | 5.79E-04 |
| ENSMUSG00000042719 | Naa25 | -1.022 | 1.23E-04 | 5.63E-03 |
| ENSMUSG00000018965 | Ywhah | -1.020 | 5.67E-04 | 1.69E-02 |
| ENSMUSG00000026926 | Pmpca | -1.017 | 3.11E-04 | 1.10E-02 |
| ENSMUSG00000022792 | Yars2 | -1.017 | 6.25E-04 | 1.80E-02 |
| ENSMUSG00000047649 | Cd3eap | -1.016 | 7.58E-04 | 2.06E-02 |
| ENSMUSG00000018697 | Aatf | -1.014 | 4.68E-04 | 1.45E-02 |
| ENSMUSG00000004393 | Ddx56 | -1.013 | 2.17E-03 | 4.29E-02 |
| ENSMUSG00000028333 | Anp32b | -1.013 | 8.30E-06 | 7.71E-04 |
| ENSMUSG00000042225 | Ammecr1 | -1.011 | 1.49E-04 | 6.43E-03 |
| ENSMUSG00000021276 | Cinp | -1.000 | 2.92E-04 | 1.05E-02 |
| ENSMUSG00000022663 | Atg3 | -0.998 | 4.58E-04 | 1.43E-02 |
| ENSMUSG00000039067 | Psmd7 | -0.996 | 2.52E-03 | 4.79E-02 |
| ENSMUSG00000030265 | Kras | -0.996 | 3.38E-05 | 2.24E-03 |
| ENSMUSG00000030521 | Mphosph10 | -0.992 | 2.51E-03 | 4.79E-02 |
| ENSMUSG00000032026 | Rexo2 | -0.989 | 1.62E-03 | 3.54E-02 |
| ENSMUSG00000019951 | Uhrf1bp1l | -0.989 | 2.32E-04 | 8.85E-03 |

| | | | | |
|---|---|---|---|---|
| ENSMUSG00000032557 | Uba5 | -0.982 | 1.68E-03 | 3.59E-02 |
| ENSMUSG00000038671 | Arfrp1 | -0.981 | 2.45E-03 | 4.70E-02 |
| ENSMUSG00000026036 | Nif3l1 | -0.975 | 1.37E-04 | 6.05E-03 |
| ENSMUSG00000057113 | Npm1 | -0.973 | 1.02E-04 | 5.07E-03 |
| ENSMUSG00000030942 | Thumpd1 | -0.970 | 1.92E-04 | 7.79E-03 |
| ENSMUSG00000030662 | Ipo5 | -0.969 | 2.90E-05 | 2.00E-03 |
| ENSMUSG00000111877 | Gm6477 | -0.958 | 1.75E-03 | 3.68E-02 |
| ENSMUSG00000028869 | Gnl2 | -0.958 | 1.05E-03 | 2.64E-02 |
| ENSMUSG00000021692 | Dimt1 | -0.956 | 2.02E-03 | 4.08E-02 |
| ENSMUSG00000022538 | Lsg1 | -0.948 | 2.53E-03 | 4.80E-02 |
| ENSMUSG00000021235 | Coq6 | -0.948 | 7.96E-04 | 2.14E-02 |
| ENSMUSG00000021737 | Psmd6 | -0.946 | 7.38E-07 | 1.30E-04 |
| ENSMUSG00000017999 | Ddx27 | -0.941 | 4.58E-04 | 1.43E-02 |
| ENSMUSG00000027357 | Crls1 | -0.941 | 2.66E-05 | 1.88E-03 |
| ENSMUSG00000024436 | Mrps18b | -0.939 | 1.19E-03 | 2.85E-02 |
| ENSMUSG00000020224 | Llph | -0.936 | 7.53E-04 | 2.06E-02 |
| ENSMUSG00000018293 | Pfn1 | -0.936 | 2.44E-03 | 4.68E-02 |
| ENSMUSG00000024712 | Rfk | -0.934 | 2.25E-04 | 8.70E-03 |
| ENSMUSG00000041028 | Ghitm | -0.929 | 1.06E-03 | 2.65E-02 |
| ENSMUSG00000043445 | Pgp | -0.927 | 7.08E-04 | 1.99E-02 |
| ENSMUSG00000031671 | Setd6 | -0.926 | 7.03E-04 | 1.98E-02 |
| ENSMUSG00000027671 | Actl6a | -0.920 | 7.23E-04 | 2.01E-02 |
| ENSMUSG00000061613 | U2af1 | -0.919 | 1.09E-03 | 2.70E-02 |
| ENSMUSG00000020464 | Pnpt1 | -0.918 | 1.63E-03 | 3.56E-02 |
| ENSMUSG00000025439 | Clns1a | -0.909 | 1.02E-03 | 2.59E-02 |
| ENSMUSG00000020088 | Sar1a | -0.908 | 1.31E-03 | 3.08E-02 |
| ENSMUSG00000026377 | Nifk | -0.907 | 1.10E-03 | 2.72E-02 |
| ENSMUSG00000020248 | Nfyb | -0.906 | 1.04E-04 | 5.13E-03 |
| ENSMUSG00000019782 | Rwdd1 | -0.904 | 8.41E-04 | 2.24E-02 |
| ENSMUSG00000036430 | Tbcc | -0.901 | 1.74E-03 | 3.68E-02 |
| ENSMUSG00000025393 | Atp5b | -0.897 | 1.38E-03 | 3.18E-02 |
| ENSMUSG00000022035 | Ccdc25 | -0.897 | 3.91E-04 | 1.30E-02 |
| ENSMUSG00000053907 | Mat2a | -0.896 | 1.04E-07 | 2.72E-05 |
| ENSMUSG00000028907 | Utp11 | -0.886 | 1.48E-04 | 6.43E-03 |
| ENSMUSG00000030224 | Strap | -0.880 | 2.06E-03 | 4.14E-02 |
| ENSMUSG00000020922 | Lsm12 | -0.878 | 2.75E-04 | 1.01E-02 |
| ENSMUSG00000040681 | Hmgn1 | -0.877 | 1.38E-03 | 3.18E-02 |
| ENSMUSG00000019795 | Pcmt1 | -0.876 | 3.82E-06 | 4.24E-04 |
| ENSMUSG00000045503 | Sys1 | -0.871 | 1.23E-04 | 5.62E-03 |
| ENSMUSG00000017428 | Psmd11 | -0.860 | 1.73E-03 | 3.67E-02 |
| ENSMUSG00000001774 | Chordc1 | -0.860 | 3.96E-05 | 2.52E-03 |
| ENSMUSG00000017801 | Mlx | -0.859 | 1.88E-03 | 3.87E-02 |
| ENSMUSG00000003438 | Timm50 | -0.856 | 1.24E-03 | 2.95E-02 |
| ENSMUSG00000020720 | Psmd12 | -0.853 | 7.71E-04 | 2.09E-02 |
| ENSMUSG00000015120 | Ube2i | -0.845 | 1.42E-05 | 1.13E-03 |
| ENSMUSG00000061207 | Stk19 | -0.844 | 1.85E-03 | 3.83E-02 |
| ENSMUSG00000107470 | Gm3375 | -0.841 | 1.02E-03 | 2.58E-02 |

| | | | | |
|---|---|---|---|---|
| ENSMUSG00000026869 | Psmd5 | -0.840 | 4.42E-04 | 1.40E-02 |
| ENSMUSG00000029551 | Psmg3 | -0.840 | 1.31E-04 | 5.82E-03 |
| ENSMUSG00000028187 | Rpf1 | -0.838 | 1.15E-03 | 2.80E-02 |
| ENSMUSG00000036371 | Serbp1 | -0.834 | 1.13E-04 | 5.40E-03 |
| ENSMUSG00000030879 | Mrpl17 | -0.832 | 1.78E-04 | 7.39E-03 |
| ENSMUSG00000028322 | Exosc3 | -0.831 | 4.27E-04 | 1.37E-02 |
| ENSMUSG00000040521 | Tsfm | -0.831 | 1.55E-03 | 3.44E-02 |
| ENSMUSG00000039886 | Tmem120a | -0.830 | 3.33E-04 | 1.15E-02 |
| ENSMUSG00000010554 | Mettl16 | -0.828 | 1.03E-03 | 2.60E-02 |
| ENSMUSG00000004268 | Emg1 | -0.826 | 3.08E-04 | 1.09E-02 |
| ENSMUSG00000030603 | Psmc4 | -0.821 | 1.20E-03 | 2.88E-02 |
| ENSMUSG00000038482 | Tfdp1 | -0.818 | 9.74E-04 | 2.50E-02 |
| ENSMUSG00000026159 | Agfg1 | -0.816 | 8.71E-04 | 2.30E-02 |
| ENSMUSG00000037740 | Mrps26 | -0.816 | 1.97E-04 | 7.95E-03 |
| ENSMUSG00000021660 | Btf3 | -0.815 | 1.69E-03 | 3.61E-02 |
| ENSMUSG00000018565 | Elp5 | -0.808 | 4.55E-06 | 4.86E-04 |
| ENSMUSG00000027601 | Mtfr1 | -0.806 | 2.84E-04 | 1.03E-02 |
| ENSMUSG00000021131 | Erh | -0.797 | 1.04E-03 | 2.62E-02 |
| ENSMUSG00000029922 | Mkrn1 | -0.795 | 3.19E-04 | 1.11E-02 |
| ENSMUSG00000020180 | Snrpd3 | -0.793 | 2.11E-04 | 8.25E-03 |
| ENSMUSG00000030512 | Snrpa1 | -0.788 | 2.32E-03 | 4.51E-02 |
| ENSMUSG00000024580 | Grpel2 | -0.785 | 4.27E-05 | 2.68E-03 |
| ENSMUSG00000028988 | Ctnnbip1 | -0.783 | 1.17E-03 | 2.83E-02 |
| ENSMUSG00000020775 | Mrpl38 | -0.780 | 2.04E-04 | 8.11E-03 |
| ENSMUSG00000051518 | Rps19bp1 | -0.779 | 1.36E-03 | 3.16E-02 |
| ENSMUSG00000035021 | Baz1a | -0.779 | 7.39E-04 | 2.04E-02 |
| ENSMUSG00000030880 | Polr3e | -0.777 | 1.35E-03 | 3.14E-02 |
| ENSMUSG00000014195 | Dnajc7 | -0.760 | 9.00E-04 | 2.36E-02 |
| ENSMUSG00000070697 | Utp3 | -0.756 | 9.93E-04 | 2.53E-02 |
| ENSMUSG00000023147 | Wrb | -0.755 | 2.29E-03 | 4.47E-02 |
| ENSMUSG00000021578 | Ccdc127 | -0.754 | 1.44E-03 | 3.27E-02 |
| ENSMUSG00000032002 | Dcun1d5 | -0.751 | 8.52E-05 | 4.52E-03 |
| ENSMUSG00000078652 | Psme3 | -0.738 | 2.11E-03 | 4.21E-02 |
| ENSMUSG00000022204 | Ngdn | -0.737 | 1.75E-05 | 1.36E-03 |
| ENSMUSG00000040843 | Tiprl | -0.736 | 1.68E-03 | 3.59E-02 |
| ENSMUSG00000034620 | Tmem5 | -0.720 | 1.90E-03 | 3.89E-02 |
| ENSMUSG00000032563 | Mrpl3 | -0.691 | 2.45E-04 | 9.29E-03 |
| ENSMUSG00000079478 | Sssca1 | -0.684 | 2.01E-03 | 4.06E-02 |
| ENSMUSG00000024878 | Cbwd1 | -0.682 | 1.25E-03 | 2.96E-02 |
| ENSMUSG00000015290 | Ubl4a | -0.677 | 1.64E-03 | 3.57E-02 |
| ENSMUSG00000021018 | Polr2h | -0.674 | 2.13E-03 | 4.24E-02 |
| ENSMUSG00000039640 | Mrpl12 | -0.674 | 2.72E-04 | 1.00E-02 |
| ENSMUSG00000055762 | Eef1d | -0.667 | 1.90E-03 | 3.89E-02 |
| ENSMUSG00000029038 | Ssu72 | -0.657 | 4.52E-04 | 1.42E-02 |
| ENSMUSG00000022858 | Tra2b | -0.654 | 2.22E-03 | 4.38E-02 |
| ENSMUSG00000001416 | Cct3 | -0.651 | 2.02E-03 | 4.08E-02 |
| ENSMUSG00000031697 | Orc6 | -0.623 | 2.10E-03 | 4.20E-02 |

| | | | | |
|---|---|---|---|---|
| ENSMUSG00000043284 | Tmem11 | -0.622 | 7.94E-04 | 2.14E-02 |
| ENSMUSG00000061360 | Phf5a | -0.609 | 4.81E-04 | 1.49E-02 |
| ENSMUSG00000034024 | Cct2 | -0.606 | 9.05E-05 | 4.65E-03 |
| ENSMUSG00000043866 | Taf10 | -0.603 | 1.28E-03 | 3.01E-02 |
| ENSMUSG00000022427 | Tomm22 | -0.371 | 2.60E-03 | 4.90E-02 |
| ENSMUSG00000022671 | Mzt2 | 0.577 | 2.13E-03 | 4.24E-02 |
| ENSMUSG00000025102 | 3110040N11 | 0.603 | 1.46E-03 | 3.31E-02 |
| ENSMUSG00000002345 | Borcs8 | 0.644 | 2.28E-03 | 4.47E-02 |
| ENSMUSG00000027104 | Atf2 | 0.708 | 9.52E-05 | 4.82E-03 |
| ENSMUSG00000031360 | Ctps2 | 0.729 | 6.15E-05 | 3.57E-03 |
| ENSMUSG00000031023 | Akip1 | 0.736 | 1.24E-04 | 5.63E-03 |
| ENSMUSG00000037608 | Bclaf1 | 0.742 | 6.06E-04 | 1.77E-02 |
| ENSMUSG00000025040 | Fundc1 | 0.746 | 2.58E-03 | 4.87E-02 |
| ENSMUSG00000029433 | Diablo | 0.747 | 6.74E-05 | 3.78E-03 |
| ENSMUSG00000009030 | Pdcl | 0.763 | 9.64E-05 | 4.84E-03 |
| ENSMUSG00000022051 | Bnip3l | 0.767 | 2.60E-03 | 4.89E-02 |
| ENSMUSG00000051285 | Pcmtd1 | 0.799 | 1.92E-03 | 3.91E-02 |
| ENSMUSG00000066724 | Gm10175 | 0.802 | 1.17E-03 | 2.83E-02 |
| ENSMUSG00000028070 | Naxe | 0.805 | 1.15E-03 | 2.80E-02 |
| ENSMUSG00000052459 | Atp6v1a | 0.810 | 2.52E-03 | 4.79E-02 |
| ENSMUSG00000056211 | R3hdm1 | 0.828 | 5.28E-04 | 1.60E-02 |
| ENSMUSG00000058325 | Dock1 | 0.842 | 8.01E-04 | 2.15E-02 |
| ENSMUSG00000025505 | Tmem80 | 0.851 | 2.35E-03 | 4.57E-02 |
| ENSMUSG00000039159 | Ube2h | 0.857 | 1.08E-03 | 2.70E-02 |
| ENSMUSG00000001018 | Snapin | 0.861 | 1.05E-03 | 2.64E-02 |
| ENSMUSG00000025035 | Arl3 | 0.862 | 6.94E-04 | 1.96E-02 |
| ENSMUSG00000027709 | Mccc1 | 0.864 | 1.79E-03 | 3.74E-02 |
| ENSMUSG00000028062 | Lamtor2 | 0.866 | 1.25E-03 | 2.96E-02 |
| ENSMUSG00000053929 | Cyhr1 | 0.879 | 3.27E-05 | 2.18E-03 |
| ENSMUSG00000030613 | Ccdc90b | 0.879 | 3.14E-04 | 1.11E-02 |
| ENSMUSG00000034951 | Cog7 | 0.888 | 7.97E-04 | 2.14E-02 |
| ENSMUSG00000033596 | Rfwd3 | 0.900 | 1.39E-03 | 3.19E-02 |
| ENSMUSG00000024038 | Ndufv3 | 0.905 | 6.32E-04 | 1.82E-02 |
| ENSMUSG00000022092 | Ppp3cc | 0.912 | 1.58E-03 | 3.49E-02 |
| ENSMUSG00000050552 | Lamtor4 | 0.924 | 9.28E-04 | 2.42E-02 |
| ENSMUSG00000032398 | Snapc5 | 0.932 | 1.25E-05 | 1.02E-03 |
| ENSMUSG00000021109 | Hif1a | 0.935 | 1.09E-04 | 5.28E-03 |
| ENSMUSG00000034744 | Nagk | 0.941 | 1.92E-03 | 3.91E-02 |
| ENSMUSG00000020091 | Eif4ebp2 | 0.944 | 3.15E-04 | 1.11E-02 |
| ENSMUSG00000030515 | Tarsl2 | 0.962 | 7.93E-04 | 2.14E-02 |
| ENSMUSG00000021948 | Prkcd | 0.967 | 1.64E-03 | 3.58E-02 |
| ENSMUSG00000030213 | Atf7ip | 0.970 | 3.52E-04 | 1.21E-02 |
| ENSMUSG00000079427 | Mthfsl | 0.982 | 2.22E-04 | 8.62E-03 |
| ENSMUSG00000039795 | Zfand1 | 0.987 | 1.68E-03 | 3.59E-02 |
| ENSMUSG00000028580 | Pum1 | 0.987 | 2.36E-03 | 4.58E-02 |
| ENSMUSG00000036880 | Acaa2 | 0.999 | 2.59E-03 | 4.89E-02 |
| ENSMUSG00000044201 | Cdc25c | 1.002 | 1.66E-03 | 3.58E-02 |

| | | | | |
|---|---|---|---|---|
| ENSMUSG00000022617 | Chkb | 1.002 | 6.75E-04 | 1.92E-02 |
| ENSMUSG00000040128 | Pnrc1 | 1.011 | 6.86E-04 | 1.94E-02 |
| ENSMUSG00000078566 | Bnip3 | 1.033 | 7.59E-04 | 2.06E-02 |
| ENSMUSG00000025860 | Xiap | 1.036 | 1.63E-03 | 3.56E-02 |
| ENSMUSG00000035637 | Grhpr | 1.039 | 6.09E-04 | 1.77E-02 |
| ENSMUSG00000057367 | Birc2 | 1.048 | 5.00E-04 | 1.53E-02 |
| ENSMUSG00000073198 | Bnip3l-ps | 1.050 | 1.52E-03 | 3.38E-02 |
| ENSMUSG00000057572 | Zbtb8os | 1.055 | 1.40E-03 | 3.22E-02 |
| ENSMUSG00000023966 | Rsph9 | 1.058 | 1.82E-04 | 7.52E-03 |
| ENSMUSG00000029401 | Rilpl2 | 1.065 | 4.18E-04 | 1.35E-02 |
| ENSMUSG00000006021 | Kptn | 1.067 | 8.28E-04 | 2.21E-02 |
| ENSMUSG00000040795 | Iqcc | 1.069 | 2.02E-03 | 4.08E-02 |
| ENSMUSG00000035901 | Dennd5a | 1.078 | 4.44E-04 | 1.40E-02 |
| ENSMUSG00000042364 | Snx18 | 1.084 | 1.17E-04 | 5.48E-03 |
| ENSMUSG00000026721 | Rabgap1l | 1.090 | 1.15E-03 | 2.80E-02 |
| ENSMUSG00000002580 | Mien1 | 1.115 | 3.85E-05 | 2.46E-03 |
| ENSMUSG00000033111 | 3830406C13 | 1.115 | 6.80E-06 | 6.61E-04 |
| ENSMUSG00000036644 | Tbc1d9b | 1.116 | 2.38E-03 | 4.62E-02 |
| ENSMUSG00000031641 | Cbr4 | 1.120 | 3.73E-05 | 2.42E-03 |
| ENSMUSG00000042408 | Zmym6 | 1.124 | 9.56E-04 | 2.47E-02 |
| ENSMUSG00000028382 | Ptbp3 | 1.125 | 3.56E-04 | 1.21E-02 |
| ENSMUSG00000040549 | Ckap5 | 1.139 | 4.58E-04 | 1.43E-02 |
| ENSMUSG00000021643 | Serf1 | 1.148 | 2.30E-03 | 4.48E-02 |
| ENSMUSG00000054894 | Atp5s | 1.148 | 1.60E-04 | 6.85E-03 |
| ENSMUSG00000000486 | 45170 | 1.149 | 1.46E-03 | 3.31E-02 |
| ENSMUSG00000042055 | Wdr11 | 1.164 | 1.99E-03 | 4.04E-02 |
| ENSMUSG00000037847 | Nmrk1 | 1.166 | 2.17E-03 | 4.29E-02 |
| ENSMUSG00000031529 | Tnks | 1.167 | 2.52E-03 | 4.79E-02 |
| ENSMUSG00000021033 | Gstz1 | 1.181 | 6.22E-06 | 6.19E-04 |
| ENSMUSG00000029471 | Camkk2 | 1.191 | 9.81E-04 | 2.51E-02 |
| ENSMUSG00000026154 | Sdhaf4 | 1.194 | 1.69E-03 | 3.61E-02 |
| ENSMUSG00000037152 | Ndufc1 | 1.196 | 1.28E-03 | 3.01E-02 |
| ENSMUSG00000032038 | St3gal4 | 1.198 | 1.74E-04 | 7.31E-03 |
| ENSMUSG00000074749 | Kiz | 1.203 | 2.31E-04 | 8.82E-03 |
| ENSMUSG00000035142 | Nubpl | 1.204 | 6.17E-04 | 1.79E-02 |
| ENSMUSG00000026385 | Dbi | 1.206 | 2.45E-03 | 4.70E-02 |
| ENSMUSG00000027984 | Hadh | 1.207 | 3.16E-05 | 2.12E-03 |
| ENSMUSG00000037628 | Cdkn3 | 1.219 | 2.63E-03 | 4.95E-02 |
| ENSMUSG00000039745 | Htatip2 | 1.225 | 1.06E-03 | 2.66E-02 |
| ENSMUSG00000028101 | Pias3 | 1.231 | 1.25E-03 | 2.97E-02 |
| ENSMUSG00000060147 | Serpinb6a | 1.234 | 1.49E-04 | 6.43E-03 |
| ENSMUSG00000006717 | Acot13 | 1.240 | 3.35E-06 | 3.97E-04 |
| ENSMUSG00000024392 | Bag6 | 1.242 | 6.00E-04 | 1.76E-02 |
| ENSMUSG00000042506 | Usp22 | 1.244 | 7.80E-04 | 2.11E-02 |
| ENSMUSG00000022817 | Itgb5 | 1.245 | 3.16E-04 | 1.11E-02 |
| ENSMUSG00000042675 | Ypel3 | 1.249 | 1.90E-05 | 1.43E-03 |
| ENSMUSG00000038014 | Fam120a | 1.257 | 1.06E-04 | 5.21E-03 |

| | | | | |
|---|---|---|---|---|
| ENSMUSG00000025993 | Slc40a1 | 1.261 | 8.67E-05 | 4.58E-03 |
| ENSMUSG00000031161 | Hdac6 | 1.262 | 1.67E-03 | 3.59E-02 |
| ENSMUSG00000065979 | Cpped1 | 1.264 | 5.80E-04 | 1.72E-02 |
| ENSMUSG00000037499 | Nenf | 1.265 | 1.47E-03 | 3.31E-02 |
| ENSMUSG00000040841 | Six5 | 1.267 | 2.23E-05 | 1.62E-03 |
| ENSMUSG00000030779 | Rbbp6 | 1.272 | 9.71E-04 | 2.50E-02 |
| ENSMUSG00000042155 | Klhl23 | 1.274 | 3.99E-04 | 1.31E-02 |
| ENSMUSG00000014075 | Tctex1d2 | 1.278 | 5.80E-05 | 3.42E-03 |
| ENSMUSG00000027165 | B230118H07 | 1.287 | 1.32E-04 | 5.87E-03 |
| ENSMUSG00000008822 | Acyp1 | 1.287 | 4.16E-04 | 1.35E-02 |
| ENSMUSG00000042293 | Gm5617 | 1.288 | 2.55E-04 | 9.57E-03 |
| ENSMUSG00000025731 | Mettl26 | 1.289 | 2.39E-03 | 4.63E-02 |
| ENSMUSG00000023911 | Flywch2 | 1.294 | 1.91E-03 | 3.90E-02 |
| ENSMUSG00000028549 | Itgb3bp | 1.296 | 4.73E-04 | 1.46E-02 |
| ENSMUSG00000040370 | Etfrf1 | 1.297 | 3.53E-04 | 1.21E-02 |
| ENSMUSG00000038718 | Pbx3 | 1.299 | 2.69E-04 | 9.95E-03 |
| ENSMUSG00000022773 | Ypel1 | 1.299 | 1.88E-03 | 3.87E-02 |
| ENSMUSG00000008035 | Mid1ip1 | 1.303 | 1.30E-03 | 3.06E-02 |
| ENSMUSG00000028470 | Hint2 | 1.303 | 1.24E-04 | 5.63E-03 |
| ENSMUSG00000070394 | Tmem256 | 1.310 | 1.33E-03 | 3.10E-02 |
| ENSMUSG00000033429 | Mcee | 1.311 | 4.54E-05 | 2.80E-03 |
| ENSMUSG00000023094 | Msrb2 | 1.313 | 5.16E-04 | 1.57E-02 |
| ENSMUSG00000041126 | H2afv | 1.314 | 9.22E-06 | 8.27E-04 |
| ENSMUSG00000078695 | Cisd3 | 1.320 | 4.67E-04 | 1.45E-02 |
| ENSMUSG00000072772 | Grcc10 | 1.320 | 3.71E-05 | 2.42E-03 |
| ENSMUSG00000024960 | Plcb3 | 1.321 | 1.57E-04 | 6.75E-03 |
| ENSMUSG00000057103 | Nat8f1 | 1.323 | 7.33E-04 | 2.03E-02 |
| ENSMUSG00000044475 | Ascc1 | 1.328 | 1.98E-04 | 7.95E-03 |
| ENSMUSG00000022708 | Zbtb20 | 1.345 | 2.77E-04 | 1.01E-02 |
| ENSMUSG00000052102 | Gnpda1 | 1.347 | 8.91E-05 | 4.64E-03 |
| ENSMUSG00000097867 | Lppos | 1.352 | 1.46E-03 | 3.31E-02 |
| ENSMUSG00000073633 | Fbxo36 | 1.366 | 4.42E-04 | 1.40E-02 |
| ENSMUSG00000032216 | Nedd4 | 1.366 | 7.46E-05 | 4.08E-03 |
| ENSMUSG00000028756 | Pink1 | 1.367 | 2.03E-03 | 4.09E-02 |
| ENSMUSG00000032702 | Kank1 | 1.373 | 1.16E-03 | 2.82E-02 |
| ENSMUSG00000029521 | Chek2 | 1.381 | 1.94E-03 | 3.94E-02 |
| ENSMUSG00000036782 | Klhl13 | 1.385 | 1.58E-03 | 3.49E-02 |
| ENSMUSG00000045867 | Cradd | 1.391 | 1.33E-04 | 5.91E-03 |
| ENSMUSG00000027654 | Fam83d | 1.391 | 6.53E-04 | 1.87E-02 |
| ENSMUSG00000022550 | Adck5 | 1.394 | 6.07E-04 | 1.77E-02 |
| ENSMUSG00000027187 | Cat | 1.403 | 7.86E-05 | 4.21E-03 |
| ENSMUSG00000052139 | Babam2 | 1.407 | 9.80E-06 | 8.61E-04 |
| ENSMUSG00000028795 | Ccdc28b | 1.408 | 3.21E-08 | 1.11E-05 |
| ENSMUSG00000020150 | Gamt | 1.411 | 8.04E-04 | 2.16E-02 |
| ENSMUSG00000002058 | Unc119 | 1.419 | 3.73E-05 | 2.42E-03 |
| ENSMUSG00000049225 | Pdp1 | 1.419 | 1.26E-08 | 5.17E-06 |
| ENSMUSG00000002043 | Trappc6a | 1.420 | 1.18E-04 | 5.50E-03 |

| | | | | |
|---|---|---|---|---|
| ENSMUSG00000097059 | Fam120aos | 1.427 | 1.26E-03 | 2.98E-02 |
| ENSMUSG00000001666 | Ddt | 1.428 | 1.02E-03 | 2.59E-02 |
| ENSMUSG00000030990 | Pgap2 | 1.431 | 3.53E-06 | 4.11E-04 |
| ENSMUSG00000046982 | Tshz1 | 1.434 | 1.56E-03 | 3.45E-02 |
| ENSMUSG00000035642 | Aamdc | 1.436 | 6.22E-05 | 3.57E-03 |
| ENSMUSG00000019689 | Fmc1 | 1.438 | 3.07E-05 | 2.07E-03 |
| ENSMUSG00000020898 | Ctc1 | 1.443 | 4.13E-04 | 1.35E-02 |
| ENSMUSG00000034610 | Zcchc11 | 1.454 | 2.00E-04 | 8.04E-03 |
| ENSMUSG00000033685 | Ucp2 | 1.459 | 3.36E-05 | 2.23E-03 |
| ENSMUSG00000044252 | Osbpl1a | 1.461 | 2.53E-06 | 3.39E-04 |
| ENSMUSG00000021792 | Fam213a | 1.465 | 1.47E-03 | 3.31E-02 |
| ENSMUSG00000029385 | Ccng2 | 1.468 | 6.29E-06 | 6.22E-04 |
| ENSMUSG00000064215 | Ifi27 | 1.485 | 9.16E-06 | 8.27E-04 |
| ENSMUSG00000033065 | Pfkm | 1.506 | 1.43E-03 | 3.26E-02 |
| ENSMUSG00000027332 | Ivd | 1.509 | 1.12E-05 | 9.42E-04 |
| ENSMUSG00000037253 | Mex3c | 1.516 | 2.64E-04 | 9.88E-03 |
| ENSMUSG00000010362 | Rdm1 | 1.521 | 2.57E-05 | 1.84E-03 |
| ENSMUSG00000060261 | Gtf2i | 1.537 | 1.91E-04 | 7.79E-03 |
| ENSMUSG00000039286 | Fndc3b | 1.556 | 1.60E-03 | 3.52E-02 |
| ENSMUSG00000032220 | Myo1e | 1.565 | 8.97E-04 | 2.36E-02 |
| ENSMUSG00000035048 | Anapc13 | 1.567 | 7.84E-05 | 4.21E-03 |
| ENSMUSG00000058756 | Thra | 1.567 | 9.65E-04 | 2.48E-02 |
| ENSMUSG00000020601 | Trib2 | 1.572 | 8.02E-06 | 7.56E-04 |
| ENSMUSG00000050538 | B230217C12 | 1.573 | 2.06E-03 | 4.13E-02 |
| ENSMUSG00000045257 | Morn2 | 1.573 | 2.36E-04 | 9.01E-03 |
| ENSMUSG00000037493 | Cib2 | 1.579 | 4.64E-04 | 1.45E-02 |
| ENSMUSG00000034449 | Dhrs11 | 1.583 | 1.78E-04 | 7.39E-03 |
| ENSMUSG00000022040 | Ephx2 | 1.593 | 1.53E-03 | 3.40E-02 |
| ENSMUSG00000025955 | Akr1cl | 1.593 | 6.20E-05 | 3.57E-03 |
| ENSMUSG00000023055 | Calcoco1 | 1.595 | 5.55E-05 | 3.30E-03 |
| ENSMUSG00000023150 | Ivns1abp | 1.601 | 6.07E-04 | 1.77E-02 |
| ENSMUSG00000024866 | Acy3 | 1.610 | 2.09E-04 | 8.20E-03 |
| ENSMUSG00000036879 | Phkb | 1.610 | 6.50E-04 | 1.87E-02 |
| ENSMUSG00000066456 | Hmgn3 | 1.611 | 1.29E-05 | 1.05E-03 |
| ENSMUSG00000021065 | Fut8 | 1.611 | 6.86E-04 | 1.94E-02 |
| ENSMUSG00000048490 | Nrip1 | 1.623 | 1.87E-03 | 3.86E-02 |
| ENSMUSG00000097415 | AU020206 | 1.624 | 1.61E-04 | 6.85E-03 |
| ENSMUSG00000022048 | Dpysl2 | 1.633 | 7.81E-04 | 2.11E-02 |
| ENSMUSG00000030122 | Ptms | 1.633 | 2.26E-04 | 8.70E-03 |
| ENSMUSG00000038286 | Bphl | 1.639 | 3.14E-06 | 3.94E-04 |
| ENSMUSG00000031666 | Rbl2 | 1.641 | 4.03E-04 | 1.32E-02 |
| ENSMUSG00000038070 | Cntln | 1.648 | 5.49E-04 | 1.64E-02 |
| ENSMUSG00000018417 | Myo1b | 1.656 | 3.84E-04 | 1.28E-02 |
| ENSMUSG00000034522 | Zfp395 | 1.667 | 4.62E-06 | 4.88E-04 |
| ENSMUSG00000021775 | Nr1d2 | 1.670 | 1.34E-06 | 2.08E-04 |
| ENSMUSG00000022587 | Ly6e | 1.680 | 3.80E-04 | 1.27E-02 |
| ENSMUSG00000039450 | Dcxr | 1.697 | 1.05E-06 | 1.70E-04 |

| | | | | |
|---|---|---|---|---|
| ENSMUSG00000032067 | Pts | 1.698 | 2.79E-08 | 9.81E-06 |
| ENSMUSG00000022323 | Rida | 1.699 | 4.45E-05 | 2.77E-03 |
| ENSMUSG00000046718 | Bst2 | 1.703 | 1.77E-03 | 3.72E-02 |
| ENSMUSG00000027589 | Pcmtd2 | 1.710 | 4.18E-06 | 4.56E-04 |
| ENSMUSG00000100975 | Gm28875 | 1.723 | 1.00E-03 | 2.55E-02 |
| ENSMUSG00000086158 | Ccpg1os | 1.730 | 4.38E-05 | 2.74E-03 |
| ENSMUSG00000026211 | Obsl1 | 1.737 | 1.76E-06 | 2.55E-04 |
| ENSMUSG00000026495 | Efcab2 | 1.737 | 9.22E-05 | 4.71E-03 |
| ENSMUSG00000087165 | 2010001A14 | 1.739 | 8.55E-04 | 2.27E-02 |
| ENSMUSG00000026839 | Upp2 | 1.746 | 2.68E-04 | 9.93E-03 |
| ENSMUSG00000041390 | Mdfic | 1.749 | 2.30E-03 | 4.48E-02 |
| ENSMUSG00000047003 | Zfp41 | 1.753 | 2.16E-04 | 8.42E-03 |
| ENSMUSG00000073755 | 5730409E04 | 1.753 | 1.13E-03 | 2.77E-02 |
| ENSMUSG00000025404 | R3hdm2 | 1.760 | 1.69E-05 | 1.33E-03 |
| ENSMUSG00000051817 | Sox12 | 1.766 | 2.07E-06 | 2.88E-04 |
| ENSMUSG00000032609 | Klhdc8b | 1.781 | 1.83E-03 | 3.80E-02 |
| ENSMUSG00000097162 | 2310010J17I | 1.789 | 1.21E-04 | 5.58E-03 |
| ENSMUSG00000032221 | Mns1 | 1.800 | 1.66E-03 | 3.58E-02 |
| ENSMUSG00000028108 | Ecm1 | 1.800 | 1.26E-04 | 5.66E-03 |
| ENSMUSG00000054723 | Vmac | 1.804 | 2.57E-05 | 1.84E-03 |
| ENSMUSG00000086468 | Etaa1os | 1.811 | 1.16E-04 | 5.48E-03 |
| ENSMUSG00000025466 | Fuom | 1.816 | 5.48E-06 | 5.66E-04 |
| ENSMUSG00000046329 | Slc25a23 | 1.822 | 3.80E-04 | 1.27E-02 |
| ENSMUSG00000043644 | 0610009L18 | 1.824 | 3.17E-04 | 1.11E-02 |
| ENSMUSG00000040249 | Lrp1 | 1.826 | 7.22E-04 | 2.01E-02 |
| ENSMUSG00000051343 | Rab11fip5 | 1.828 | 2.88E-04 | 1.04E-02 |
| ENSMUSG00000010307 | Tmem86a | 1.837 | 5.82E-04 | 1.72E-02 |
| ENSMUSG00000079598 | Clec2l | 1.838 | 1.58E-03 | 3.49E-02 |
| ENSMUSG00000021238 | Aldh6a1 | 1.844 | 5.19E-04 | 1.57E-02 |
| ENSMUSG00000026664 | Phyh | 1.844 | 6.39E-08 | 1.82E-05 |
| ENSMUSG00000107369 | Gstm2-ps1 | 1.853 | 1.22E-05 | 1.00E-03 |
| ENSMUSG00000091561 | Gm6665 | 1.854 | 9.23E-05 | 4.71E-03 |
| ENSMUSG00000024253 | Dync2li1 | 1.861 | 1.65E-09 | 9.65E-07 |
| ENSMUSG00000034101 | Ctnnd1 | 1.863 | 1.50E-03 | 3.36E-02 |
| ENSMUSG00000087336 | Gm15860 | 1.883 | 9.75E-04 | 2.50E-02 |
| ENSMUSG00000039958 | Etfbkmt | 1.889 | 4.21E-04 | 1.36E-02 |
| ENSMUSG00000048581 | E130311K13 | 1.904 | 4.25E-06 | 4.61E-04 |
| ENSMUSG00000021690 | Jmy | 1.914 | 3.85E-05 | 2.46E-03 |
| ENSMUSG00000022661 | Cd200 | 1.927 | 1.44E-05 | 1.15E-03 |
| ENSMUSG00000031453 | Rasa3 | 1.935 | 6.48E-06 | 6.33E-04 |
| ENSMUSG00000004633 | Chn2 | 1.936 | 1.87E-03 | 3.86E-02 |
| ENSMUSG00000086587 | Gm11837 | 1.945 | 1.08E-04 | 5.26E-03 |
| ENSMUSG00000000594 | Gm2a | 1.948 | 2.48E-03 | 4.73E-02 |
| ENSMUSG00000033577 | Myo6 | 1.954 | 1.49E-05 | 1.18E-03 |
| ENSMUSG00000044080 | S100a1 | 1.962 | 1.62E-06 | 2.44E-04 |
| ENSMUSG00000058135 | Gstm1 | 1.966 | 2.99E-06 | 3.83E-04 |
| ENSMUSG00000040562 | Gstm2 | 1.974 | 4.74E-05 | 2.88E-03 |

| | | | | |
|---|---|---|---|---|
| ENSMUSG00000053279 | Aldh1a1 | 1.977 | 2.64E-05 | 1.88E-03 |
| ENSMUSG00000044986 | Tst | 1.982 | 1.25E-04 | 5.63E-03 |
| ENSMUSG00000032009 | Sesn3 | 1.988 | 6.02E-06 | 6.06E-04 |
| ENSMUSG00000060675 | Pla2g16 | 1.989 | 4.64E-05 | 2.84E-03 |
| ENSMUSG00000036894 | Rap2b | 2.000 | 1.50E-03 | 3.36E-02 |
| ENSMUSG00000067336 | Bmpr2 | 2.014 | 4.57E-04 | 1.43E-02 |
| ENSMUSG00000037606 | Osbpl5 | 2.049 | 4.16E-04 | 1.35E-02 |
| ENSMUSG00000003031 | Cdkn1b | 2.050 | 1.18E-05 | 9.74E-04 |
| ENSMUSG00000024646 | Cyb5a | 2.050 | 6.19E-05 | 3.57E-03 |
| ENSMUSG00000049086 | Bmyc | 2.057 | 2.88E-04 | 1.04E-02 |
| ENSMUSG00000062866 | Phactr2 | 2.058 | 1.85E-07 | 4.07E-05 |
| ENSMUSG00000027935 | Rab13 | 2.059 | 1.33E-05 | 1.07E-03 |
| ENSMUSG00000042712 | Tceal9 | 2.061 | 7.20E-07 | 1.28E-04 |
| ENSMUSG00000020216 | Jsrp1 | 2.063 | 2.68E-04 | 9.93E-03 |
| ENSMUSG00000043079 | Synpo | 2.068 | 1.83E-03 | 3.80E-02 |
| ENSMUSG00000021619 | Atg10 | 2.082 | 6.65E-05 | 3.75E-03 |
| ENSMUSG00000022390 | Zc3h7b | 2.100 | 4.52E-05 | 2.79E-03 |
| ENSMUSG00000019718 | L3hypdh | 2.111 | 4.27E-06 | 4.61E-04 |
| ENSMUSG00000023191 | P3h3 | 2.114 | 5.89E-04 | 1.74E-02 |
| ENSMUSG00000013033 | Adgrl1 | 2.115 | 7.45E-04 | 2.05E-02 |
| ENSMUSG00000036053 | Fmnl2 | 2.115 | 1.46E-03 | 3.31E-02 |
| ENSMUSG00000021094 | Dhrs7 | 2.134 | 2.40E-03 | 4.65E-02 |
| ENSMUSG00000098318 | Lockd | 2.140 | 1.04E-04 | 5.13E-03 |
| ENSMUSG00000031099 | Smarca1 | 2.141 | 2.61E-07 | 5.49E-05 |
| ENSMUSG00000026365 | Cfh | 2.163 | 2.22E-03 | 4.38E-02 |
| ENSMUSG00000011382 | Dhdh | 2.168 | 1.13E-03 | 2.77E-02 |
| ENSMUSG00000111080 | Gm20300 | 2.173 | 4.79E-05 | 2.90E-03 |
| ENSMUSG00000073985 | Gm10602 | 2.177 | 3.85E-04 | 1.28E-02 |
| ENSMUSG00000030630 | Fah | 2.193 | 2.70E-05 | 1.90E-03 |
| ENSMUSG00000066357 | Wdr6 | 2.195 | 2.75E-10 | 1.97E-07 |
| ENSMUSG00000067878 | Map7d3 | 2.196 | 8.96E-04 | 2.36E-02 |
| ENSMUSG00000031760 | Mt3 | 2.205 | 6.72E-04 | 1.92E-02 |
| ENSMUSG00000089768 | Tmsb15b1 | 2.209 | 8.99E-04 | 2.36E-02 |
| ENSMUSG00000027223 | Mapk8ip1 | 2.216 | 2.42E-03 | 4.66E-02 |
| ENSMUSG00000059645 | Gm7361 | 2.220 | 3.61E-04 | 1.22E-02 |
| ENSMUSG00000112876 | Gm32443 | 2.234 | 2.03E-04 | 8.10E-03 |
| ENSMUSG00000031613 | Hpgd | 2.246 | 5.75E-04 | 1.71E-02 |
| ENSMUSG00000030329 | Pianp | 2.247 | 2.43E-03 | 4.67E-02 |
| ENSMUSG00000040557 | Mettl27 | 2.267 | 2.41E-10 | 1.79E-07 |
| ENSMUSG00000037826 | Ppm1k | 2.273 | 1.32E-03 | 3.08E-02 |
| ENSMUSG00000033910 | Gucy1a1 | 2.307 | 8.74E-04 | 2.30E-02 |
| ENSMUSG00000024713 | Pcsk5 | 2.308 | 2.52E-04 | 9.48E-03 |
| ENSMUSG00000026817 | Ak1 | 2.310 | 1.83E-03 | 3.80E-02 |
| ENSMUSG00000002409 | Dyrk1b | 2.318 | 1.49E-03 | 3.34E-02 |
| ENSMUSG00000030055 | Rab43 | 2.322 | 8.66E-04 | 2.29E-02 |
| ENSMUSG00000096727 | Psmb9 | 2.323 | 1.13E-03 | 2.77E-02 |
| ENSMUSG00000027848 | Olfml3 | 2.327 | 5.39E-04 | 1.62E-02 |

| | | | | |
|---|---|---|---|---|
| ENSMUSG00000027227 | Sord | 2.365 | 2.07E-04 | 8.19E-03 |
| ENSMUSG00000034371 | Tkfc | 2.367 | 1.61E-04 | 6.85E-03 |
| ENSMUSG00000020486 | 45173 | 2.373 | 1.42E-04 | 6.22E-03 |
| ENSMUSG00000085614 | 1700123M08 | 2.377 | 8.23E-05 | 4.38E-03 |
| ENSMUSG00000062901 | Klhl24 | 2.377 | 8.57E-09 | 3.77E-06 |
| ENSMUSG00000051515 | Fam181b | 2.397 | 6.80E-05 | 3.78E-03 |
| ENSMUSG00000027875 | Hmgcs2 | 2.403 | 5.50E-04 | 1.64E-02 |
| ENSMUSG00000038248 | Sobp | 2.425 | 5.58E-06 | 5.74E-04 |
| ENSMUSG00000051098 | Mblac2 | 2.428 | 2.41E-03 | 4.65E-02 |
| ENSMUSG00000020962 | Gtf2a1 | 2.441 | 1.24E-04 | 5.63E-03 |
| ENSMUSG00000033488 | Cryzl2 | 2.442 | 2.10E-08 | 8.13E-06 |
| ENSMUSG00000039765 | Cc2d2a | 2.445 | 7.43E-04 | 2.04E-02 |
| ENSMUSG00000020263 | Appl2 | 2.454 | 5.24E-07 | 9.93E-05 |
| ENSMUSG00000061535 | C1qtnf7 | 2.468 | 1.77E-04 | 7.38E-03 |
| ENSMUSG00000020388 | Pdlim4 | 2.506 | 1.36E-07 | 3.36E-05 |
| ENSMUSG00000116656 | AC131339.2 | 2.536 | 2.08E-04 | 8.20E-03 |
| ENSMUSG00000038543 | BC028528 | 2.539 | 3.95E-04 | 1.31E-02 |
| ENSMUSG00000055435 | Maf | 2.543 | 1.18E-03 | 2.84E-02 |
| ENSMUSG00000031284 | Pak3 | 2.558 | 2.22E-03 | 4.38E-02 |
| ENSMUSG00000025875 | Tspan17 | 2.565 | 1.54E-04 | 6.62E-03 |
| ENSMUSG00000028015 | Ctso | 2.571 | 9.22E-04 | 2.41E-02 |
| ENSMUSG00000024598 | Fbn2 | 2.578 | 1.83E-03 | 3.80E-02 |
| ENSMUSG00000030793 | Pycard | 2.587 | 6.26E-05 | 3.58E-03 |
| ENSMUSG00000031310 | Zmym3 | 2.598 | 7.04E-06 | 6.81E-04 |
| ENSMUSG00000099034 | 2810039B14 | 2.604 | 3.04E-04 | 1.08E-02 |
| ENSMUSG00000110935 | Gm8834 | 2.606 | 8.64E-05 | 4.58E-03 |
| ENSMUSG00000001520 | Nrip2 | 2.619 | 1.48E-04 | 6.43E-03 |
| ENSMUSG00000070315 | 4930581F22 | 2.623 | 1.62E-03 | 3.55E-02 |
| ENSMUSG00000002997 | Prkar2b | 2.654 | 1.10E-07 | 2.81E-05 |
| ENSMUSG00000042109 | Csdc2 | 2.658 | 2.20E-05 | 1.62E-03 |
| ENSMUSG00000021884 | Hacl1 | 2.688 | 2.69E-08 | 9.78E-06 |
| ENSMUSG00000028773 | Fabp3 | 2.697 | 1.90E-04 | 7.77E-03 |
| ENSMUSG00000041132 | N4bp2l1 | 2.788 | 1.03E-03 | 2.61E-02 |
| ENSMUSG00000036545 | Adamts2 | 2.790 | 5.94E-05 | 3.48E-03 |
| ENSMUSG00000022440 | C1qtnf6 | 2.791 | 1.97E-06 | 2.80E-04 |
| ENSMUSG00000087380 | 2210408F21 | 2.793 | 2.72E-05 | 1.90E-03 |
| ENSMUSG00000089996 | Tmsb15b2 | 2.795 | 1.52E-04 | 6.54E-03 |
| ENSMUSG00000078350 | Smim1 | 2.808 | 1.16E-04 | 5.47E-03 |
| ENSMUSG00000001313 | Rnd2 | 2.812 | 1.95E-04 | 7.91E-03 |
| ENSMUSG00000116097 | AL590144.2 | 2.820 | 6.79E-05 | 3.78E-03 |
| ENSMUSG00000052957 | Gas1 | 2.838 | 2.65E-03 | 4.96E-02 |
| ENSMUSG00000033420 | Antxr1 | 2.843 | 1.18E-04 | 5.50E-03 |
| ENSMUSG00000022389 | Tef | 2.873 | 5.18E-07 | 9.92E-05 |
| ENSMUSG00000040666 | Sh3bgr | 2.876 | 1.80E-03 | 3.76E-02 |
| ENSMUSG00000023982 | Guca1a | 2.885 | 1.15E-04 | 5.47E-03 |
| ENSMUSG00000002346 | Slc25a42 | 2.896 | 1.88E-03 | 3.87E-02 |
| ENSMUSG00000101655 | 2310040G24 | 2.900 | 6.63E-04 | 1.89E-02 |

| | | | | |
|---|---|---|---|---|
| ENSMUSG00000029838 | Ptn | 2.902 | 2.56E-03 | 4.85E-02 |
| ENSMUSG00000085260 | Med9os | 2.925 | 6.14E-04 | 1.78E-02 |
| ENSMUSG00000072966 | Gprasp2 | 2.949 | 1.33E-03 | 3.10E-02 |
| ENSMUSG00000033208 | S100b | 2.961 | 1.82E-03 | 3.80E-02 |
| ENSMUSG00000030074 | Gxylt2 | 2.985 | 9.82E-04 | 2.51E-02 |
| ENSMUSG00000039533 | Mmd2 | 2.986 | 7.15E-04 | 2.00E-02 |
| ENSMUSG00000057948 | Unc13d | 2.988 | 4.55E-04 | 1.43E-02 |
| ENSMUSG00000064342 | mt-Ti | 2.993 | 1.64E-03 | 3.57E-02 |
| ENSMUSG00000112646 | Gm5654 | 3.024 | 2.88E-04 | 1.04E-02 |
| ENSMUSG00000059022 | Kcp | 3.026 | 1.64E-04 | 6.93E-03 |
| ENSMUSG00000013083 | 2200002J24I | 3.029 | 1.65E-03 | 3.58E-02 |
| ENSMUSG00000060224 | Pyroxd2 | 3.043 | 1.48E-03 | 3.32E-02 |
| ENSMUSG00000055692 | Tmem191c | 3.070 | 1.14E-04 | 5.45E-03 |
| ENSMUSG00000050953 | Gja1 | 3.081 | 3.72E-04 | 1.25E-02 |
| ENSMUSG00000086382 | Chrna1os | 3.095 | 7.48E-08 | 2.04E-05 |
| ENSMUSG00000086900 | Kcnab3os | 3.103 | 4.26E-04 | 1.37E-02 |
| ENSMUSG00000026489 | Coq8a | 3.114 | 2.39E-09 | 1.25E-06 |
| ENSMUSG00000041476 | Smpx | 3.115 | 2.25E-08 | 8.36E-06 |
| ENSMUSG00000020099 | Unc5b | 3.167 | 2.01E-04 | 8.04E-03 |
| ENSMUSG00000086863 | Gm15543 | 3.194 | 4.36E-04 | 1.38E-02 |
| ENSMUSG00000022883 | Robo1 | 3.217 | 8.92E-04 | 2.35E-02 |
| ENSMUSG00000042500 | Ago4 | 3.229 | 1.23E-04 | 5.62E-03 |
| ENSMUSG00000074595 | Wfdc6a | 3.237 | 7.31E-04 | 2.03E-02 |
| ENSMUSG00000085347 | Hk1os | 3.268 | 1.22E-05 | 1.00E-03 |
| ENSMUSG00000055493 | Epm2a | 3.272 | 2.14E-03 | 4.24E-02 |
| ENSMUSG00000027966 | Col11a1 | 3.297 | 8.14E-07 | 1.39E-04 |
| ENSMUSG00000024818 | Slc25a45 | 3.319 | 7.23E-04 | 2.01E-02 |
| ENSMUSG00000027217 | Tspan18 | 3.326 | 2.01E-04 | 8.04E-03 |
| ENSMUSG00000038776 | Ephx1 | 3.338 | 1.44E-03 | 3.28E-02 |
| ENSMUSG00000012819 | Cdh23 | 3.352 | 1.39E-03 | 3.20E-02 |
| ENSMUSG00000045731 | Pnoc | 3.359 | 2.18E-06 | 3.01E-04 |
| ENSMUSG00000097354 | 2310001H17 | 3.388 | 5.53E-04 | 1.65E-02 |
| ENSMUSG00000041536 | Serpina3a | 3.401 | 3.59E-04 | 1.22E-02 |
| ENSMUSG00000033610 | Pank1 | 3.405 | 2.77E-06 | 3.64E-04 |
| ENSMUSG00000031877 | Ces2g | 3.412 | 1.78E-03 | 3.74E-02 |
| ENSMUSG00000090925 | 1810064F22 | 3.426 | 1.85E-03 | 3.84E-02 |
| ENSMUSG00000081359 | Cox20-ps | 3.426 | 1.06E-03 | 2.65E-02 |
| ENSMUSG00000034810 | Scn7a | 3.432 | 1.47E-03 | 3.31E-02 |
| ENSMUSG00000097413 | A830052D11 | 3.444 | 1.47E-03 | 3.31E-02 |
| ENSMUSG00000022235 | Cmbl | 3.450 | 9.50E-09 | 4.04E-06 |
| ENSMUSG00000032224 | Fam81a | 3.458 | 5.07E-07 | 9.80E-05 |
| ENSMUSG00000029163 | Emilin1 | 3.469 | 4.97E-04 | 1.53E-02 |
| ENSMUSG00000100005 | B130024G19 | 3.475 | 4.36E-04 | 1.38E-02 |
| ENSMUSG00000035561 | Aldh1b1 | 3.478 | 1.75E-03 | 3.68E-02 |
| ENSMUSG00000041000 | Trim62 | 3.481 | 1.90E-09 | 1.08E-06 |
| ENSMUSG00000033715 | Akr1c14 | 3.483 | 4.08E-11 | 3.94E-08 |
| ENSMUSG00000031871 | Cdh5 | 3.488 | 1.25E-04 | 5.63E-03 |

| | | | | |
|---|---|---|---|---|
| ENSMUSG00000067279 | Ppp1r3c | 3.507 | 2.12E-03 | 4.22E-02 |
| ENSMUSG00000107736 | Gm44148 | 3.509 | 6.04E-04 | 1.77E-02 |
| ENSMUSG00000050645 | Defb19 | 3.517 | 2.31E-04 | 8.82E-03 |
| ENSMUSG00000025934 | Gsta3 | 3.559 | 7.80E-11 | 6.56E-08 |
| ENSMUSG00000040111 | Gramd1b | 3.601 | 1.73E-05 | 1.35E-03 |
| ENSMUSG00000021185 | Dglucy | 3.614 | 5.47E-05 | 3.27E-03 |
| ENSMUSG00000026831 | 1700007K13 | 3.615 | 1.73E-03 | 3.67E-02 |
| ENSMUSG00000079013 | Serpina3j | 3.659 | 1.66E-04 | 7.01E-03 |
| ENSMUSG00000044835 | Ankrd45 | 3.670 | 4.07E-04 | 1.33E-02 |
| ENSMUSG00000025468 | Caly | 3.683 | 3.92E-04 | 1.30E-02 |
| ENSMUSG00000030474 | Siglece | 3.684 | 2.57E-03 | 4.86E-02 |
| ENSMUSG00000059857 | Ntng1 | 3.689 | 2.41E-03 | 4.66E-02 |
| ENSMUSG00000017453 | Pipox | 3.706 | 1.63E-03 | 3.56E-02 |
| ENSMUSG00000045672 | Col27a1 | 3.754 | 1.66E-03 | 3.58E-02 |
| ENSMUSG00000027489 | Necab3 | 3.756 | 7.42E-04 | 2.04E-02 |
| ENSMUSG00000019301 | Hsd17b1 | 3.761 | 6.43E-05 | 3.65E-03 |
| ENSMUSG00000004892 | Bcan | 3.799 | 5.00E-04 | 1.53E-02 |
| ENSMUSG00000030257 | Srgap3 | 3.806 | 4.62E-05 | 2.84E-03 |
| ENSMUSG00000046182 | Gsg1l | 3.817 | 1.28E-03 | 3.01E-02 |
| ENSMUSG00000042129 | Rassf4 | 3.825 | 2.67E-04 | 9.93E-03 |
| ENSMUSG00000001663 | Gstt1 | 3.833 | 2.12E-09 | 1.17E-06 |
| ENSMUSG00000021294 | Kif26a | 3.834 | 1.29E-03 | 3.03E-02 |
| ENSMUSG00000037161 | Mgarp | 3.847 | 1.96E-04 | 7.93E-03 |
| ENSMUSG00000068854 | Hist2h2be | 3.849 | 1.21E-03 | 2.90E-02 |
| ENSMUSG00000034059 | Ypel4 | 3.849 | 9.56E-04 | 2.47E-02 |
| ENSMUSG00000109080 | Gm38944 | 3.939 | 1.22E-03 | 2.91E-02 |
| ENSMUSG00000039676 | Capsl | 3.946 | 7.07E-04 | 1.99E-02 |
| ENSMUSG00000109992 | Gm45650 | 3.954 | 1.66E-03 | 3.58E-02 |
| ENSMUSG00000074665 | Bpifb4 | 3.961 | 6.16E-04 | 1.79E-02 |
| ENSMUSG00000033082 | Clec1a | 3.965 | 2.40E-03 | 4.64E-02 |
| ENSMUSG00000021806 | Nid2 | 3.973 | 2.01E-05 | 1.50E-03 |
| ENSMUSG00000001665 | Gstt3 | 3.993 | 3.76E-08 | 1.25E-05 |
| ENSMUSG00000028415 | Spink4 | 3.994 | 7.49E-04 | 2.05E-02 |
| ENSMUSG00000024620 | Pdgfrb | 4.013 | 1.46E-04 | 6.35E-03 |
| ENSMUSG00000045392 | Olfr1033 | 4.015 | 1.47E-03 | 3.31E-02 |
| ENSMUSG00000034413 | Neurl1b | 4.022 | 8.53E-04 | 2.26E-02 |
| ENSMUSG00000053821 | Gm9922 | 4.027 | 1.71E-03 | 3.64E-02 |
| ENSMUSG00000021213 | Akr1c13 | 4.028 | 1.16E-04 | 5.48E-03 |
| ENSMUSG00000032135 | Mcam | 4.049 | 7.83E-04 | 2.12E-02 |
| ENSMUSG00000047363 | Cstad | 4.062 | 5.42E-04 | 1.63E-02 |
| ENSMUSG00000025094 | Slc18a2 | 4.066 | 2.28E-03 | 4.47E-02 |
| ENSMUSG00000037754 | Ppp1r16b | 4.076 | 2.70E-05 | 1.90E-03 |
| ENSMUSG00000054675 | Tmem119 | 4.082 | 2.65E-03 | 4.96E-02 |
| ENSMUSG00000027004 | Frzb | 4.112 | 1.61E-03 | 3.53E-02 |
| ENSMUSG00000101378 | Gm28086 | 4.132 | 2.77E-04 | 1.01E-02 |
| ENSMUSG00000020467 | Efemp1 | 4.145 | 1.90E-04 | 7.77E-03 |
| ENSMUSG00000027347 | Rasgrp1 | 4.166 | 1.43E-03 | 3.26E-02 |

| | | | | |
|---|---|---|---|---|
| ENSMUSG00000037129 | Tmprss13 | 4.250 | 1.02E-04 | 5.08E-03 |
| ENSMUSG00000102979 | Gm37350 | 4.255 | 6.81E-05 | 3.78E-03 |
| ENSMUSG00000038453 | Srcin1 | 4.274 | 8.73E-05 | 4.60E-03 |
| ENSMUSG00000013653 | 1810065E05 | 4.299 | 2.43E-03 | 4.67E-02 |
| ENSMUSG00000030935 | Acsm3 | 4.302 | 1.11E-03 | 2.74E-02 |
| ENSMUSG00000075410 | Prcd | 4.306 | 5.66E-07 | 1.04E-04 |
| ENSMUSG00000041329 | Atp1b2 | 4.309 | 3.44E-04 | 1.19E-02 |
| ENSMUSG00000112220 | Gm36208 | 4.316 | 1.32E-03 | 3.08E-02 |
| ENSMUSG00000029650 | Slc46a3 | 4.326 | 9.60E-05 | 4.84E-03 |
| ENSMUSG00000001827 | Folr1 | 4.330 | 3.58E-04 | 1.22E-02 |
| ENSMUSG00000042249 | Grk3 | 4.330 | 1.81E-03 | 3.78E-02 |
| ENSMUSG00000074218 | Cox7a1 | 4.333 | 8.36E-10 | 5.39E-07 |
| ENSMUSG00000074678 | Defb25 | 4.335 | 1.57E-03 | 3.48E-02 |
| ENSMUSG00000074594 | Wfdc9 | 4.356 | 5.98E-04 | 1.76E-02 |
| ENSMUSG00000025176 | Hoga1 | 4.386 | 1.14E-03 | 2.77E-02 |
| ENSMUSG00000084915 | C230037L18 | 4.388 | 2.55E-03 | 4.83E-02 |
| ENSMUSG00000085354 | Gm2044 | 4.406 | 1.41E-04 | 6.22E-03 |
| ENSMUSG00000023885 | Thbs2 | 4.413 | 1.61E-04 | 6.85E-03 |
| ENSMUSG00000089671 | Gm16537 | 4.424 | 3.65E-04 | 1.23E-02 |
| ENSMUSG00000024747 | Aldh1a7 | 4.456 | 4.84E-08 | 1.42E-05 |
| ENSMUSG00000026241 | Nppc | 4.509 | 2.14E-03 | 4.25E-02 |
| ENSMUSG00000037206 | Islr | 4.536 | 1.39E-04 | 6.13E-03 |
| ENSMUSG00000040569 | Slc26a7 | 4.538 | 1.65E-03 | 3.58E-02 |
| ENSMUSG00000073821 | 8030451A03 | 4.555 | 4.47E-05 | 2.77E-03 |
| ENSMUSG00000110489 | Gm31659 | 4.576 | 3.16E-04 | 1.11E-02 |
| ENSMUSG00000102923 | Gm31728 | 4.602 | 1.32E-03 | 3.08E-02 |
| ENSMUSG00000107689 | Gm44386 | 4.609 | 1.82E-11 | 1.85E-08 |
| ENSMUSG00000083222 | Gm12427 | 4.620 | 1.50E-03 | 3.36E-02 |
| ENSMUSG00000074207 | Adh1 | 4.653 | 8.90E-05 | 4.64E-03 |
| ENSMUSG00000031789 | Cngb1 | 4.655 | 3.08E-06 | 3.90E-04 |
| ENSMUSG00000046807 | Lrrc75b | 4.701 | 1.92E-03 | 3.91E-02 |
| ENSMUSG00000070304 | Scn2b | 4.709 | 2.57E-05 | 1.84E-03 |
| ENSMUSG00000074604 | Mgst2 | 4.757 | 8.09E-06 | 7.59E-04 |
| ENSMUSG00000056174 | Col8a2 | 4.782 | 5.27E-04 | 1.59E-02 |
| ENSMUSG00000065952 | C330021F23 | 4.842 | 8.23E-04 | 2.20E-02 |
| ENSMUSG00000020096 | Tbata | 4.846 | 2.14E-05 | 1.58E-03 |
| ENSMUSG00000087612 | A230005M16 | 4.852 | 7.89E-05 | 4.21E-03 |
| ENSMUSG00000035407 | Kank4 | 4.854 | 2.68E-07 | 5.57E-05 |
| ENSMUSG00000086909 | Gm4926 | 4.868 | 9.03E-05 | 4.65E-03 |
| ENSMUSG00000072941 | Sod3 | 4.889 | 9.98E-06 | 8.65E-04 |
| ENSMUSG00000101438 | Gm19412 | 4.902 | 2.03E-03 | 4.09E-02 |
| ENSMUSG00000037709 | Fam13a | 4.930 | 6.16E-06 | 6.17E-04 |
| ENSMUSG00000048038 | Ccdc187 | 4.950 | 2.89E-04 | 1.04E-02 |
| ENSMUSG00000021416 | Eci3 | 4.979 | 1.44E-06 | 2.19E-04 |
| ENSMUSG00000027870 | Hao2 | 5.082 | 4.61E-06 | 4.88E-04 |
| ENSMUSG00000043664 | Tmem221 | 5.101 | 3.97E-04 | 1.31E-02 |
| ENSMUSG00000100022 | Gm29590 | 5.150 | 9.24E-04 | 2.41E-02 |

| | | | | |
|---|---|---|---|---|
| ENSMUSG00000031872 | Bean1 | 5.198 | 5.99E-06 | 6.06E-04 |
| ENSMUSG00000066361 | Serpina3c | 5.198 | 1.27E-03 | 3.01E-02 |
| ENSMUSG00000093479 | Gm20629 | 5.207 | 1.59E-03 | 3.51E-02 |
| ENSMUSG00000024245 | Tmem178 | 5.245 | 3.80E-06 | 4.24E-04 |
| ENSMUSG00000055430 | Nap1l5 | 5.262 | 8.63E-06 | 7.95E-04 |
| ENSMUSG00000092323 | BB365896 | 5.302 | 3.75E-04 | 1.26E-02 |
| ENSMUSG00000029343 | Crybb1 | 5.305 | 1.21E-03 | 2.90E-02 |
| ENSMUSG00000027871 | Hsd3b1 | 5.315 | 5.54E-16 | 1.78E-12 |
| ENSMUSG00000023153 | Tmem52 | 5.345 | 7.32E-04 | 2.03E-02 |
| ENSMUSG00000097717 | 1700066J03I | 5.470 | 6.47E-04 | 1.86E-02 |
| ENSMUSG00000086813 | Gm13657 | 5.551 | 1.01E-04 | 5.04E-03 |
| ENSMUSG00000113965 | A730091E23 | 5.577 | 6.59E-04 | 1.89E-02 |
| ENSMUSG00000044062 | Plekhd1os | 5.577 | 6.82E-04 | 1.94E-02 |
| ENSMUSG00000113598 | Gm48287 | 5.583 | 4.39E-05 | 2.74E-03 |
| ENSMUSG00000069385 | Gm10267 | 5.617 | 2.27E-03 | 4.45E-02 |
| ENSMUSG00000020475 | Pgam2 | 5.640 | 1.96E-04 | 7.93E-03 |
| ENSMUSG00000112736 | Gm48718 | 5.690 | 9.33E-05 | 4.75E-03 |
| ENSMUSG00000020562 | Efcab10 | 5.702 | 1.06E-04 | 5.19E-03 |
| ENSMUSG00000063594 | Gng8 | 5.747 | 1.52E-03 | 3.38E-02 |
| ENSMUSG00000001985 | Grik3 | 5.780 | 3.30E-06 | 3.97E-04 |
| ENSMUSG00000116573 | AC161816.1 | 5.782 | 7.31E-04 | 2.03E-02 |
| ENSMUSG00000084942 | 7330404K18 | 5.827 | 9.01E-05 | 4.65E-03 |
| ENSMUSG00000047786 | Lix1 | 5.872 | 9.89E-06 | 8.61E-04 |
| ENSMUSG00000034488 | Edil3 | 5.962 | 8.81E-05 | 4.61E-03 |
| ENSMUSG00000104020 | Gm37215 | 6.082 | 9.44E-04 | 2.45E-02 |
| ENSMUSG00000085876 | Gm12409 | 6.095 | 4.07E-04 | 1.33E-02 |
| ENSMUSG00000041193 | Pla2g5 | 6.134 | 1.00E-04 | 5.01E-03 |
| ENSMUSG00000077058 | Gm24542 | 6.153 | 2.45E-05 | 1.78E-03 |
| ENSMUSG00000074596 | Spint3 | 6.166 | 7.16E-05 | 3.95E-03 |
| ENSMUSG00000069814 | Ccdc92b | 6.265 | 5.51E-05 | 3.29E-03 |
| ENSMUSG00000107876 | Gm43936 | 6.266 | 6.30E-06 | 6.22E-04 |
| ENSMUSG00000027584 | Oprl1 | 6.339 | 4.83E-04 | 1.49E-02 |
| ENSMUSG00000038354 | Ankrd35 | 6.402 | 1.07E-06 | 1.71E-04 |
| ENSMUSG00000085419 | Gm11734 | 6.538 | 8.23E-07 | 1.40E-04 |
| ENSMUSG00000085121 | 5730437C11 | 6.584 | 9.14E-05 | 4.69E-03 |
| ENSMUSG00000104420 | E230020A03 | 6.674 | 2.50E-04 | 9.45E-03 |
| ENSMUSG00000113974 | 4930442G10 | 6.811 | 3.43E-05 | 2.26E-03 |
| ENSMUSG00000030249 | Abcc9 | 6.819 | 3.04E-06 | 3.86E-04 |
| ENSMUSG00000113744 | Gm47925 | 7.216 | 3.57E-06 | 4.13E-04 |
| ENSMUSG00000040026 | Saa3 | 7.399 | 2.22E-05 | 1.62E-03 |
| ENSMUSG00000104109 | Gm30292 | 7.424 | 3.60E-06 | 4.13E-04 |
| ENSMUSG00000021613 | Hapln1 | 7.526 | 7.49E-05 | 4.09E-03 |
| ENSMUSG00000075010 | AW112010 | 8.355 | 5.20E-04 | 1.58E-02 |
| ENSMUSG00000041550 | Serpina5 | 8.604 | 9.53E-06 | 8.49E-04 |
| ENSMUSG00000002324 | Rec8 | 8.619 | 2.02E-06 | 2.83E-04 |
| ENSMUSG00000041445 | Mmrn2 | 19.080 | 5.46E-05 | 3.27E-03 |
| ENSMUSG00000020928 | Higd1b | 19.494 | 9.35E-04 | 2.43E-02 |
